# In Vitro and Viral Evolution Convergence Reveal the Selective Pressures Driving Omicron Emergence

**DOI:** 10.1101/2025.04.23.650148

**Authors:** Aviv Shoshany, Ruojin Tian, Miguel Padilla-Blanco, Adam Hruška, Aditi Konar, Katarina Baxova, Eyal Zoler, Martin Mokrejš, Gideon Schreiber, Jiří Zahradník

## Abstract

In vitro protein evolution provides powerful insights into the amino acid sequences that underlie key biological functions. Here, we used this approach to explore the evolutionary trajectories of the SARS-CoV-2 spike protein receptor-binding domain (RBD) constrained to engage the human ACE2 receptor—an essential first step in viral infection. Applying mild (LSS) or stringent (HSS) selection pressures starting from the ancestral Wuhan strain, we found that HSS, but not LSS rapidly converged on mutations characteristic of the Omicron variant. HSS resulted in fewer, but dominant, non-synonymous mutations mirroring Omicron mutations and its advanced sub-lineages. Conversely, LSS produced only some Omicron-like mutations at much lower frequencies and with incomplete representation. Notably, initiating evolution from Omicron itself resulted in high-fidelity maintenance of Omicron-defining mutations under both HSS and LSS conditions. This evolutionary pattern parallels global SARS-CoV-2 mutation trends as well as in silico simulations, emphasizing the critical role of receptor-binding constraints in shaping viral adaptation, which may be a frequent driver during zoonosis. Predominantly immune evasion associated mutations not selected in vitro. Our findings demonstrate the predictive capacity of in vitro evolution, suggesting Omicron’s abrupt emergence resulted from rare, high-stringency selection, superimposed on a background of broader, milder pressures, with Omicron being the humanized SARS-CoV-2.

## Introduction

The emergence of SARS-CoV-2 in late 2019 initiated a global pandemic with profound impacts on public health. The initial Wuhan strain was rapidly replaced by successive viral lineages with higher fitness. Viral mutations are accumulated and selected during the virus life cycle, with most mutations being lost due to strong selection during the tight bottlenecks imposed ^1,2^. Transmissibility, usually expressed as the net reproduction number (*R*t), is assumed to closely approximate their fitness at the host population level ^3^. The mutation rate of SARS-CoV-2 is estimated to be 10^−6^ mutations per nucleotide per replication cycle, similar to other beta coronaviruses ^4^. In addition, frequent C → U mutations associated with host APOBEC enzyme account in part, for the strikingly high ratio of non-synonymous changes in SARS-CoV-2 genomes compared with those at synonymous sites ^5,6^. Due to the randomness of mutation events, single-nucleotide mutations dominate the early phases of the pandemic, which allows for substitution to 7-10 other amino-acids, as indeed observed for SARS-CoV-2. Double nucleotide mutations and epistatic mutations emerge gradually later ^7^.

Central to SARS-CoV-2 transmission and pathogenesis is the Spike (S) protein, which underwent rapid adaptive evolution in the human population, shaping the trajectory of the pandemic ^8-10^. The S protein consists of two subunits: S1, which contains the receptor binding domain (RBD) and S2, responsible for membrane fusion ^11^. The RBD directly interacts with the ACE2 receptor of the host cell, making it critical for viral entry and a primary target for neutralizing antibodies ^12^. The RBD has become a focal point of research as its structure and function are intricately tied to viral fitness, transmissibility, and immune escape. Molecular studies have revealed that even subtle modifications in the RBD, such as found in the Alpha, Beta, Gamma and Delta variants, can significantly alter viral transmission efficiency and host cell tropism ^13,14^. Its adaptive evolution underlies the emergence of high-fitness variants, most of which have been designated by the WHO as Variants of Concern (VOCs) or variants of interest (VOIs) and have been closely monitored ^15^. Omicron, which evolved at the end of 2021, was the most radical alteration of the RBD, and since then all VOCs are of the Omicron lineage. The emergence of the Omicron variant marked a pivotal shift in the pandemic’s trajectory. Omicron displayed an unprecedented number of over 30 mutations in the Spike protein alone (15 of them in the RBD) ^12,16^, which altered its interaction with ACE2 and neutralizing antibodies ^12,17^. These mutations enhanced its ability to evade pre-existing immunity from vaccination and prior infections, contributing to widespread breakthrough infections. Omicron also demonstrated increased transmissibility, which, alongside immune escape, raised questions about its origins. Theories regarding Omicron’s evolution include prolonged infection in an immunocompromised host, facilitating the accumulation of mutations, or cross-species transmission, followed by adaptive evolution in an animal reservoir before re-entering the human population ^3^. The mutations in Omicron altered the virus’s behavior, including a shift from TMPRSS2-dependent entry to preferential entry via the endosomal pathway ^11,18^. This change potentially modified the virus’s tissue tropism, favoring the upper respiratory tract over the lower respiratory tract, which may have contributed to increased transmissibility but decreased pathogenicity ^10^. Despite significant progress in characterizing Omicron’s genomic and phenotypic properties, the precise evolutionary pathway remains an enigma.

Selective pressures on the SARS-CoV-2 RBD are multifactorial, involving a balance between ACE2 binding affinity, immune evasion, and structural stability. Studies have highlighted the role of ACE2 affinity in viral fitness, with variants exhibiting increased ACE2 binding often demonstrating enhanced transmissibility ^14,19^. This relationship is modulated by compensatory mutations that balance the competing demands of immune evasion and structural integrity. For example, the Omicron variant harbors key mutations in the ACE2 binding region—such as K417N, S477N, E484A, Q493R, G496S, Q498R, N501Y, and Y505H—that enhance ACE2 binding and/or immune evasion. Other mutations like G339D, N440K, S373P, S375F, and T478K, while not directly interacting with ACE2, are frequently found in Omicron variants. Multiple studies showed the trend of increasing binding throughout the SARS-CoV-2 evolution ^19,20^. In addition, deep mutational scanning studies have shown that mutations at protein-protein interfaces rarely have neutral effects and often influence binding properties, demonstrating the complexity of immune escape mechanisms ^21-23^. While numerous studies have explored RBD evolution in the context of immune escape, the precise interplay of selective pressures shaping these mutations remains incompletely understood, highlighting the need for alternative approaches to deconvolute these forces.

In a previous study, affinity maturation of the SARS-CoV-2 RBD towards ACE2 resulted in a mutant RBD with 600-fold increase in binding affinity. This work demonstrated the rapid selection of affinity-enhancing mutations observed in viral population ^24^. Notably, we quickly acquired the mutations N460K, S477N, E484K, Q498R, and N501Y, which form part of the Omicron BA.1 variant ^12^. However, that study had certain limitations as its primary objective was developing an infection inhibitor ^25^ rather than studying evolutionary processes. First, the codons were optimized for expression in *Saccharomyces cerevisiae* for yeast display, meaning that single-point mutations could result in different amino acids than those derived from the original sequence. Second, the in vitro evolution included a pre-equilibrium selection step favoring faster on-rates, which may not align with natural viral evolution. Therefore, we decided to conduct a study that would lay a robust foundation for using in vitro evolution to predict viral evolution. Accordingly, in this study, we provide evidence that the key mutations defining Omicron lineage RBDs can be explained primarily by adaptation to human ACE2 receptor ortholog and stability maturation, but only under stringent selection pressures. Moreover, our analysis of low-stringency selection, combined with fitness/reward modeling, elucidates how mutations with differing effects on fitness behave under varying selective pressures. As a large proportion of modern epidemics are caused by zoonosis and adaptation to the orthologous receptor, these insights lay the groundwork for more robust predictions of viral variant emergence and evolutionary trajectories, showing a limited number of tipping points in the adaptive trajectory.

## Results and discussion

### Experimental design and setup for parallel in vitro evolution

In this study we evaluated RBD evolution driven by ACE2 binding and protein-stability, starting from multiple lineages and using different regimes of evolutionary pressures. Our goal was to determine how different selective pressures for affinity and stability shape mutational outcomes excluding immune pressures, whether convergent or divergent adaptive pathways emerge and to what extent the mutational paths will mirror SARS-CoV-2 viral evolution. First, we established a high-throughput system capable of accurately identifying the maximum number of clones from individual libraries throughout multiple rounds of in vitro evolution. For this, we extended our original strategy, which employed an enhanced yeast surface-display protocol incorporating two distinct detection strategies—eUnaG2 and DnbALFA ^26^. We introduced the SpyTag–SpyCatcher003 system (pJYDC4) ^27^, designed stability and expression enhanced monomeric avidin based on rhizavidin (DeMA; Supporting information part PS1, pJYDC6 ^28^), made an alternative plasmid encoding for chloramphenicol resistance and incorporated barcodes at both the N- and C-gene termini ensuring library stability throughout in vitro evolution. The N-terminal barcode was added as an extension of the existing GS linker within the expression construct. To prevent any potential interference with yeast display performance, we tested the barcodes using the expression of 3EFR-Cfr-Anti-Streptavidin ^26^ and its binding to Streptavidin-APC (Table S1). Overall, our system consists of ten barcodes, four labeling strategies, and two selection antibiotics (Fig. 1A), representing a scalable and reliable framework for parallel in vitro evolutionary studies with minimal risk of cross-contamination or library loss due to unwanted impurities.

**Fig. 1.**
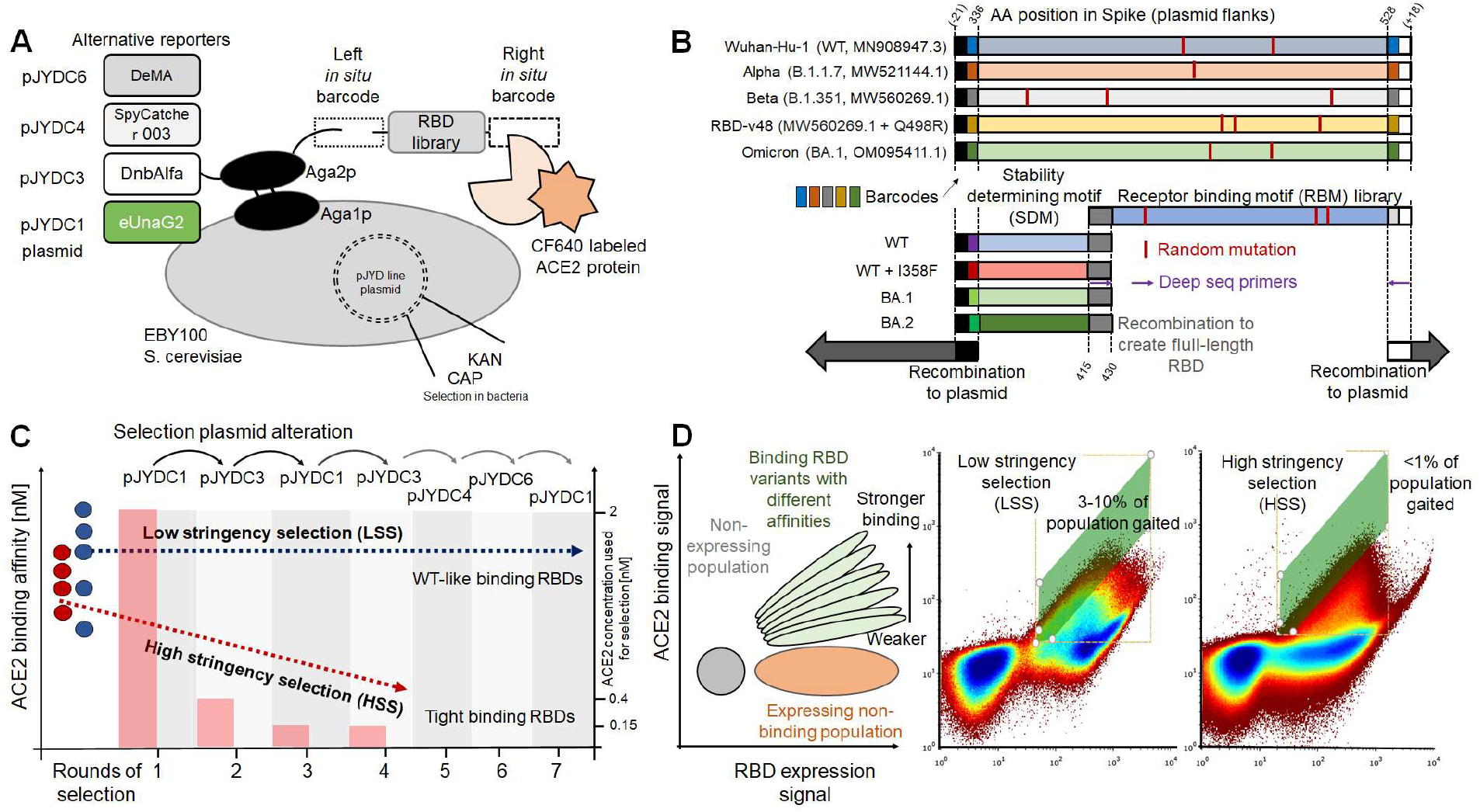
Setup for high-throughput yeast display for in vitro evolution. (A) General setup of the yeast display system including alternative reporter proteins, in situ barcodes, and additional plasmid antibiotics for precise identification of selected clones. (B) Schematic representation of RBD constructs used in vitro evolution libraries tailored for LSS and HSS regimes. (C) Framework for HSS and LSS selection cycles, highlighting the use of distinct plasmid alterations to mitigate the risk of cross-contamination between libraries and respective evolutionary pathways. Constant or decreasing concentrations of ACE2 were used for HSS and LSS, respectively. (D) Scheme (left panel) and FACS density dot plots showing using different gating strategies for selection (middle LSS and right panel HSS, for details see Fig. S1).

### Exploring mutational trajectories under varying selective pressures

We adopted a strategy with a high degree of redundancy, characterized by starting with RBD sequences of different lineages and two distinct sorting strategies (Fig. 1B, 1C). The first strategy represents a low stringency selection (LSS), designed to mimic conditions where selective pressure maintains the initial binding affinity. This is achieved using Fluorescence-activated Cell Sorting (FACS) by selecting for 3–10% of the RBD expressing population and a bait (ACE2) concentration of 2 nM ACE2 (which is ∼ EC25 of WT). Each library underwent 7 cycles of random mutagenesis, with each cycle including two rounds of FACS sorting against ACE2 (Fig. 1D, Fig. S1). These permissive conditions allow mutations that slightly reduce binding affinity and protein stability to persist within the population. This strategy was applied in parallel to five different lineage-derived RBD sequences using error-prone mutagenesis covering the full RBD region (Fig. 1B). The starting RBDs originated from SARS-CoV-2 variants: Wuhan-Hu-1 (WT), Alpha, Beta, Omicron BA.1, and RBD variant 48 (RBD-v48, Beta + Q498R mutation (Table S2) ^24^).

The second strategy employed high-stringency selection (HSS), imposing stringent requirements to enhance binding affinity and maintain thermal-stability. Each cycle included random mutagenesis, heating of the yeast cells to 40 °C for 20 minutes (removing less stable mutants) ^29,30^ and three rounds of FACS selection for the top 1% of ACE2-binding by clones. The first cycle of randomly mutated RBDs were selected against 2 nM of ACE2, followed by 0.4 nM ACE2, and finally 0.15 nM ACE2. In addition to WT and WT+I358F (a mutation that increases thermal-stability by 4°C, allowing to explore additional evolutionary pathways ^24^), the RBDs of BA.1 and BA.2 lineages were used as starting sequences for in vitro evolution. To increase the likelihood of isolating high-affinity clones and comprehensively covering potential variants, error-prone mutagenesis was restricted to the receptor-binding motif (RBM, AA431–528, Fig. 1C). Residues 337-430, which do not participate in ACE2 binding, were not mutated (Fig. S2).

Error-prone mutagenesis was experimentally optimized to predominantly generate sequences with a single mutation per gene. Fig. 2 shows an in-depth analysis of the WT library used for HSS. The libraries were analyzed by next generation sequencing (NGS-Illumina paired-end deep sequencing 2 × 250 bp). A two-level barcoding system was employed to process the NGS data, incorporating additional barcodes into amplicons that already contained in situ barcodes embedded within the in vitro evolution plasmids (Fig. 1B and Table S1 and S3). This dual barcoding strategy enabled robust and unambiguous identification of clones originating from distinct libraries and also sorting rounds, ensuring high fidelity in tracking evolutionary trajectories. Fig. 2A shows the sequence Logo plot as determined from NGS, displaying mutation frequencies prior to any selection. Fig. 2B is an analysis of single-nucleotide substitution coverage in the random library prior to selection at specific positions, showing the presence of all theoretically expected mutations across nearly the entire gene length. Fig. 2C shows the sum of frequencies for all single- and double-nucleotide substitutions in the starting library, indicating a homogeneous distribution of frequencies with minimal positional deviations. The frequency of double-nucleotide substitutions is 1/100 relative to single nucleotide substitutions, showing how rare double-nucleotide mutations are, reflecting known observations from viral evolution ^3,7,12^. Fig. 2D sums up the single-nucleotide mutation frequencies per mutant nucleotide and position, showing an equal distribution. Overall, Fig. 2A-D shows the high quality of the starting library before selection. Fig. 2E provides a snapshot of the number of mutations per sequence (similarity score relative to WT) before selection and after two and four rounds of in vitro evolution. While single-nucleotide mutations dominate the starting library, in vitro evolution results in higher numbers of mutations per sequence (with the highest number after the 4^th^ library, Fig. 2E). A complete analysis of the mutant libraries for WT+I358F, BA.1 and BA.2 following the same scheme as shown for WT in Fig. 2A-E is given in Fig. S3-S6 (with WT as reference). The high frequency mutations seen in Figs S4 and S5 are from Omicron relative two WT. Fig. 2F shows the results of Sanger sequencing of the LSS Omicron BA.1 library, showing a gradual increase in the number of mutations in individual sequences upon increasing rounds of in vitro evolution, from 1 to 7 (details for all libraries Fig. S7-S13).

**Fig. 2.**
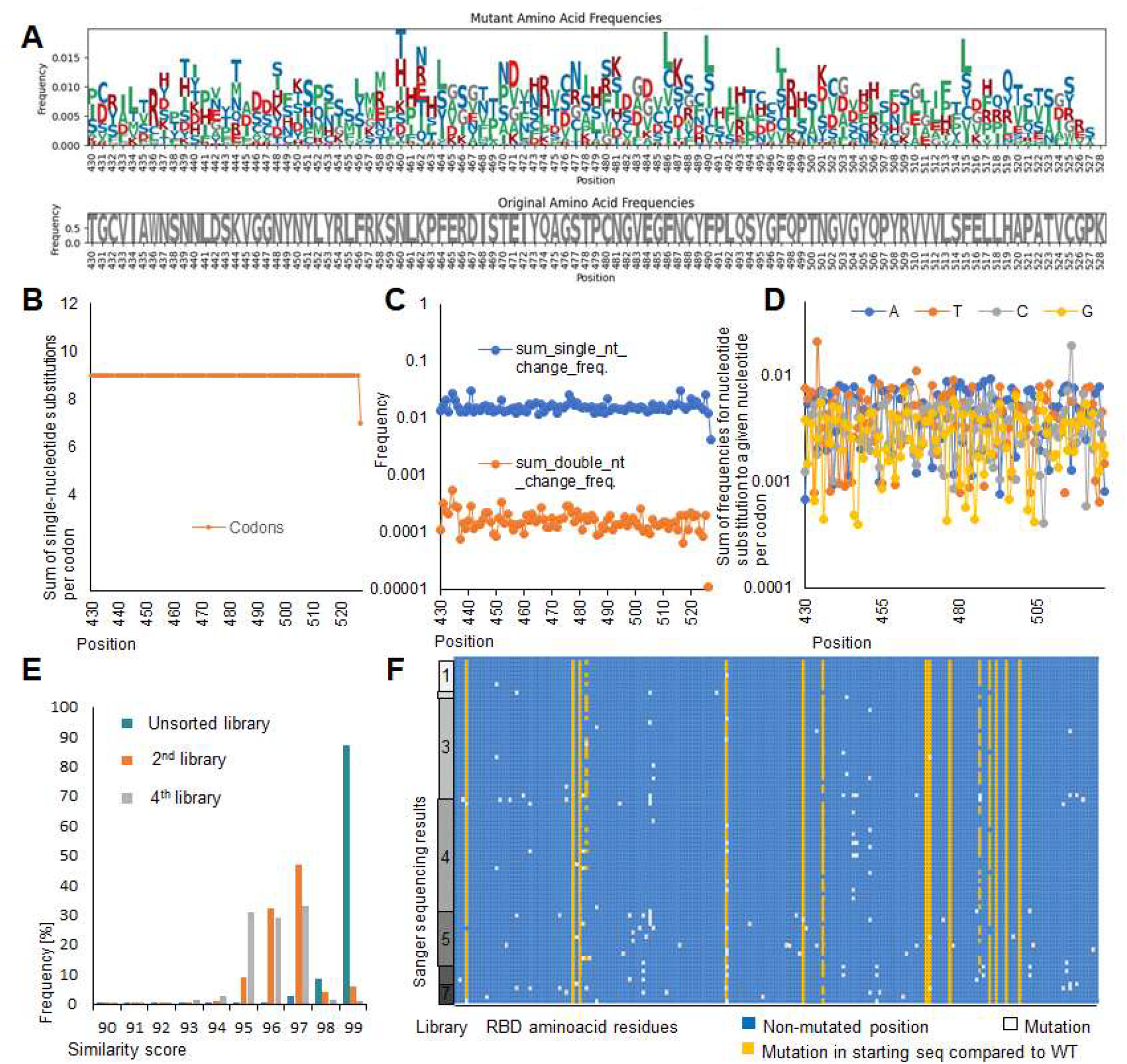
WT library quality assessment. (A) Sequence Logo plot displaying amino-acid frequencies in the HSS WT RBD first library, prior to selections. (B) The number of observed single nucleotide substitutions per codon along the RBM, with 9 representing all mutations in all three positions. (C) Frequencies for all single- and double-nucleotide substitutions per codon, showing minimal positional deviations. (D) Sum of single-nucleotide mutation frequencies per mutant nucleotide and codon. (E) Comparison of sequence similarity scores of individual merged paired-end reads into a single sequence with their respective reference (WT RBD sequence for the unsorted first library) showing increasing number of mutations with progression of in vitro evolution. (F) Sanger sequencing of LSS BA.1 libraries following in vitro evolution, showing a gradual increase in the number of mutations. Similar analyses for LSS and HSS WT+I358F, BA.1 and BA.2 libraries are provided in Supporting information Fig. S3-S13.

### Selection stringency dictates the outcome of in vitro evolution

Mutation frequencies from LSS and HSS for the WT RBD are shown in Fig. 3A and 3B. Bars indicate the fraction of mutations per position (right axis), while circles represent the frequency of the specific mutation. Comparing LSS (Fig. 3A) and HSS (Fig. 3B) reveals that well documented Omicron mutations 440, 450, 452, 460, 477, 484, 498, and 501 ^31-36^ occur at increased frequencies in both protocols. However, the evolution under both protocols is fundamentally different. HSS fixate specific mutations at very high frequencies, contrary, LSS mutation frequencies are lower and the mutations are more variable. Logo plots for the other LSS and HSS libraries are shown in Fig. 3C and 3D, respectively demonstrating similar trends. Detailed mutation analyses are provided in Fig. S14–S18 for LSS and Table S4 and Fig. S19–S30 for HSS. For example, N460K is selected in both LSS and HSS but at much higher frequency, ∼1, in HSS vs. 0.14 in LSS. Similarly, S477N and T478K are enriched to frequencies of ∼1 in HSS, and 0.2 and 0.002 in LSS. Overall, LSS results in a greater number of mutations above the 1% frequency threshold than HSS, with 99 mutations, of which 31 are non-synonymous for LSS versus 58 of which 37 are non-synonymous for HSS (Fig. 3A). Comparing mutation frequencies after two and four rounds of HSS shows that dominant mutations were already selected by round two, but their prevalence increased by round four (Table S4, Fig. S19–S30). A detailed comparison of high frequency mutations with SARS-CoV-2 evolution is shown in Table S5. The complete list of mutations and selected frequencies can be found deposited at Zenodo.org under DOI:10.5281/zenodo.15102607 accession and also as interactive mutation scatter plots at https://host-patho-evo.github.io/mutation_scatter_plot/.

**Fig. 3.**
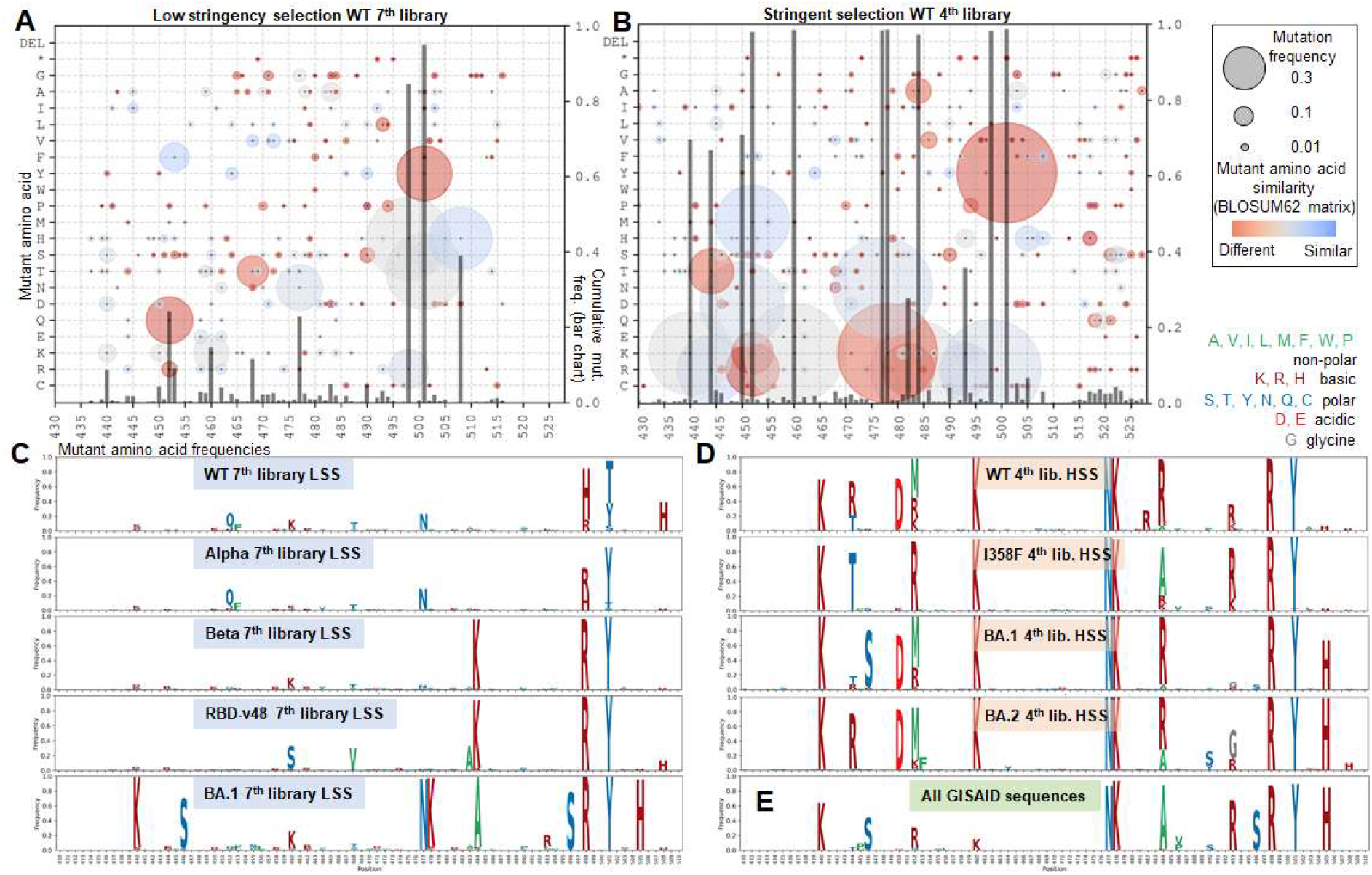
Mutation accumulation following in vitro evolution in comparison to SARS-CoV-2. (A) Mutation scatter plot of in vitro evolution under LSS. (B) Mutation scatter plot of in vitro evolution under HSS. The size of the circles reflects the frequency of the mutation in the population, while their color indicates similarity to the original amino acid as defined by BLOSUM62 ^40^. The bars represent the sum of the frequencies of mutations at each position (right y-axis). (C) Sequence Logo plots showing the frequencies of mutations from LSS libraries (color legend in upper right). (D) Sequence Logo plots showing the frequencies of mutations from HSS libraries. (D) Mutation frequencies of the Spike protein from sequences deposited in the GISAID database (10^th^ December 2024, 16.9 mil seq). More detailed analysis including logo and mutation scatter plots for all libraries and also Pango named lineages is provided in Supplementary Material S19-S31. Interactive versions for corresponding mutation scatter plots are available at https://host-patho-evo.github.io/mutation_scatter_plot/ enabling zooming, and mouse-selection-based readouts of corresponding frequencies for given position/codon.

Positions 498 and 501 are of particular interest as the double-mutation Q498R/N501Y emerged with Omicron, and is conserved since, while N501Y was established already from the Alpha lineage. Under HSS, the Q498R/N501Y double mutation was established in round 2 and did not further evolve. In contrast, LSS starting from WT led to N501Y in 30% and Q498R in 15% of clones after seven rounds, with Q498H appearing in 69% of the sequences. Notably, Q498H alone emerged rapidly under low-stringency selection in mammalian display experiments ^37^ and is associated with adaptation to mice ^34^. Furthermore, Q498H acquisition is often followed by N501T or N501S, suggesting epistasis similar to Q498R/N501Y. When LSS started from Alpha or Beta lineages, where N501Y is present, it is maintained, with Q498R emerging in 35–96% of the sequences (Table S6). When both Q498R and N501Y were present at the outset of LSS (in BA.1 and RBD-v48), they remained fixed throughout selection (Fig. 3C and Table S6). Our previous work demonstrated the synergistic effect of Q498R and N501Y ^24^, which is strongly selected under HSS and maintained under LSS, representing an evolutionary tipping point directing RBD evolution.

We included the RBD-v48 variant for LSS evolution as an intriguing experiment to evaluate how its 50-fold increased binding affinity will contribute towards evolutionary plasticity ^24^. Indeed, this increased affinity provided a “surplus” of affinity providing broader mutational tolerance, allowing the fixation of unique mutations such as A348P, A363P, K386E, N388D, N460S, I468V, V483A, and S514T identified by NGS. Available deep mutational scanning data for the WT RBD indicate minimal impact of these mutations on binding (± 0.07 Δlog10(KD,app), ^22^) likely explaining their tolerance.

Interestingly, the S375F mutation, characteristic of Omicron but not interacting with ACE2, was rapidly eliminated and reverted in the in vitro evolution (LSS BA.1), suggesting a strong selective disadvantage. The display of RBD variants with and without this mutation revealed a striking difference, indicating its dramatic structural and functional impact (Fig. S1). The continuous presence of S375F in all Omicron lineages is likely driven by factors other than adaptation to human receptor binding ^38,39^.

HSS produced a distinct mutation profile (Fig. 3D) including N440K, S477N, T478K, Q498R, and N501Y when starting from WT and WT+I358F, already after two rounds of mutagenesis. This confirms their role in enhancing ACE2 binding while maintaining protein stability. These mutations are native in BA.1 and BA.2, and did not further evolve when used as starting libraries (Table S4). N460 evolved to K from both WT and Omicron. Position 484, which changed from E (WT) to K (Beta variant) and A (in BA.1 and BA.2), evolved to R or A under HSS in WT and WT+I358F, a pattern also seen in BA.1 and BA.2 (Table S4). Position 450 evolved from N to D in WT, BA.1, and BA.2, while 452 (L in WT, R in delta) evolved to M or R under HSS in WT, WT+I358F, BA.1, and BA.2. Positions 446 and 496, G in WT and BA.2 but S in BA.1, reverted in BA.1 to G. Position 505, R in WT and H in Omicron, remained H in BA.1 and BA.2, but starting from WT it initially mutated to H before reverting to Y. Overall, our results reveal a high degree of similarity between HSS-driven evolution of WT, BA.1, and BA.2 RBDs, resulting in Omicron like variants (Fig. 3D, S19-S30). Interestingly, the SARS-CoV-2 Y453F mutation, previously observed in minks ^35,36^ and known to strongly enhance ACE2 binding affinity ^22^, was selected only under LSS. This suggests a potential incompatibility between Y453F and other affinity-enhancing mutations selected under HSS.

### High stringency selection for ACE2 binding recapitulates viral evolution

BA.1 and BA.2 were the first variants of Omicron that were identified in November 2021. These were followed by additional Omicron lineages, accompanied by a selective sweep that caused the previous lineages to disappear. To compare HSS, either starting from WT or Omicron sequences towards the overall variability of the RBD sequences as determined through virus sequencing from 2020-2024, we gathered all GISAID sequences ^41^, incorporating either the full genome set as of December 10, 2024 (16,924,617 sequences, bottom panel of Fig. 3D and Table S5), or the complete set of named Pango lineages curated on the same date (https://github.com/corneliusroemer/pango-sequences, Fig. S31) ^42^. While this analysis is biased towards countries with abundant sequencing, with the fraction of patients sequenced being sharply reduced over time, it provides a qualitative view of the evolution of the RBD of SARS-CoV-2. Table S5 shows a comparison of the overall frequencies of amino-acids calculated from GISAID (only frequencies >0.01 are shown) and those calculated from 2023 and onwards (termed advanced) in comparison to the same residues after 4 rounds of HSS starting from WT, WT+I358F, BA.1 and BA.2 sequences. The data show a clear similarity between HSS and natural evolution. From the 29 mutations with frequencies in SARS-CoV-2 >0.01, 24 were selected by HSS. Interestingly, those that were not selected had all relative low frequencies in SARS-CoV-2, which increased in the advanced Omicron lineages maybe due to an advantage in immune evasion. Also vice versa, from the 33 amino-acid variations selected by HSS with frequency of >0.03 from at least one starting sequence, 25 were evolved also in SARS-CoV-2 with frequencies >0.01.

Interestingly, K444T, N450D, L452M/R and E484R did evolve in HSS, but their frequency in SARS-CoV-2 is highly lineage specific. K444T and N450D are observed in more advanced Omicron lineages, such as BA.2.75 and BQ.1 descendants (BR.4, CH.1, BN.3.1). L452M appeared in CM and BA.2.74. L452R is frequent in many lineages including descendants of BA.5. Particularly interesting is E484R, which evolved in HSS from all four starting sequences. The natural frequency of the this mutation is relatively rare, appearing in B.1.351 ^43^, CM.x lineages and a few others. This suggests that E484R contributes to ACE2 binding (as is the case for E484K, ^22^), but is likely hindering viral fitness.

Additional mutations observed in more advanced Omicron variants, including F486P ^44,45^, and the so called Flip mutations L455F, F456L ^46,47^, were selected by HSS starting from WT and WT+I358F sequences, yet not to high-frequencies. Additionally, V445H and N481K, that were detected across all HSS libraries, are characteristic of BA.2.86 and its descendant lineages ^48,49^. This shows that in vitro evolution catches not only the highly advantaged mutations (which were rapidly fixed), but, at lower frequencies also mutations that evolved later in Omicron.

In contrast to the mutations discussed above, the mutations G446S, Q493R, G496S and Y505H exhibit distinct evolutionary trends. These residues are present in natural evolution, but did not evolve to high frequencies using LSS or HSS. Y505H did evolve in the second round of HSS, but its frequency was reduced in the 4^th^ round (Table S4). Y505H interacts with the neighboring Spike chain residue P373 in closed conformations ^50^, raising the possibility of evolutionary advantage for the closed conformation. Mutation G446S emerged in BA.1 and there persisted similarly to its persistence in both LSS and HSS BA.1 libraries. In other libraries, as also in BA.2 and its descendants, it evolved to relative low frequency (of 0.01-0.03, which is still 1000-fold above baseline). Thus, it seems that the replacement to S brings only a limited advantage over G. Similarly, Q493R appears to be context-dependent, persisting under certain conditions but being eliminated under others (Table S5). The G496S was eliminated from both natural and HSS evolution (Table S5). The transient presence of G446S, Q493R and G496S mutations, despite their eventual removal, supports the hypothesis that Omicron evolved in immunocompromised patients. In such individuals, prolonged intra-host viral evolution was likely influenced by highly specific selective pressures leading to the selection of unique mutations, which were eliminated upon transmission to the general population^51-54^. Still, we cannot exclude the possibility of adaptive evolution in an animal reservoir to different ACE2 orthologs where their fitness is positive ^3^.

### HSS and LSS selection for ACE2 binding results in tighter-binding clones

To evaluate and confirm the driving force behind the selections, we measured binding affinities and RBD thermostability for different clones in the HSS and LSS evolved libraries (Fig. 4 and Table S7). An intriguing outcome is that in all cases, HSS selection resulted in increased binding affinity by about 10-fold, reaching 0.4-1.5 nM, independent of the evolved variant (Fig. 4A). This is in line with the decreased binding affinity values of many of the SARS-CoV-2 lineages (Fig. 4). Also, the thermostability of the evolved variants were ∼55 °C,, independent of the starting library. These results clearly highlight the strong selective pressure favoring a subset of mutations creating high-affinity variants with sufficient thermostability and increased binding affinity. Binding affinities of LSS derived clones were in a broader range compared to those observed after HSS, with either retaining the affinity characteristics of their parental lineage or evolving towards tighter binding (Fig. 4C and Table S7).

**Fig. 4.**
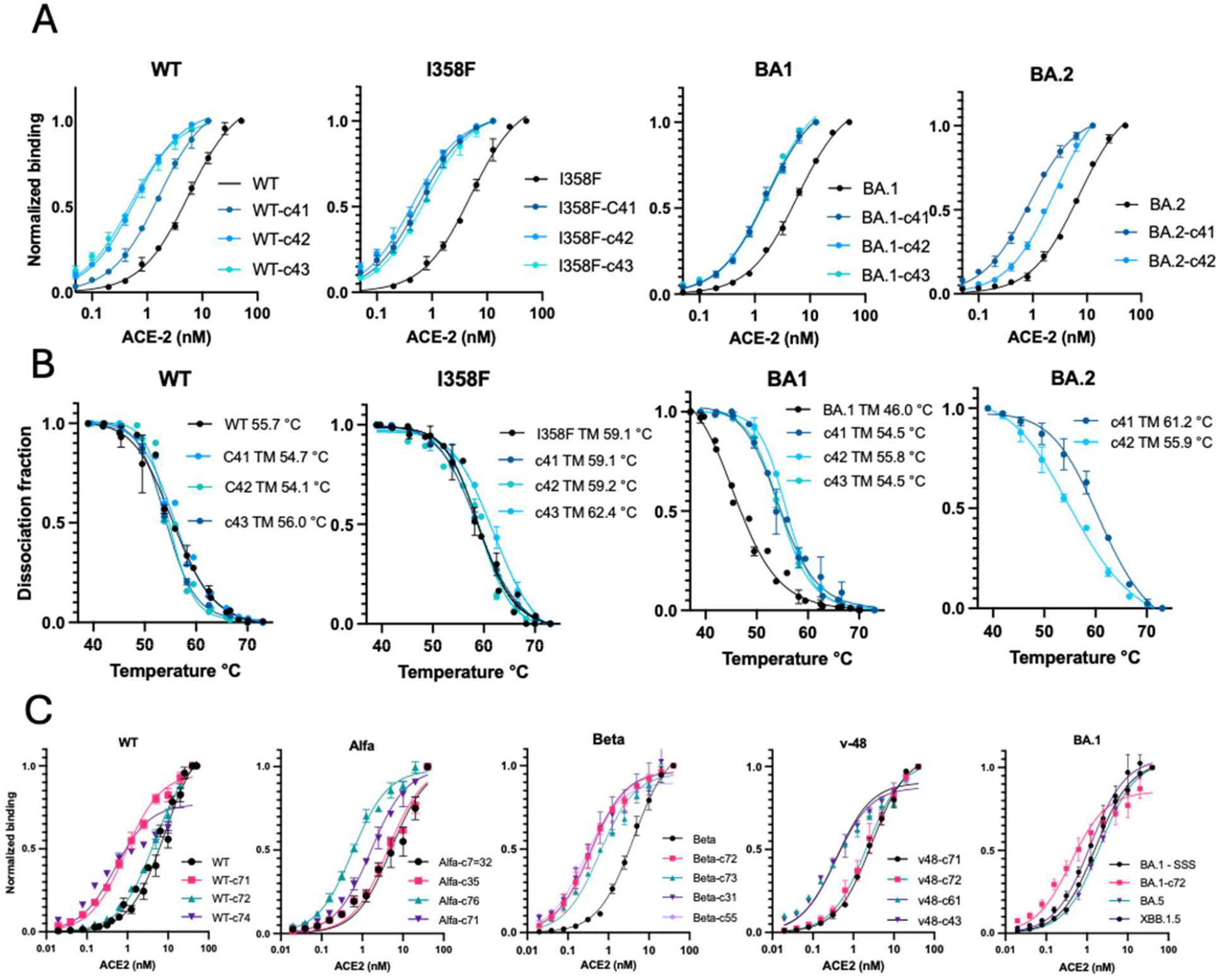
Binding affinities and melting temperatures of selected clones. (A) Binding affinities were determined by titration curves of increasing concentrations of ACE2, bound to RBD proteins displayed on the surface of yeast, with the black line representing the parental variant. The data are from HSS evolved RBD clones after 4 rounds of selection (shades of blue), compared to the parental RBD (black). (B) Melting temperature assays of RBD clones as measured by ACE2 binding of the yeast surface displayed protein. The Tm of the parental BA2 RBD clone could not be measured due to low display. (C) LSS evolved clones. The original SARS-CoV-2 lineage is designated in all cases. *K*_D_ values and sequences for the different clones are given in Table S7. Individual clones are labeled with the following code: parental_RBD, library number, clone_number e.g. Alfa-c71.

### Simulations of evolutionary trajectories

The robust convergence of mutations underscores how stringent selection conditions shape mutational landscapes, driving adaptation toward highly optimized receptor binding domains while limiting the diversity seen in low-stringency selection. This conclusion is further supported by the smaller number of synonymous mutations after HSS in comparison to LSS (Fig. 5A), which was also observed in the BA.1 and BA.2 lineages. To further support this observation, we investigated evolutionary trajectories using simulations based on a number of rules as outlined below:

**Fig. 5.**
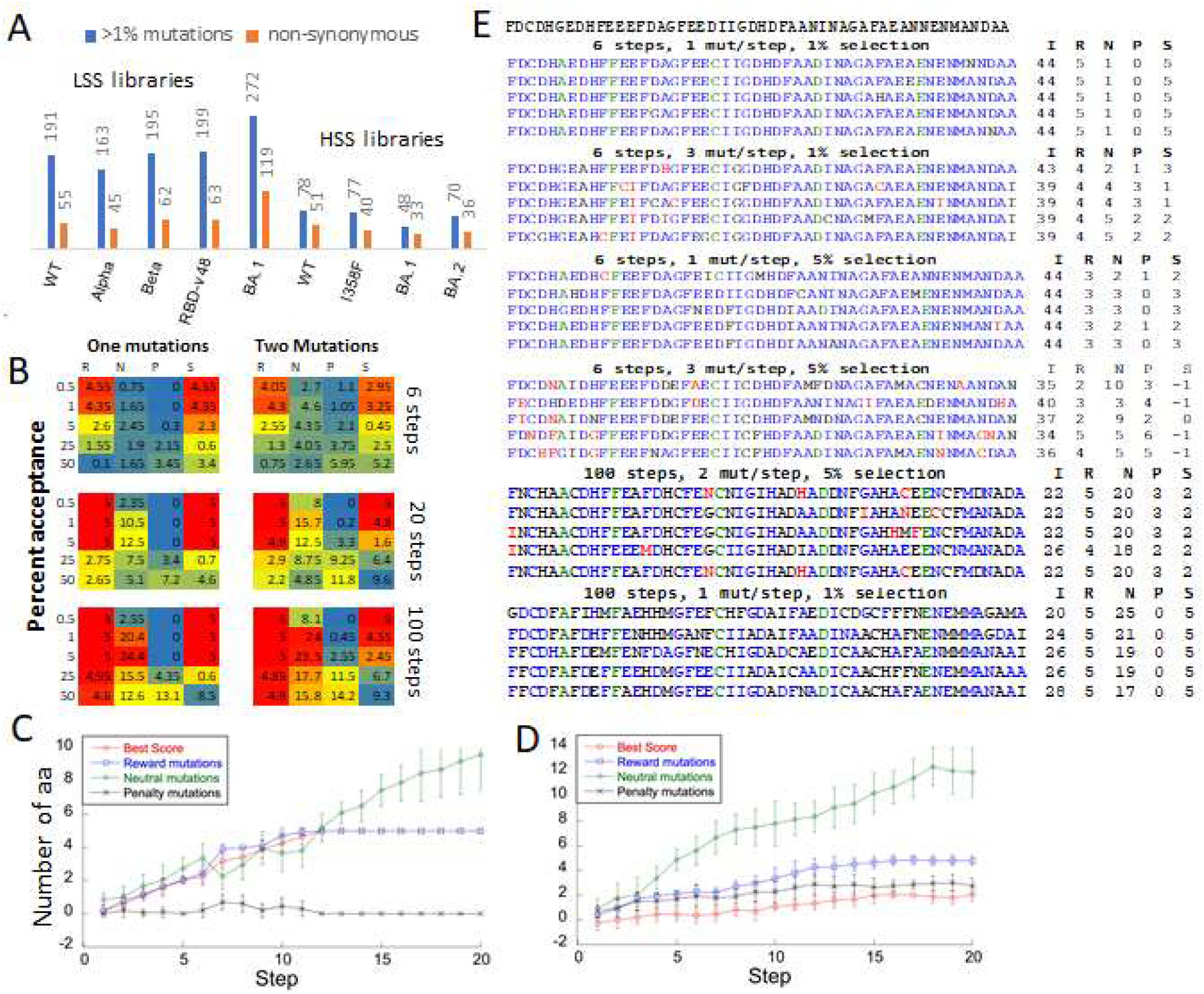
Simulating the acquisition of mutations throughout evolution. (A) The number of mutant codons (synonymous + non-synonymous) with frequencies over 0.01 in the given library in comparison to non-synonymous mutant codons following 7 and 4 rounds of LSS and HSS. (B-E) A 50-letter string composed of 10 different characters is subjected to random mutagenesis, with the top scoring sequences (0.5-50%) progressing to the next round of mutagenesis and selection. Specific positions (6, 11, 21, 31, and 41) are designated as reward (R) positions, yielding a positive score when mutated to A, F, C, D, and E, respectively. Each position permits two neutral (N) letters, including the initiating letter, while the remaining 8 (or 7 at reward positions) were assigned a negative score, penalty (P). (B) Quantification of reward, neutral, and penalty mutations, as well as final scores (S), after applying different selection stringencies (top 0.5%, 1%, 5%, 25%, and 50% of scoring sequences). Results for single mutations per step (left) and double mutations per step (right) are shown. Heatmaps (red to blue) depict the average score from 20 sequences. (C) Simulation progression over 20 steps with single mutations (2000 per step) and the top 5% of scoring sequences transferred to the next round. A rapid rise in scores is observed, plateauing at approximately 10 steps. (D) Simulation progression with double mutations (2000 per step), showing slower score improvements, reduced incorporation of reward mutations, and increased accumulation of neutral and penalty mutations. (E) Final sequences obtained after 6 and 100 steps, using selection thresholds of the top 1% or 5%.

1. As an input sequence we used a string of 50 letters, each drawn from a set of 10 different characters (arbitrary, not amino-acids). Five specific positions (6, 11, 21, 31, and 41) were designated as reward positions. Mutations to the letters A, F, C, D, and E at these positions, respectively, yielded rewards. Each position started with one neutral letter and allowed for one additional neutral letter, while the other 8 letters (or 7 at reward positions) imposed penalties.
2. Selection Criteria: At each step, the top 0.5%, 1%, 5%, 25%, and 50% of scoring strings were selected for the next round of mutagenesis and selection.
3. Simulation Setup: 2000 single or double mutations were randomly introduced per step, with varying selection stringency.

Fig. 5C illustrates simulation progression over 20 steps, introducing a single mutation per step with a 5% acceptance rate. The scores increased rapidly, plateauing after 10 steps, achieving the maximum score of 5. This is accompanied by an increase in neutral mutations at each step. Fig. 5D shows that the introduction of two mutations in parallel result in slower progression. The scores did not reach the maximum value, with fewer reward mutations introduced and a greater accumulation of neutral and penalty mutations. Fig. 5B provides a broader perspective on mutation accumulation over 6, 20, and 100 steps. When single mutations were introduced with stringent acceptance criteria (0.5% or 1%), reward residues were rapidly established (already after 6 steps), with minimal accumulation of neutral or penalty mutations. By contrast, introducing two mutations in parallel led to lower scores and greater accumulation of neutral and penalty mutations even under stringent selection (Fig. 5B). Increasing the number of steps in the simulations (20 or 100 steps) increased the number of neutral mutations at all positions, with penalty mutations appearing more frequently at lower selection stringency (Fig. 5B and 5E). The results of the simulations explain the rapid accumulation of Omicron mutations, which enhance binding affinity, within only two rounds of yeast display selection. Furthermore, the simulations explain the minimal number of bystander mutations observed in early rounds of HSS yeast display and in natural viral evolution.

## Discussion

In vitro evolution of proteins is a technique that selects for the amino-acid sequences that provide for a given trait. Here, we applied this technique to follow the sequence evolution of the RBD of the spike protein of SARS-CoV-2 upon binding to the human receptor ACE2, which is the first step in viral infection. As many of the hallmark mutations along the evolution of SARS-CoV-2 are located on the RBD (and more specifically on the RBM) we wondered whether in vitro evolution of the RBD-ACE2 binding would mimic viral evolution, and the environmental pressure driving it. For this, two regimes of evolution were employed, mild and stringent. In the mild regime, the bait protein (ACE2) concentration was in the range of WT affinity (2 nM), with 3-10% of the clones being selected for the next round. In the stringent regime, the concentration of ACE2 was gradually reduced to 0.15 nM, with 1% of the best binding clones being selected. Moreover, in the stringent regime, a heating step to 40 °C for 20 min was added, to eliminate mutations causing marginally stable prey proteins. As a result, LSS selects for a variety of sequences resulting in “good” binding, while HSS rapidly selects for the “best” binders.

LSS was done by random mutagenesis of the complete RBD starting from 5 different sequences representing different SARS-CoV-2 variants (WT, Alpha, Beta, RBD v48, BA.1). HSS was done from 4 different starting sequences (WT, WT+I358F, BA.1, and BA.2) mutating only the RBM (and not the stability-determining motif AA 337 - 430). Random mutagenesis, with ∼1 mutation per clone was verified on the starting libraries, showing that two adjacent nucleotide mutations are rare (in line with virus evolution ^7^ - Fig. 2C). Consecutive steps of random mutagenesis and selection were employed to reach substantial divergence from the starting sequence. This was achieved after 7 rounds of mutagenesis and sorting for LSS and 4 rounds for HSS. HSS rapidly converged, achieving a high frequency of mutations characteristic for Omicron already after 2 rounds of mutations (Table S4), with only few background mutations, while LSS did not converge. Detailed analysis using Sanger sequencing following 2, 3 and 4 rounds of HSS showed very little differences in the evolved mutations, with even higher frequencies of accumulation of critical mutations in individual clones (Fig. S12 and S13).

Comparing LSS to HSS shows a very different pattern of accepted mutations, with the absolute number of mutations after LSS being higher, however, most at low frequencies, with a higher number of synonymous mutations. Conversely following HSS fewer mutations were recorded, however, selected mutations were at much higher frequencies (see also Fig. S7- S30 and Table S7). Moreover, a much higher fraction of mutations were non-synonymous after HSS than after LSS, showing that HSS is more target oriented (Fig. 5A). This is in line with mutation patterns of the virus, where most mutations of the S-protein in BA.1 and BA.2 were non-synonymous ^55^ and the mutated residues were of high frequency (Fig. 3E). Simulating the conditions of high and low stringency mutations (Fig. 5) shows that HSS leads to fast accumulation of rewarding mutations, while non-rewarding mutations are scarcely selected during the first few rounds of in silico evolution (Fig. 5B and 5C). Conversely, LSS results in slower accumulation of reward mutations and higher accumulation of neutral mutations. The agreement between mutation accumulation in the RBD between the virus, in vitro evolution and in silico evolution points towards the mechanism driving mutation accumulation in the RBD and supports the substantial role of receptor adaptation in shaping SARS-CoV-2 evolution.

Notably, key mutations such as N440K, L452R, N460K, S477N, T478K, Q498R, and N501Y were enriched under HSS, mirroring their emergence in Omicron and other SARS-CoV-2 lineages. Additionally, we observed the selection of mutations associated with more advanced Omicron variants, including F486P and Flip 455–456, as well as some mutations characteristic of BA.2.86. The presence of these mutations in both in vitro and real-world viral evolution suggests that receptor-binding constraints played a crucial role in SARS-CoV-2 adaptation. Conversely, certain binding interface mutations, such as G446S, Q493R and G496S, were rapidly eliminated during in vitro evolution, suggesting structural or functional disadvantage. This supports the hypothesis that specific immune-driven selection in immunocompromised individuals, rather than receptor adaptation, was responsible for the fixation of some Omicron mutations and explains their eventual disappearance in descendant lineages. Interestingly, the non-RBM mutation S375F that destabilizes the RBD and reduces its binding affinity to ACE2 and yeast surface expression was quickly eliminated in our experiment (Fig. S11 and S18), yet it persists in the virus evolution, suggesting an overall fitness advantage. Mutations primarily associated with immune evasion, particularly in the 444–456 and 486–490 regions, were not significantly enriched in our experiments, reinforcing the idea that immune pressure, rather than receptor adaptation, drove their selection in vivo ^56-58^. Yet, they were presented in the libraries in contrast to most mutations that were excluded suggesting their tolerance and not large negative effect to binding or stability. Importantly, our experiments were intentionally designed to exclude antibodies from the selection process, allowing us to examine RBD evolution in the absence of immune-related selective pressures. This approach provides a clear view of how receptor binding alone shapes SARS-CoV-2 RBD evolution, independent of antibody-driven selection. It is important to note that although the fixation of major HSS mutations did not require immune system pressure, these mutations consequently contributed to immune escape by altering the RBD protein surface physicochemical properties. Our analysis using the BLOSUM62 score matrix (color coding, Fig. 3A, 3B, Fig. S19-S31) shows that properties of most fixed HSS mutations differ significantly from the original amino acid, including frequent mutations altering surface charges. One of the likely mechanistic explanations for these changes are improved charge complementarity of the interaction ^24^.

An interesting question that can be answered by our experiments is whether Omicron emergence, which is clearly the human adapted SARS-CoV-2 variant, was inevitable. We show that it emerged rapidly under HSS, but did not emerge under LSS even after 7 rounds of selection. However, using BA.1 as the starting library, Omicron was conserved also under LSS. This shows that HSS was crucial for the evolution of Omicron but not its maintenance, which is driven by its evolutionary advantage. Omicron did not emerge through stepwise evolution in the general public, but through a drastic shift in the mutation landscape, apparently driven by HSS within a small closed environment (maybe in immunocompromised patients). Within the general public, LSS together with immune evasion were the dominant evolutionary drivers for a long time, until Omicron emerged. Could another drastic event change the mutation landscape again? At least from the stability of Omicron mutations under our HSS and LSS conditions the answer is no. However, a drastic shift in evolutionary pressure could change this prediction.

Our findings also suggest that similar in vitro evolution can be used to predict adaptive landscapes for other viruses. The repeated and stable selection of Omicron-like RBD sequences under stringent conditions despite different starting conditions supports the feasibility of this approach for studying viral adaptation. Given that Omicron and its sub-variants have dominated SARS-CoV-2 evolution since late 2021, we postulate that Omicron represents a “humanized” form of SARS-CoV-2, optimized for human ACE2 binding and its key mutations will stay while the immune escape mutations will continue to evolve at different positions. The ability to recapitulate its emergence through directed evolution highlights the predictive power of our approach and its potential for anticipating evolutionary constraints in emerging viruses.

## Methods

### Cloning and DNA manipulations

For the modification of yeast display vectors, *S. cerevisiae* optimized genes for SpyCatcher003 and DeMA monomeric avidin (Supporting information part PS1) were purchased from Twist Bioscience and subsequently plasmids pJYDC1 and pJYDC3 (Addgene IDs 162458 and 162460) were modified by restriction free cloning ^59^ similarly to previous modifications ^26^. Barcodes (Table S1) were incorporated into amplification primers and introduced into pJYDC1-3EFR-Cfr-anti-StreptavidinAPC plasmid by homologous recombination ^26^. Non-optimized RBD genes (amino acids 330–529, Table S2, Twist Bioscience) and the previously mentioned genes were PCR-amplified using KAPA HiFi polymerase (Roche) with specific primers (Table S3). Amplicons were purified using the NucleoSpin Gel and PCR Clean-up Kit (Macherey-Nagel) and eluted in double-distilled water. Site-directed mutagenesis to introduce the I358F mutation was performed on the WT RBD using restriction-free cloning by PCR similarly to incorporation of genes ^59^. Parental plasmids were eliminated by *Dpn*I (1 h) mediated digestion, and the crude reaction mixture (0.8 µl) was directly electroporated (MicroPulser Electroporator, BioRad) into electrocompetent *E. coli* Cloni 10G cells (Lucigen). Clones were screened by colony PCR and verified by Sanger sequencing. Four RBD variants (WT, WT +I358F, BA.1, and BA.2) genes were analogously cloned to mammalian expression vector pHLsec-1^12^, verified by sequencing and produced at large quantities with high purity for transfection by using Macherey-Nagel NucleoBond Xtra Maxi kit (Macherey-Nagel).

### DNA libraries preparation

Random mutagenesis of the C-terminal domain for HSS (CTD; amino acids 430–528) was performed using the GeneMorph II Random Mutagenesis Kit. PCR conditions were optimized (template amount, cycle number) to yield 1–2 nucleotide mutation per per CTD. Resulting CTD amplicons were cloned into pJYDC vectors via RF cloning, and mutation rates were validated by colony PCR. Optimal mutagenesis was achieved using 10 ng template DNA and 30 PCR cycles. In parallel, non-mutated N-terminal domains (NTD; amino acids 330–429) were amplified using KAPA HiFi polymerase (Roche). Yeast plasmids were linearized with *Nde*I-HF and *BamH*I-HF restriction enzyme cleavage, all three DNA fragments (CTD, NTD, plasmids) were gel-purified, isolated by NucleoSpin Gel and PCR Clean-up Kit (Macherey-Nagel) and assembled by yeast mediated homologous recombination ^71^. Random mutagenesis for LSS libraries was done by previously published Taq polymerase based error prone mutagenesis procedure over the whole RBD gene with 1 to 5 nucleotide mutations per gene ^24^.

### Yeast libraries preparation

Yeast libraries were prepared by the procedure analogous to published protocol by Benatui et. al. ^60^. *Saccharomyces cerevisiae* EBY100 cells were grown in 400 ml YPD to OD_600_ ∼ 1.6 (approx. 3×10^7^ cells/ml). Following two consecutive washes with ice-cold HPLC-grade water, the cells were washed once with electroporation buffer (1 M sorbitol, 1 mM CaCl_2_) and conditioned in conditioning buffer (0.1 M LiAc, 10 mM DTT) for 30 min at 30°C. Following conditioning, cells were washed twice with electroporation buffer, resuspended in 4 ml of electroporation buffer, divided to 1 ml aliquots and incubated with DNA for 5 min on ice. For HSS each electroporation mixture contained 6 µg of mutated CTD fragment, 6 µg of NTD, and 4 µg of linearized plasmid (three component assembly, Fig. 1B). For LSS, the mixture contained 12 µg of mutated RBD gene and 4 µg of linearized plasmid. Electroporation was performed using a BioRad Micropulser (2.5 kV, 5 ms) in 2 mm cuvettes. Cells were recovered for 2 h in 1:1 YPD media + 1 M sorbitol, then transferred to SDCAA selective media (–Trp) and grown overnight at 30 °C, 225 rpm. Diluted aliquots (10^6^ cells) were plated on SD-W agar to estimate transformation efficiency (more than 1.5×10^7^ clones per library was accepted).

### Yeast cultivation and expression procedures

Aliquots of yeast libraries grown in SDCAA media overnight were collected by centrifugation (4000g, 3 minutes, 4 °C), resuspended in 1/9 expression media (1:9 glucose : galactose ratio ^26^), and expression cultures were grown overnight at 20 °C (225 rpm). For subsequent experiments, yeast expressed cells were collected by centrifugation and washed with ice cold PBSB (PBS + 1g/L BSA).

### ACE2 labeling and quantification of expression

Yeast samples with surface expressed proteins were resuspended in ice cold PBSB. Quantification of yeast surface expression levels was achieved by co-cultivation or post-cultivation incubation (1 hr) with one of the following substances depending on used plasmid: 10 nM DMSO solubilized bilirubin for pJYDC1 (eUnaG2 reporter holo-form formation, green/yellow fluorescence, excitation 498 nm, emission 527 nm), 5 nM ALFA-tagged mNeonGreen for pJYDC3 (DnbALFA reporter nanobody), 50 nM SPY-tagged mNeonGreen for pJYDC4 (SpyCatcher003 reporter), or 5 nM biotin(Avi)-tagged mNeonGreen for pJYDC6 (DeMA reporter). ACE2 protein was labeled with CF-640R succinimidyl ester (Biotium, molar ratio 1:5; 1 hr at RT), and unreacted dye was subsequently quenched with 1 mM Tris. Protein concentration was determined via A280 (NanoDrop, Thermo Scientific; ε=152,000 M^-1^cm^-1^, MW=69.3 kDa) and confirmed using the BCA assay. Labeled ACE2 at working concentration (Fig. 1C) was added to yeast samples in volumes ensuring ligand excess and incubated overnight at 4 °C on rotator (45 rpm). After incubation, cells were washed twice in ice cold PBSB and filtered through nylon mesh prior to FACS.

### Library sorting

Expression and binding labeled HSS yeast libraries were heat-treated (40 °C, 20 min) prior sorting. eUnaG2 and mNeonGreen signal spills were compensated using ProSort software. The gating strategy is shown in Fig. S1. Sorting proceeded stepwise: a minimum of 20,000 cells were sorted, expanded for 48 hr in SDCAA, induced, and re-sorted after new expression and ACE2 labeling to enrich the population. Round two usually enabled visual resolution of enriched vs. parental populations on dot plots. For HSS an additional round of sorting was applied for every library (3 consecutive sorts vs. 2 for LSS). DNAs from sorted populations were extracted using Lyticase method, and corresponding regions (either full RBD or CTD fragments) were re-amplified with in situ barcode specific primers to avoid contaminations (Table S1) for further rounds of mutagenesis and FACS. Aliquot of yeast extracted plasmids were electroporated into *E. coli* Cloni 10G cells (Lucigen) and corresponding *E. coli* mini-preps (Wizard® Plus SV Minipreps DNA Purification Systems, Promega) from selected colonies were sequences with universal primers for pJYDC vectors ^26^.

### Binding assays and affinity constant (*K*_D_) determination

Affinity constant values of selected RBD clones for different libraries were determined using the cytometry titration curve method. Small samples of yeast single clones (10ul from expression cultures, OD600 = 1.5, approximately 150,0000 cells) were co-cultivated with fixed concentration of expression labeling substrate (bilirubin 10 nM or ALFA-tagged mNeonGreen 5 nM, plasmids pJYDC4 and pJYDC6 were not used for *K*_D_ determination), and with increasing concentrations of labeled ACE2. Total of 12 concentrations were used, starting from 50 nM to 10 pM. In order to verify that ACE-2 ligand is in excess compared to RBD molecules, large incubation volumes were used (especially at low ACE2 concentrations) to prevent ligand depletion. After incubation (overnight, 4°C, mild shaking) yeast cells were washed twice with PBSB with increased concentration of BSA to 3 g/L to reduce the non-specific interactions. Cells’ fluorescence parameters were recorded by using Beckman Coulter Life Sciences CytoFLEX benchtop flow cytometer in plate format for automated acquisition. Minimum of 30 k cells in single cell population were recorded (Fig. S1). Results were analyzed using the FlowJo program. Gating was applied to normalize the expression rates across different measurements and to separate the positive population (expressing cells) from the negative population (non-expressing cells) (Fig. 1D and Fig. S1). To calculate specific binding at each single measurement, median APC-A channel intensity of the negative population was subtracted from the positive median value of the defined expression level population. The analysis done with defined expression level across measured clones increased accuracy of affinity determination for highly expression variable clones (Fig. S32). Specific binding values were plotted over corresponding ACE-2 concentrations (log Scale) and fitted to one-site specific binding equation using GraphPad v10, and best fit *K*_D_ values were generated from at least 3 replicate binding curves.

### On-surface RBD stability assay

The protein stability of surface displayed RBD variants was analyzed by heating expressed yeast cells with a gradient temperature (from 37°C to 73°C) for 14 minutes in gradient PCR cycler prior to incubation with CF-640 labeled ACE-2 (6.4 nM, 2 ml volume). Subsequently cells were washed twice with PBSB with high concentration of BSA 6 g/L. Yeasts fluorescent parameters were analyzed by using Beckman Coulter Life Sciences CytoFLEX benchtop flow cytometer, and normalized APC-A intensity (specific binding fraction) was plotted over increasing temperatures. A Sigmoidal (logistic) curve was generated using GraphPad v10 (4PL, Four-Parameter Logistic model) and IC50 (melting temperatures) were calculated from at least 3 replicate binding curves.

### Recombinant RBD and ACE2 production and purification

For recombinant RBD protein production suspension HEK293F cells were grown and transfected using PEI-MAX transfection reagent according to the manufacturer’s protocol. Briefly, each RBD variant was produced in 200 ml of cell suspension transfected with 200 ug of plasmid DNA and 600 ul of PEI reagent. The media were collected after 80 hr of incubation by centrifugation (500 g, 5 minutes) and filtered using a 0.45 µm vacuum filter unit. Secreted protein bearing C-terminal his tag was purified on Ni-NTA agarose beads (HisPur Ni-NTA Agarose Resin, ThermoFisher) and BioRad gravity flow column (Econo-Pac® Chromatography Columns, 14 ml volume), supernatant was loaded, washed with 10 CV of PBS and eluted using PBS + 300 mM Imidazole (protein purity was analysed by SDS-PAGE, see Fig. S2). The Elution buffer containing imidazole was replaced by 100% PBS using dialysis buffer exchange (GeBAflex-tube, 10 kDa MWCO). A stabilized version of the ACE2 protein (ACE2D, carrying mutations outside the receptor-binding interface) was expressed in *E*.*coli* BL21(DE3) using the plasmid pET28-bdSUMO-ACE2D, following our previously established protocol ^49^. Cells were cultured in 2×YT medium at 37 °C until reaching an optical density at 600 nm (OD_600_) of 0.6. Protein expression was induced with 0.5 mM IPTG, and cultures were incubated overnight at 20 °C. Cells were harvested by centrifugation (8,000 × g, 5 min), resuspended in lysis buffer (50 mM Tris-HCl, 300 mM NaCl, pH 8.0), and lysed by sonication. The lysate was clarified by centrifugation (16,000 × g, 30 min, 4 °C), and the supernatant was filtered and applied to a 15 mL gravity-flow column containing Ni-NTA agarose resin (HisPur Ni-NTA, Thermo Fisher Scientific). After washing with 10 column volumes (CV) of wash buffer (50 mM Tris-HCl, 300 mM NaCl, 25 mM imidazole, pH 8.0) followed by 5 CV of PBS, bdSUMO protease (50 µM) was added directly to the column. The column was incubated overnight at 4 °C with gentle shaking (60 rpm) to release the cleaved ACE2D protein. The eluted protein was subsequently purified by size-exclusion chromatography on a Superdex 75 16/600 column (Cytiva) equilibrated in PBS.

### Next generation sequencing

DNA libraries (four HSS RBD libraries at three stages: unsorted mutagenized libraries, second round of affinity maturation, fourth round of affinity maturation, and five LSS RBD libraries) were amplified using KAPA HiFi and purified using the NucleoSpin Gel and PCR Clean-up Kit. Barcodes were added as part of the amplification primers (primers and barcodes are listed in Table S3) and the number of minimal PCR amplification was determined by qPCR. PCR amplicons were sequenced on Illumina sequencers as 2×150 nt paired-end reads, and additionally as 2×250 nt paired-end reads to ensure overlap of forward and reverse reads for all amplicons. The PCR products already contained P5 and P7 Illumina sample barcodes; hence, demultiplexing was done using the standard Illumina bcl2fastq tool. Notably, the sequencing libraries were diluted with samples from other customers to avoid ghost signals. Sequencing reads were mapped against the Wuhan S protein coding sequence (GenBank accession MN908947.3) spanning from the initiator ATG to the termination codon at position 3822, comprising 1274 codons, using the “bwa mem” program (version 0.7.17-r1188). We aligned the obtained Illumina sequences using NCBI’s BLASTN to the respective reference sequence and applied a similarity cutoff of 85%, which included 99% of the data. The alignments were sorted by the start positions on the reference sequence and converted into multi-FASTA alignment (ALN) files using the “gofasta sam to MultiAlign” command (https://github.com/virus-evolution/gofasta).

For the follow-up analysis we developed in Python two programs which are now available at https://github.com/host-patho-evo/mutation_scatter_plot. Briefly, the ALN files were parsed by calculate_codon_frequencies.py using biopython (version 1.83) which iterated over the reference sequence (to respect reading-frame) and calculated frequencies of codons in a particular 3nt wide column of the multi-FASTA ALN file. The results were stored in a TAB-separated TSV file for easy post-processing by our another python-based utility named mutation_scatter_plot.py which discards codons containing unknown (N) nucleotides using python Pandas library and finally draws interactive figures using Matplotlib and Bokeh graphical libraries. To color the figures we used the widely used BLOSUM62 matrix from https://www.ncbi.nlm.nih.gov/IEB/ToolBox/C_DOC/lxr/source/data/BLOSUM62 to discern evolutionarily conservative aminoacid changes from those more drastic. The color palette “coolwarm_r” visible at https://i.sstatic.net/cmk1J.png we used for drawing ranges from red to blue (in dark red should be rather pronounced change while in dark blue should be functionally similar amino acid change). By default, mutation_scatter_plot.py omits codon changes with frequency lower than 0.001 (below one 0.1%) or when in amino acid mode it omits cumulative codon changes summed up at the resulting amino acid frequency below 0.01 (below 1%). The BAM, FASTA, TSV, HTML with javascript, PNG, JPG and PDF files for each sample were deposited at Zenodo.org under DOI:10.5281/zenodo.15102607 accession. Our software is available at https://github.com/host-patho-evo/mutation_scatter_plot and the interactive HTML files are directly accessible along JPG preview images at https://host-patho-evo.github.io/mutation_scatter_plot.

### Simulations of evolutionary trajectories

Simulations were performed as detailed in Fig. 5 and adjacent text using a Python script generated for this purpose. The program is available at …..

## Acknowledgement

The authors acknowledge Imaging Methods Core Facility at BIOCEV, institution supported by the MEYS CR (LM2023050 Czech-BioImaging) and for their support & assistance with FACS sortings. The research was supported by the project National Institute of virology and bacteriology (Programme EXCELES, ID Project No. LX22NPO5103) - Funded by the European Union - Next Generation EU, by Czech Science Foundation Grant No. 25-17643M and by the institutional grant to the Institute of Biotechnology of the Czech Academy of Sciences RVO:86652036. G.S. received funding from both the Israel Science Foundation Grant (No. 3814/19) within the KillCorona-Curbing Coronavirus Research Program and the Ben B. and Joyce E. Eisenberg Foundation of the Weizmann Institute of Science.

## References

1. Clarke, D. K. et al. Genetic bottlenecks and population passages cause profound fitness differences in RNA viruses. J Virol 67, 222–228 (1993). 10.1128/jvi.67.1.222-228.1993

2. Lumby, C. K., Nene, N. R. & Illingworth, C. J. R. A novel framework for inferring parameters of transmission from viral sequence data. PLOS Genetics 14, e1007718 (2018). 10.1371/journal.pgen.1007718

3. Markov, P. V. et al. The evolution of SARS-CoV-2. Nat Rev Microbiol 21, 361–379 (2023). 10.1038/s41579-023-00878-2

4. Amicone, M. et al. Mutation rate of SARS-CoV-2 and emergence of mutators during experimental evolution. Evol Med Public Health 10, 142–155 (2022). 10.1093/emph/eoac010

5. Simmonds, P. Rampant C→U Hypermutation in the Genomes of SARS-CoV-2 and Other Coronaviruses: Causes and Consequences for Their Short- and Long-Term Evolutionary Trajectories. mSphere 5 (2020). 10.1128/mSphere.00408-20

6. Simmonds, P. C→U transition biases in SARS-CoV-2: still rampant 4 years from the start of the COVID-19 pandemic. mBio 15, e02493–02424 (2024). 10.1128/mbio.02493-24

7. Zahradník, J., Nunvar, J. & Schreiber, G. Perspectives: SARS-CoV-2 Spike Convergent Evolution as a Guide to Explore Adaptive Advantage. Front Cell Infect Microbiol 12, 748948 (2022). 10.3389/fcimb.2022.748948

8. Cai, Y. et al. Structural basis for enhanced infectivity and immune evasion of SARS-CoV-2 variants. Science 373, 642–648 (2021). 10.1126/science.abi9745

9. Hoffmann, M. et al. SARS-CoV-2 Cell Entry Depends on ACE2 and TMPRSS2 and Is Blocked by a Clinically Proven Protease Inhibitor. Cell 181, 271-280.e278 (2020). 10.1016/j.cell.2020.02.052

10. Carrascosa-Sàez, M. et al. Cell type-specific adaptation of the SARS-CoV-2 spike. Virus Evolution 10 (2024). 10.1093/ve/veae032

11. Jackson, C. B., Farzan, M., Chen, B. & Choe, H. Mechanisms of SARS-CoV-2 entry into cells. Nature Reviews Molecular Cell Biology 23, 3–20 (2022). 10.1038/s41580-021-00418-x

12. Dejnirattisai, W. et al. SARS-CoV-2 Omicron-B.1.1.529 leads to widespread escape from neutralizing antibody responses. Cell 185, 467-484.e415 (2022). 10.1016/j.cell.2021.12.046

13. Wan, Y., Shang, J., Graham, R., Baric, R. S. & Li, F. Receptor Recognition by the Novel Coronavirus from Wuhan: an Analysis Based on Decade-Long Structural Studies of SARS Coronavirus. J Virol 94 (2020). 10.1128/jvi.00127-20

14. Carabelli, A. M. et al. SARS-CoV-2 variant biology: immune escape, transmission and fitness. Nat Rev Microbiol 21, 162–177 (2023). 10.1038/s41579-022-00841-7

15. Subissi, L. et al. An updated framework for SARS-CoV-2 variants reflects the unpredictability of viral evolution. Nature Medicine 30, 2400–2403 (2024). 10.1038/s41591-024-02949-0

16. Viana, R. et al. Rapid epidemic expansion of the SARS-CoV-2 Omicron variant in southern Africa. Nature 603, 679–686 (2022). 10.1038/s41586-022-04411-y

17. Starr, T. N. et al. Shifting mutational constraints in the SARS-CoV-2 receptor-binding domain during viral evolution. Science 377, 420–424 (2022). 10.1126/science.abo7896

18. Meng, B. et al. Altered TMPRSS2 usage by SARS-CoV-2 Omicron impacts infectivity and fusogenicity. Nature 603, 706–714 (2022). 10.1038/s41586-022-04474-x

19. Ma, W., Fu, H., Jian, F., Cao, Y. & Li, M. Immune evasion and ACE2 binding affinity contribute to SARS-CoV-2 evolution. Nature Ecology & Evolution 7, 1457–1466 (2023). 10.1038/s41559-023-02123-8

20. Wang, Q. et al. Evolving antibody evasion and receptor affinity of the Omicron BA.2.75 sublineage of SARS-CoV-2. iScience 26 (2023). 10.1016/j.isci.2023.108254

21. Beltran, A., Jiang, X. e., Shen, Y. & Lehner, B. Site-saturation mutagenesis of 500 human protein domains. Nature 637, 885–894 (2025). 10.1038/s41586-024-08370-4

22. Starr, T. N. et al. Deep Mutational Scanning of SARS-CoV-2 Receptor Binding Domain Reveals Constraints on Folding and ACE2 Binding. Cell 182, 1295-1310.e1220 (2020). 10.1016/j.cell.2020.08.012

23. Greaney, A. J. et al. Comprehensive mapping of mutations in the SARS-CoV-2 receptor-binding domain that affect recognition by polyclonal human plasma antibodies. Cell Host & Microbe 29, 463-476.e466 (2021). 10.1016/j.chom.2021.02.003

24. Zahradník, J. et al. SARS-CoV-2 variant prediction and antiviral drug design are enabled by RBD in vitro evolution. Nature Microbiology 6, 1188–1198 (2021). 10.1038/s41564-021-00954-4

25. Gagne, M. et al. RBD-based high affinity ACE2 antagonist limits SARS-CoV-2 replication in upper and lower airways. bioRxiv (2023). 10.1101/2023.06.09.544432

26. Zahradník, J. et al. A Protein-Engineered, Enhanced Yeast Display Platform for Rapid Evolution of Challenging Targets. ACS Synthetic Biology 10, 3445–3460 (2021). 10.1021/acssynbio.1c00395

27. Keeble, A. H. et al. Approaching infinite affinity through engineering of peptide–protein interaction. Proceedings of the National Academy of Sciences 116, 26523–26533 (2019). doi:10.1073/pnas.1909653116

28. Lee, J. M. et al. A Rhizavidin Monomer with Nearly Multimeric Avidin-Like Binding Stability Against Biotin Conjugates. Angewandte Chemie International Edition 55, 3393–3397 (2016). 10.1002/anie.201510885

29. Traxlmayr, M. W. & Obinger, C. Directed evolution of proteins for increased stability and expression using yeast display. Arch Biochem Biophys 526, 174–180 (2012). 10.1016/j.abb.2012.04.022

30. Zajc, C. U., Teufl, M. & Traxlmayr, M. W. Affinity and Stability Analysis of Yeast Displayed Proteins. Methods Mol Biol 2491, 155–173 (2022). 10.1007/978-1-0716-2285-8_9

31. Pastorio, C. et al. Impact of mutations defining SARS-CoV-2 Omicron subvariants BA.2.12.1 and BA.4/5 on Spike function and neutralization. iScience 26, 108299 (2023). 10.1016/j.isci.2023.108299

32. Wang, Q. et al. Key mutations in the spike protein of SARS-CoV-2 affecting neutralization resistance and viral internalization. J Med Virol 95, e28407 (2023). 10.1002/jmv.28407

33. Harvey, W. T. et al. SARS-CoV-2 variants, spike mutations and immune escape. Nat Rev Microbiol 19, 409–424 (2021). 10.1038/s41579-021-00573-0

34. Huang, K. et al. Q493K and Q498H substitutions in Spike promote adaptation of SARS-CoV-2 in mice. EBioMedicine 67, 103381 (2021). 10.1016/j.ebiom.2021.103381

35. Ren, W. et al. Mutation Y453F in the spike protein of SARS-CoV-2 enhances interaction with the mink ACE2 receptor for host adaption. PLoS Pathog 17, e1010053 (2021). 10.1371/journal.ppat.1010053

36. Bayarri-Olmos, R. et al. The SARS-CoV-2 Y453F mink variant displays a pronounced increase in ACE-2 affinity but does not challenge antibody neutralization. Journal of Biological Chemistry 296, 100536 (2021). 10.1016/j.jbc.2021.100536

37. Bate, N. et al. In vitro evolution predicts emerging SARS-CoV-2 mutations with high affinity for ACE2 and cross-species binding. PLOS Pathogens 18, e1010733 (2022). 10.1371/journal.ppat.1010733

38. Kimura, I. et al. The SARS-CoV-2 spike S375F mutation characterizes the Omicron BA.1 variant. iScience 25, 105720 (2022). 10.1016/j.isci.2022.105720

39. Liu, S. et al. The Compensatory Effect of S375F on S371F Is Vital for Maintaining the Infectivity of SARS-CoV-2 Omicron Variants. J Med Virol 97, e70242 (2025). 10.1002/jmv.70242

40. Henikoff, S. & Henikoff, J. G. Amino acid substitution matrices from protein blocks. Proc Natl Acad Sci U S A 89, 10915–10919 (1992). 10.1073/pnas.89.22.10915

41. Shu, Y. & McCauley, J. GISAID: Global initiative on sharing all influenza data - from vision to reality. Euro Surveill 22 (2017). 10.2807/1560-7917.Es.2017.22.13.30494

42. Rambaut, A. et al. A dynamic nomenclature proposal for SARS-CoV-2 lineages to assist genomic epidemiology. Nature Microbiology 5, 1403–1407 (2020). 10.1038/s41564-020-0770-5

43. Zhou, D. et al. Evidence of escape of SARS-CoV-2 variant B.1.351 from natural and vaccine-induced sera. Cell 184, 2348-2361.e2346 (2021). 10.1016/j.cell.2021.02.037

44. Uriu, K. et al. Enhanced transmissibility, infectivity, and immune resistance of the SARS-CoV-2 omicron XBB.1.5 variant. Lancet Infect Dis 23, 280–281 (2023). 10.1016/s1473-3099(23)00051-8

45. Parums, D. V. Editorial: The XBB.1.5 (‘Kraken’) Subvariant of Omicron SARS-CoV-2 and its Rapid Global Spread. Med Sci Monit 29, e939580 (2023). 10.12659/msm.939580

46. Chakraborty, C. & Bhattacharya, M. FLip mutations (L455F + F456L) in newly emerging VOI, JN.1: Its antibody and immune escape. International Immunopharmacology 133, 112146 (2024). 10.1016/j.intimp.2024.112146

47. Jian, F. et al. Convergent evolution of SARS-CoV-2 XBB lineages on receptor-binding domain 455–456 synergistically enhances antibody evasion and ACE2 binding. PLOS Pathogens 19, e1011868 (2023). 10.1371/journal.ppat.1011868

48. Bdeir, N. et al. Reverse mutational scanning of SARS-CoV-2 spike BA.2.86 identifies epitopes contributing to immune escape from polyclonal sera. Nature Communications 16, 809 (2025). 10.1038/s41467-025-55871-5

49. Tamura, T. et al. Virological characteristics of the SARS-CoV-2 BA.2.86 variant. Cell Host Microbe 32, 170-180.e112 (2024). 10.1016/j.chom.2024.01.001

50. Tamura, T. et al. Virological characteristics of the SARS-CoV-2 XBB variant derived from recombination of two Omicron subvariants. Nature Communications 14, 2800 (2023). 10.1038/s41467-023-38435-3

51. Raglow, Z. et al. SARS-CoV-2 shedding and evolution in patients who were immunocompromised during the omicron period: a multicentre, prospective analysis. Lancet Microbe 5, e235–e246 (2024). 10.1016/s2666-5247(23)00336-1

52. Smith, C. A. & Ashby, B. Antigenic evolution of SARS-CoV-2 in immunocompromised hosts. Evolution, Medicine, and Public Health 11, 90–100 (2022). 10.1093/emph/eoac037

53. Marques, A. D. et al. SARS-CoV-2 evolution during prolonged infection in immunocompromised patients. mBio 15, e00110–00124 (2024). doi:10.1128/mbio.00110-24

54. Willett, J. D. S. et al. SARS-CoV-2 rapidly evolves lineage-specific phenotypic differences when passaged repeatedly in immune-naïve mice. Communications Biology 7, 191 (2024). 10.1038/s42003-024-05878-3

55. Elssaig, E. H., Alnour, T. M. S., Ullah, M. F. & Ahmed-Abakur, E. H. Omicron SARS-CoV-2 Variants in an In Silico Genomic Comparison Study with the Original Wuhan Strain and WHO-Recognized Variants of Concern. Pol J Microbiol 71, 577–587 (2022). 10.33073/pjm-2022-053

56. Motozono, C. et al. SARS-CoV-2 spike L452R variant evades cellular immunity and increases infectivity. Cell Host Microbe 29, 1124-1136.e1111 (2021). 10.1016/j.chom.2021.06.006

57. Raharinirina, N. A. et al. SARS-CoV-2 evolution on a dynamic immune landscape. Nature 639, 196–204 (2025). 10.1038/s41586-024-08477-8

58. Meijers, M., Ruchnewitz, D., Eberhardt, J., Łuksza, M. & Lässig, M. Population immunity predicts evolutionary trajectories of SARS-CoV-2. Cell 186, 5151-5164.e5113 (2023). 10.1016/j.cell.2023.09.022

59. Peleg, Y. & Unger, T. Application of the Restriction-Free (RF) cloning for multicomponents assembly. Methods Mol Biol 1116, 73–87 (2014). 10.1007/978-1-62703-764-8_6

60. Benatuil, L., Perez, J. M., Belk, J. & Hsieh, C. M. An improved yeast transformation method for the generation of very large human antibody libraries. Protein Eng Des Sel 23, 155–159 (2010). 10.1093/protein/gzq002

