## Supplementary material for "In Vitro and Viral Evolution Convergence Reveal the Selective Pressures Driving Omicron Emergence"

#### Supporting information part PS1 - platform for parallelized In vitro evolution

The pJYDC4 plasmid was created using a cut-and-paste replacement of the eUnaG2 reporter with SpyCatcher003. Similarly, the pJYDC6 plasmid was constructed using a designed monomeric avidin (DeMA). The design of this reporter was based on the PDB structure 1avd of rhizavidin. The design process was identical to the preparation of the eUnaG2 and DnbALFA reporters in Zahradník, J. *et al.* A Protein-Engineered, Enhanced Yeast Display Platform for Rapid Evolution of Challenging Targets. *ACS Synthetic Biology* 10, 3445-3460 (2021). https://doi.org:10.1021/acssynbio.1c00395

>SpyCatcher003_S.cer_optimized

ATGGTTACCACACTGAGCGGTCTGAGTGGTGAACAGGGTCCGAGCGGTGATATG
ACCACCGAAGAAGATAGCGCAACCCATATCAAATTTAGCAAACGTGATGAAGAT
GGTCGTGAACTGGCAGGCGCAACCATGGAACTGCGTGATAGCAGCGGTAAAACC
ATTAGCACCTGGATTAGTGATGGTCACGTGAAAGATTTTTATCTGTATCCGGGTA
AATATACCTTCGTTGAAACCGCAGCACCGGATGGTTATGAAGTTGCAACCCCGAT
TGAATTCACCGTTAACGAAGATGGCCAGGTTACCGTTGATGGTGAAGCAACCGAA
GGTGATGCACATACC

>DeMA_S.cer_optimized

TTCGATGCCTCTAACTTCAAGGACTTCTCTTCTATTGCTGGTACTTCTACTACCTG
GCAAAATCAACATGGTTCCACTATGGTTATCACCGTTGATTCTCAAGGTAACGTTT
CTGGTCAATACGTTAATAGAGCTGAAGGTACTGGTTGTCAGAATTCTCCATATCC

ATTGACTGGTTGGGTTAACGGTACTTTCATTGATTTCTCTGTTACCTGGAACAACT
CTACCGAAAACTGTAACTCTGTTACTCAATGGACTGGTTACGCTCAAGTTAATGG
TAACAACACTGAAATCGTTACCGATTGGAACTTGGTTTATGAAGGTCCATCTGGT
CCAGCTATTTGGCAAGGTCAAGATACTTTTCAATACGTCCCAACTACCGAG

pJYDC-barcF-RBDo-Wu plasmid sequence segment with expression of whole yeast display cassette used: eUnaG2 – Aga2p – barcode_F_fwd - RBDo-Wu_sequence - barcode_F_rev.

Left and right barcodes were integrated flanking the expression cassette within the expression construct. Their sequences are in the Table S1

>Expression_construct

ATGAGGTTCCCATCTATTTTCACCGCTGTTGTTTTTGCTGCTTCTTCTGCTTTGGCTGCTCCAGCTAA
TGGTATGTTAGAAAAATTTGTTGGCACCTGGAAGATCGAATCCTCTGAAAATTTTGGTGAATACTT
GAAGGCTATCGGTGCCCCAAAAGAATTGGCTGATGCTGGTGATGCTACTACTCCAGTCTTGTACAT
TTCTCAAAAGGATGGTGATAAGATGACCGTCAAGATTGAAAACGGTCCACCAACTTTTTTGGATAC
CCAAGTTTCTTTCAAGTTGGGTGAAGAATTCGACGAATTTCCATCCGATAGAAGAAAGGGTGTTAA
GTCCGTTGTTAACTTGTCTGGTGAGAAGTTGGTTTACGTTCAAAAGTGGGATGGTAAAGAAACCAC
TTACGTCAGAGAAATCAAGGACGGTAAATTGGTTGTTACTTTGACCATGGGTGATGTTGTTGCTGTT
AGATCTTATAGAAGGGCCTCTGAAGTTTCT**GCACAGGAACTGACAACTATATGCGAGCAAATCC**
**CCTCACCAACTTTAGAATCGACGCCGTACTCTTTGTCAACGACTACTATTTTGGCCAACGGGA**
**AGGCAATGCAAGGAGTTTTTGAATATTACAAATCAGTAACGTTTGTCAGTAATTGCGGTTCTC**
**ACCCCTCAACAACTAGCAAAGGCAGCCCCATAAACACACAGTAT**GTTTTTAAGGACAATAGCTC
GACGattgaaggtagatacccatacgacgttccagactacgctctgcaggctagtggtggaggaggctctggtggaggcggtagcggaggcggaCAAG
GTACCGGAAGTACAAGTgctagccatatgggtTGCCCTTTTGGTGAAGTTTTTAACGCCACCAGATTTGCAT
CTGTTTATGCTTGGAACAGGAAGAGAATCAGCAACTGTGTTGCTGATTATTCTGTCCTATATAATTC
CGCATCATTTTCCACTTTTAAGTGTTATGGAGTGTCTCCTACTAAATTAAATGATCTCTGCTTTACTA
ATGTCTATGCAGATTCATTTGTAATTAGAGGTGATGAAGTCAGACAAATCGCTCCAGGGCAAACTG
GAAAGATTGCTGATTATAATTATAAATTACCAGATGATTTTACAGGCTGCGTTATAGCTTGGAATTC
TAACAATCTTGATTCTAAGGTTGGTGGTAATTATAATTACCTGTATAGATTGTTTAGGAAGTCTAAT
CTCAAACCTTTTGAGAGAGATATTTCAACTGAAATCTATCAGGCCGGTAGCACACCTTGTAATGGT
GTTGAAGGTTTTAATTGTTACTTTCCTTTACAATCATATGGTTTCCAACCCACTAATGGTGTTGGTTA
CCAACCATACAGAGTAGTAGTACTTTCTTTTGAACTTCTACATGCACCAGCAACTGTTTGTGGACCT
AAAggatccAGATCCGGAAATGGCAGTACCgaacaaaagcttatttctgaagaggacttgtaa

- in red short AppS4 leader sequence

- in green eUnaG2 reporter protein

- in bold Aga2p sequence

- yellow background highlighted sequences – fwd and rev barcodes

- in blue RBD sequence

- in magenta c-myc sequence

**Table S1. Expression cassette barcodes for identification of plasmids in parallelized In vitro evolution**

| Barcode name | Barcode sequence | AA sequence |
|---|---|---|
| A_fwd | AGCAACGGTACTGGATCTAGT | SNGTGSS |
| A_rev | GGAGGCAACTCAAGTGGAACT | GGASSGT |
| B_fwd | AACACTTCAGGAGGTAGAGGA | NTSGGRG |
| B_rev | ACCGGATCTAATGGTTCAGGT | TGSNGSG |
| C_fwd | TCCTCTGGAACTAATGGCACA | SSGTNGT |
| C_rev | GGTTCAACTGGCGGTTCCTCG | GSTGGSS |
| D_fwd | acaaacggttccggcacaggc | TNGSGTG |
| D_rev | aataccggttctggaaccgca | NTGSGTA |
| E_fwd | AGTGGCAATGCAACTGGTAGC | SGNATGS |
| E_rev | TCGGGCAATAGTGGATCATCT | SGNSGSS |
| F_fwd | CAAGGTACCGGAAGTACAAGT | QGTGSTS |
| F_rev | AGATCCGGAAATGGCAGTACC | RSGNGST |
| G_fwd | aacagcaccggcggcagcggc | NSTGGSG |
| G_rev | gctcgcgctgccggtgcggcc | GRTGSAS |
| H_fwd | accaccaacagcagcggcagc | TTNSSGS |
| H_rev | gccgccgctgttcgcggtgcg | RTANSGG |

- lowercase letters show barcodes that were not used due to potential expression influence

**Table S2. Initial RBD sequences, barcodes and plasmids**

| Original SARS-CoV2 sequence (Low Stringency selections) | Barcode | Plasmid (1st round) |
|---|---|---|
| isolate Wuhan-Hu-1 (MN908947.3) | FF | pJYDC3 |
| isolate SARS-CoV-2/human/ITA/VA-English-2021-01-22/2021 (MW521144.1) | CC | pJYDC3 |
| isolate SARS-CoV-2/human/ITA/VA-South Africa 2021-02-03/2021 (MW560269.1) | AA | pJYDC3 |
| MW560269.1 + Q498R (Abbreviated RY) | BB | pJYDC3 |
| Omicron BA.1; isolate SARS-CoV-2/human/USA/TX-CDC-ASC210646153/2022 (OM652834.1) | EE | pJYDC3 |
| Original SARS-CoV2 sequence (Low Stringency selections) | Barcode | Plasmid (1st round) |
| isolate Wuhan-Hu-1 (MN908947.3) | AA | pJYDC1 |
| isolate Wuhan-Hu-1 (MN908947.3) + I358F mutation introduced by SDM | BB | pJYDC3 |
| Omicron BA.1; isolate SARS-CoV-2/human/USA/TX-CDC-ASC210646153/2022 (OM652834.1) | C_fwd - | pJYDC1 |
| Omicron BA.2, isolate SARS-CoV-2/human/USA/CA-CDC-LC0582623/2022 (ON373904.1) | - C_rev | pJYDC3 |

- combination of barcodes A_fwd + A_rev is simplified to AA; C_fwd – means that only forward barcode was used in this case.

**Table S3. Illumina paired-end deep sequencing primers and barcodes**

| Sample | Forward primer | Fw-seq-barcode | Reverse primer | Rev-seq-barcode |
|---|---|---|---|---|
| HSS WT lib non-sorted | TTGCTGATTATAATTATAAATTACCAGATGATTTT | Not used | CAAGTCCTCTTCAGAAATAAGCTTTTGTTCGGATCC | AGTTCCACTTGAGTTGCCTCC |
| HSS I358F non-sorted | TTGCTGATTATAATTATAAATTACCAGATGATTTT | Not used | CAAGTCCTCTTCAGAAATAAGCTTTTGTTCGGATCC | TCCTGAACCATTAGATCCGGT |
| HSS BA1 non-sorted | TTGCTGATTATAATTATAAATTACCAGATGATTTT | Not used | CAAGTCCTCTTCAGAAATAAGCTTTTGTTCGGATCC | TGTGCCATTAGTTCCAGAGGA |
| HSS BA2 non-sorted | TTGCTGATTATAATTATAAATTACCAGATGATTTT | Not used | CAAGTCCTCTTCAGAAATAAGCTTTTGTTCGGATCC | CGAGGAACCGCCAGTTGAACC |
| HSS WT lib 2 | TTGCTGATTATAATTATAAATTACCAGATGATTTT | Not used | CAAGTCCTCTTCAGAAATAAGCTTTTGTTCGGATCC | AGTTCCACTTGAGTTGCCTCC |
| HSS I358F lib 2 | TTGCTGATTATAATTATAAATTACCAGATGATTTT | Not used | CAAGTCCTCTTCAGAAATAAGCTTTTGTTCGGATCC | TCCTGAACCATTAGATCCGGT |
| HSS BA1 lib 2 | TTGCTGATTATAATTATAAATTACCAGATGATTTT | Not used | CAAGTCCTCTTCAGAAATAAGCTTTTGTTCGGATCC | TGTGCCATTAGTTCCAGAGGA |
| HSS BA2 lib 2 | TTGCTGATTATAATTATAAATTACCAGATGATTTT | Not used | CAAGTCCTCTTCAGAAATAAGCTTTTGTTCGGATCC | CGAGGAACCGCCAGTTGAACC |
| HSS WT lib 4 | TTGCTGATTATAATTATAAATTACCAGATGATTTT | Not used | CAAGTCCTCTTCAGAAATAAGCTTTTGTTCGGATCC | AGTTCCACTTGAGTTGCCTCC |
| HSS I358F lib 4 | TTGCTGATTATAATTATAAATTACCAGATGATTTT | Not used | CAAGTCCTCTTCAGAAATAAGCTTTTGTTCGGATCC | TCCTGAACCATTAGATCCGGT |
| HSS BA1 lib 4 | TTGCTGATTATAATTATAAATTACCAGATGATTTT | Not used | CAAGTCCTCTTCAGAAATAAGCTTTTGTTCGGATCC | TGTGCCATTAGTTCCAGAGGA |
| HSS BA2 lib 4 | TTGCTGATTATAATTATAAATTACCAGATGATTTT | Not used | CAAGTCCTCTTCAGAAATAAGCTTTTGTTCGGATCC | CGAGGAACCGCCAGTTGAACC |
| LSS WT | GTACAAGTGCTAGCCATATGGGTTGCC | GCCCCTCT | GGCCTGATAGATTTCAGTTGAAATATCTCTC | TTCTCAGA |
| LSS Alpha | ATGGCACAGCTAGCCATATGGGTTG | AACTTTAC | GGCCTGATAGATTTCAGTTGAAATATCTCTC | CCGTACAC |
| LSS Beta | GGATCTAGTGCTAGCCATATGGGTTG | GACAGCCC | ACCGGCCTGATAGATTTCAGTTG | TGCCGTGG |
| LSS RBD-v48 | GTAGAGGAGCTAGCCATATGGGTTG | ACGCAATC | ACCGGCCTGATAGATTTCAGTTG | GCACCTAG |
| LSS BA.1 | CTGGTAGCGCTAGCCATATGGGTAC | AGATGAGT | GGCCTGATAGATTTCAGTTGAAATATCTCTC | CTCACAAT |

**Table S4. Codon frequencies upon HSS evolution of the RBM**

| Position | WT aa | BA.1 aa | BA2 aa | Mutant aa | WT codon | Mutant codon | Freq Lib WT | Freq WT 2nd | Freq WT 4th | Freq Lib I358F | Freq I358F 2nd | Freq I358F 4th | Freq Lib BA.1 | Freq BA.1 2nd | Freq BA.1 4th | Freq Lib BA.2 | Freq BA.2 2nd | Freq BA.2 4th |
|---|---|---|---|---|---|---|---|---|---|---|---|---|---|---|---|---|---|---|
| 440 | N | K | K | K | AAT | AAG | 5.4E-04 | 5.3E-01 | 6.8E-01 | 3.1E-04 | 5.5E-01 | 8.6E-01 | 9.8E-01 | 9.9E-01 | 9.7E-01 | 9.7E-01 | 9.9E-01 | 9.7E-01 |
| 444 | K | | | T | AAG | ACG | 3.8E-03 | 2.1E-01 | 1.9E-01 | 3.7E-03 | 5.5E-01 | 7.6E-01 | 4.1E-03 | 2.7E-01 | 1.1E-01 | 4.1E-03 | 9.7E-03 | 8.9E-03 |
| 444 | K | | | R | AAG | AGG | 1.8E-03 | 3.2E-01 | 4.6E-01 | 3.1E-03 | 2.2E-03 | 1.2E-03 | 8.4E-04 | 1.5E-01 | 6.7E-02 | 3.2E-03 | 6.7E-01 | 7.5E-01 |
| 445 | V | | | V | GTT | GTC | 2.2E-03 | 2.3E-03 | 7.4E-03 | 4.0E-03 | 3.0E-03 | 7.8E-03 | 1.0E-03 | 3.5E-01 | 5.7E-01 | 3.8E-03 | 1.7E-03 | 6.9E-03 |
| 446 | G | S | G | S | GGT | AGT | 2.3E-03 | 8.6E-03 | 1.2E-02 | 2.8E-03 | 2.0E-02 | 3.4E-02 | 9.9E-01 | 5.4E-01 | 7.8E-01 | 2.9E-03 | 2.2E-03 | 5.5E-03 |
| 450 | N | | | D | AAT | GAT | 1.5E-03 | 6.2E-01 | 6.9E-01 | 2.1E-03 | 9.3E-04 | 4.3E-03 | 9.0E-04 | 5.6E-01 | 7.3E-01 | 2.1E-03 | 7.1E-01 | 8.2E-01 |
| 452 | L | | | K | CTG | AAG | 8.0E-06 | 1.5E-02 | 1.6E-01 | 7.0E-06 | 1.5E-05 | 1.6E-05 | 4.0E-06 | 4.6E-03 | 3.6E-02 | 1.1E-05 | 5.5E-03 | 9.8E-02 |
| 452 | L | | | M | CTG | ATG | 2.6E-03 | 6.1E-01 | 5.3E-01 | 3.1E-03 | 1.8E-03 | 1.2E-03 | 2.3E-03 | 5.6E-01 | 6.9E-01 | 3.8E-03 | 7.1E-01 | 7.2E-01 |
| 452 | L | | | R | CTG | CGG | 8.9E-04 | 2.8E-01 | 2.7E-01 | 7.7E-04 | 5.8E-01 | 9.1E-01 | 7.5E-04 | 4.0E-01 | 2.6E-01 | 8.5E-04 | 1.3E-02 | 1.3E-02 |
| 453 | Y | | | F | TAT | TTT | 2.2E-03 | 1.6E-02 | 6.4E-03 | 3.4E-03 | 1.2E-01 | 2.1E-02 | 1.3E-03 | 5.9E-03 | 3.8E-03 | 3.2E-03 | 1.4E-01 | 1.7E-01 |
| 460 | N | | | K | AAT | AAA | 1.8E-03 | 3.0E-01 | 2.3E-01 | 2.2E-03 | 5.9E-01 | 9.2E-01 | 1.4E-03 | 3.4E-01 | 2.3E-01 | 2.1E-03 | 1.5E-01 | 1.7E-01 |
| 460 | N | | | K | AAT | AAG | 2.7E-04 | 6.3E-01 | 7.4E-01 | 2.4E-04 | 9.2E-03 | 2.9E-02 | 2.3E-04 | 5.8E-01 | 7.4E-01 | 2.7E-04 | 7.2E-01 | 8.1E-01 |
| 477 | S | N | N | N | AGC | AAC | 4.8E-03 | 8.4E-01 | 9.8E-01 | 5.1E-03 | 7.5E-01 | 9.8E-01 | 9.9E-01 | 9.9E-01 | 9.9E-01 | 9.8E-01 | 9.9E-01 | 9.9E-01 |
| 478 | T | K | K | K | ACA | AAA | 1.3E-03 | 8.1E-01 | 9.7E-01 | 1.1E-03 | 6.4E-01 | 9.5E-01 | 9.9E-01 | 9.9E-01 | 9.8E-01 | 9.7E-01 | 9.9E-01 | 9.7E-01 |
| 482 | G | | | R | GGT | CGT | 3.8E-04 | 1.9E-01 | 2.6E-01 | 2.9E-04 | 7.0E-06 | 7.2E-04 | 3.2E-04 | 1.0E-05 | 2.9E-03 | 3.5E-04 | 4.0E-06 | 2.0E-05 |
| 484 | E | | | R | GAA | AGA | 3.7E-05 | 4.6E-01 | 9.0E-01 | 1.6E-05 | 7.3E-03 | 1.0E-01 | 3.1E-05 | 6.3E-01 | 9.0E-01 | 2.4E-05 | 6.0E-01 | 7.1E-01 |
| 484 | E | A | A | A | GAA | GCA | 9.4E-04 | 4.0E-01 | 6.3E-02 | 1.1E-03 | 5.7E-01 | 6.4E-01 | 9.8E-01 | 3.6E-01 | 6.2E-02 | 9.8E-01 | 3.9E-01 | 2.7E-01 |
| 486 | F | | | V | TTT | GTT | 3.0E-03 | 2.7E-01 | 2.6E-02 | 2.6E-03 | 4.2E-01 | 5.2E-02 | 3.1E-03 | 1.2E-01 | 1.3E-02 | 2.6E-03 | 3.8E-03 | 2.5E-03 |
| 493 | Q | | | K | CAA | AAA | 6.5E-04 | 3.0E-04 | 5.0E-02 | 6.8E-04 | 8.6E-03 | 1.7E-01 | 2.0E-06 | 5.9E-04 | 8.0E-03 | 2.0E-06 | 1.9E-04 | 8.5E-05 |
| 493 | Q | R | R | R | CAA | CGA | 2.5E-03 | 1.0E-02 | 2.7E-01 | 4.0E-03 | 1.1E-01 | 6.5E-01 | 9.9E-01 | 5.7E-02 | 1.6E-02 | 9.8E-01 | 1.8E-01 | 1.5E-01 |
| 493 | Q | | | G | CAA | GGA | 3.0E-06 | 1.0E-06 | 2.5E-04 | 2.0E-06 | 3.0E-05 | 3.0E-04 | 7.1E-04 | 1.1E-01 | 7.1E-02 | 7.0E-04 | 1.5E-01 | 3.9E-01 |
| 496 | G | S | G | S | GGT | AGT | 1.6E-03 | 6.5E-03 | 7.7E-03 | 2.0E-03 | 3.5E-02 | 1.3E-02 | 9.9E-01 | 2.0E-01 | 6.7E-02 | 2.2E-03 | 4.0E-04 | 1.5E-03 |
| 498 | Q | | | H | CAA | CAT | 2.5E-03 | 9.1E-03 | 2.5E-05 | 4.4E-03 | 2.4E-01 | 2.9E-02 | 2.6E-05 | 1.2E-04 | 4.4E-05 | 3.5E-05 | 1.8E-05 | 3.0E-06 |
| 498 | Q | R | R | R | CAA | CGA | 3.4E-03 | 9.1E-01 | 9.0E-01 | 5.2E-03 | 7.0E-01 | 9.2E-01 | 9.9E-01 | 9.9E-01 | 9.5E-01 | 9.8E-01 | 9.8E-01 | 9.8E-01 |
| 501 | N | | | T | AAT | ACT | 5.2E-04 | 2.5E-03 | 3.0E-06 | 6.1E-04 | 2.3E-01 | 3.5E-02 | 2.0E-06 | 6.8E-05 | 2.6E-05 | 3.0E-06 | 2.3E-05 | 1.0E-06 |
| 501 | N | Y | Y | Y | AAT | TAT | 1.8E-03 | 9.6E-01 | 9.8E-01 | 2.4E-03 | 7.3E-01 | 9.6E-01 | 9.9E-01 | 9.9E-01 | 9.9E-01 | 9.8E-01 | 9.9E-01 | 9.8E-01 |
| 505 | Y | H | H | H | TAC | CAC | 2.1E-03 | 3.6E-01 | 6.2E-02 | 3.6E-03 | 1.2E-01 | 3.6E-02 | 9.9E-01 | 9.6E-01 | 6.6E-01 | 9.8E-01 | 9.9E-01 | 9.5E-01 |

Mutant codon frequencies (Freq), as determined from NGS of the non-selected first library (Lib) in comparison to the 2nd and 4th round of library propagation and selection (2nd and 4th). Codons with frequencies of >0.1 in at least one library are shown. Mutants are in relation to WT. Blue is for low and red for high frequencies.

**Table S5. Frequencies of amino-acid mutations in RBD of the Spike protein of SARS-CoV-2 compared to frequencies of mutations in HSS libraries.**

| Residue | WT | Variant | 1 Frequency SARS-CoV-2 | 2 Frequency SARS-CoV-2 (2023+) | 3 Frequency HSS WT | 4 Frequency HSS I358F | 5 Frequency HSS BA.1 | 6 Frequency HSS BA.2 |
|---|---|---|---|---|---|---|---|---|
| **440** | N | K | 0.638 | 0.935 | 0.684 | 0.857 | 0.973 | 0.974 |
| **444** | K | T | 0.037 | 0.000 | 0.198 | 0.762 | 0.109 | 0.009 |
| **445** | V | P | 0.079 | 0.000 | 0.002 | 0.024 | 0.000 | 0.000 |
| **445** | V | H | 0.014 | 0.283 | | | | |
| **446** | G | S | 0.444 | 0.819 | 0.012 | 0.034 | 0.778 | 0.005 |
| **446** | G | G | 0.543 | 0.181 | 0.961 | 0.940 | 0.453 | 0.975 |
| **450** | N | D | 0.018 | 0.298 | 0.694 | 0.004 | 0.733 | 0.816 |
| **452** | L | R | 0.269 | 0.167 | 0.265 | 0.913 | 0.256 | 0.013 |
| **452** | L | Q | 0.023 | 0.001 | 0.001 | 0.003 | 0.000 | 0.019 |
| **452** | L | W | 0.014 | 0.292 | | | | |
| **455** | L | S | 0.013 | 0.282 | | | | |
| **456** | F | L | 0.020 | 0.341 | 0.001 | 0.011 | 0.000 | 0.000 |
| **460** | N | K | 0.156 | 0.902 | 0.973 | 0.945 | 0.976 | 0.974 |
| **477** | S | N | 0.867 | 0.951 | 0.978 | 0.980 | 0.989 | 0.990 |
| **478** | T | K | 0.927 | 0.849 | 0.971 | 0.951 | 0.983 | 0.974 |
| **478** | T | R | 0.011 | 0.096 | | | | |
| **481** | N | K | 0.010 | 0.298 | 0.011 | 0.001 | 0.006 | 0.001 |
| **484** | E | R | 0.001 | 0.002 | 0.900 | 0.105 | 0.903 | 0.709 |
| **484** | E | A | 0.860 | 0.651 | 0.063 | 0.641 | 0.062 | 0.271 |
| **484** | E | K | 0.018 | 0.290 | 0.003 | 0.078 | 0.003 | 0.002 |
| **486** | F | V | 0.108 | 0.114 | 0.026 | 0.052 | 0.013 | 0.003 |
| **486** | F | P | 0.068 | 0.777 | | | | |
| **490** | F | S | 0.070 | 0.506 | 0.017 | 0.035 | 0.004 | 0.198 |
| **493** | Q | R | 0.675 | 0.000 | 0.273 | 0.670 | 0.016 | 0.146 |
| **496** | G | S | 0.541 | 0.000 | 0.008 | 0.013 | 0.067 | 0.002 |
| **496** | G | G | 0.453 | 0.999 | 0.972 | 0.951 | 0.794 | 0.967 |
| **498** | Q | R | 0.872 | 0.951 | 0.972 | 0.953 | 0.985 | 0.994 |
| **501** | N | Y | 0.894 | 0.953 | 0.983 | 0.958 | 0.994 | 0.993 |
| **505** | Y | H | 0.851 | 0.950 | 0.062 | 0.036 | 0.667 | 0.982 |

WT corresponds to the amino-acid in the isolate Wuhan-Hu-1 (MN908947.3) sequence and Variant is the amino-acid which frequency is shown. [1] Frequencies as calculated from the GISAID database. Only frequencies >0.01 are shown. [2] Frequencies as calculated from HSS of WT. [3] Frequencies as calculated from HSS of WT+I358F. [4] Frequencies as calculated from HSS of BA.1. [5] Frequencies as calculated from HSS of BA.2. Empty field (ND) is for HSS frequencies which were <0.01.

**Table S6 – Frequencies of combinations of mutations in the RBD between residues 498 and 501.**

| Library name | Sequence Occurence (%) | | | | | | Other notable than previous > 0.01 | Total sequences fully spanning the motif* |
|---|---|---|---|---|---|---|---|---|
| | 498-QPTN-501 | 498-QPTY-501 | 498-HPTN-501 | 498-HPTY-501 | 498-HPTT/S-501 | 498-RPTY-501 | | |
| **WT 7th lib LSS** | 0.280 | 14.030 | 4.420 | 0.470 | 56.210 | 15.060 | ('HPTS', 6.99), ('QPTT', 0.69), ('HPST', 0.31), ('RPTT', 0.24), ('HPTA', 0.17), ('QPTF', 0.12), ('HPTI', 0.1) | 7865410 |
| **Alpha 7th lib LSS** | 1.430 | 38.630 | 11.920 | 0.460 | 6.910 | 36.190 | ('HPTS', 2.37), ('HPTA', 0.48), ('QPTT', 0.42), ('QPTF', 0.16), ('RPAY', 0.13) | 12873967 |
| **Beta 7th lib LSS** | 0.220 | 2.940 | 0.230 | 0.010 | 0.080 | 93.840 | ('RPAY', 0.37), ('QPTF', 0.33), ('RPTD', 0.22), ('RPTF', 0.16), ('RPSY', 0.12), ('RSTY', 0.11) | 20922 |
| **RBD v48 7th lib LSS** | 0.400 | 0.450 | 0.070 | 0.000 | 0.100 | 97.750 | ('RPAY', 0.24), ('RPSY', 0.2), ('RPTD', 0.1) | 3211512 |
| **RBD BA.1 7th lib LSS** | 0.430 | 0.100 | 0.030 | 0.000 | 0.010 | 97.470 | ('RPAY', 0.54), ('RPSY', 0.51), ('LPTY', 0.14), ('RPTD', 0.13) | 6918311 |
| **WT 4th lib HSS** | 0.910 | 0.130 | 0.000 | 0.000 | 0.000 | 96.360 | ('RRTY', 0.55), ('RHTY', 0.35), ('RSTY', 0.26), ('RPAY', 0.25), ('RPSY', 0.24), ('RPTN', 0.11) | 749658 |
| **I358F 4th lib HSS** | 0.010 | 0.240 | 0.030 | 0.010 | 2.860 | 94.580 | ('YPTT', 0.39), ('RPSY', 0.18), ('RPAY', 0.18), ('RTTY', 0.17), ('LPTY', 0.15), ('RHTY', 0.13), ('RPTC', 0.12), ('RPTT', 0.11), ('RRTY', 0.11) | 756757 |
| **BA.1 4th lib HSS** | 0.130 | 0.090 | 0.000 | 0.000 | 0.000 | 97.920 | ('RPAY', 0.37), ('RPSY', 0.36) | 614899 |
| **BA.2 4th lib HSS** | 0.000 | 0.120 | 0.000 | 0.000 | 0.000 | 98.070 | ('RPSY', 0.34), ('RHTY', 0.32), ('RRTY', 0.16), ('RPAY', 0.14), ('RPTH', 0.14), ('RPTF', 0.13), ('RSTY', 0.12) | 817816 |

WT corresponds to the amino-acid in the isolate Wuhan-Hu-1 (MN908947.3) sequence and was used as a reference with a sequence 498-QPTN-501.

| Lineages | KD (nM) | 339 | 345 | 346 | 347 | 348 | 349 | 354 | 356 | 358 | 363 | 365 | 366 | 367 | 368 | 371 | 373 | 375 | 376 | 378 | 392 | 396 | 402 | 405 | 408 | 417 | 428 | 429 | 440 | 444 | 445 | 446 | 450 | 452 | 458 | 459 | 460 | 461 | 462 | 465 | 468 | 470 | 477 | 478 | 483 | 484 | 486 | 493 | 496 | 498 | 501 | 505 |
|---|---|---|---|---|---|---|---|---|---|---|---|---|---|---|---|---|---|---|---|---|---|---|---|---|---|---|---|---|---|---|---|---|---|---|---|---|---|---|---|---|---|---|---|---|---|---|---|---|---|---|---|---|
| WT aa | 8.5 (1) | G | T | R | F | A | P |  | K | I | Y | S | V | V | L | S | S | S | T | K | F | Y | I | D | R | K | D |  | N | K | V | G | N | L |  | S | N |  | K | E | I | T | S | T | V | E | F | Q | S | Q | N | Y |
| Beta | 4.1 (0.5) |  |  |  |  |  |  |  |  |  |  |  |  |  |  |  |  |  |  |  |  |  |  |  |  | N |  |  |  |  |  |  |  |  |  |  |  |  |  |  |  |  |  |  |  | K |  |  |  |  | Y |  |
| Gamma | 5.0 (0.5) |  |  |  |  |  |  |  |  |  |  |  |  |  |  |  |  |  |  |  |  |  |  |  |  | T |  |  |  |  |  |  |  |  |  |  |  |  |  |  |  |  |  |  |  | K |  |  |  |  | Y |  |
| BA.1 | 5.9 (0.4) | D |  |  |  |  |  |  |  |  |  |  |  |  |  | L | P | F |  |  |  |  |  |  |  | N |  |  | K |  |  | S |  |  |  |  |  |  |  |  |  |  | N | K |  | A |  | R | S | R | Y | H |
| BA.2 | 7.0 (0.6) | D |  |  |  |  |  |  |  |  |  |  |  |  |  | F | P | F | A |  |  |  |  | N | S | N |  |  | K |  |  |  |  |  |  |  |  |  |  |  |  |  | N | K |  | A |  | R |  | R | Y | H |
| BN.1 | 5.0 (0.6) | H |  | T |  |  |  |  | T |  |  |  |  |  |  | F | P | F | A |  |  |  |  | N | S | N |  |  | K | T |  |  |  |  |  |  | K |  |  |  |  |  | N | K |  | A |  |  |  | R | Y | H |
| BA.5 | 2.1 (0.1) | D |  |  |  |  |  |  |  |  |  |  |  |  |  | F | P | F | A |  |  |  |  | N | S | N |  |  | K |  |  |  |  | R |  |  |  |  |  |  |  |  | N | K |  | A | V |  |  | R | Y | H |
| XBB.1.5 | 1.2 (0.17) | H |  | T |  |  |  |  |  |  |  |  |  |  | I | F | P | F | A |  |  |  |  | N | S | N |  |  | K |  | P | S |  |  |  |  | K |  |  |  |  |  | N | K |  | A | P |  |  | R | Y | H |
| BA.1-SSS | 1.6 (0.15) | D |  |  |  |  |  |  |  |  |  |  |  |  |  | L | P | F |  |  |  |  |  |  |  | N |  |  | K |  |  | S |  |  |  |  |  |  |  |  |  |  | N | K |  | A |  | R | S | R | Y | H |
| **LSS** |  |  |  |  |  |  |  |  |  |  |  |  |  |  |  |  |  |  |  |  |  |  |  |  |  |  |  |  |  |  |  |  |  |  |  |  |  |  |  |  |  |  |  |  |  |  |  |  |  |  |  |  |
| WT-c71 | 0.9 (0.15) |  |  |  |  |  |  |  |  |  |  |  |  |  |  |  |  |  |  |  |  |  |  | E |  |  |  |  | K |  |  |  |  |  |  |  | K |  |  |  |  |  |  |  | A |  |  |  |  | H | S |  |
| WT-c72 | 4.7 (0.6) |  |  |  |  |  | P |  |  |  |  |  | T | A |  |  |  |  |  | R |  |  |  |  |  |  |  |  |  |  |  |  |  |  |  |  | K |  |  | G |  |  |  |  |  |  |  |  |  | H | Y |  |
| WT-c74 | 0.5 (0.25) |  |  |  |  |  |  | I |  |  |  |  |  |  |  |  |  |  |  |  |  |  |  |  |  |  |  |  |  | D |  |  |  |  |  |  |  |  |  |  |  |  |  |  |  |  |  |  |  |  |  | R |
| Alfa-c32 | 5.0 (2.0) |  |  |  |  |  |  |  |  |  |  |  |  |  |  |  |  |  |  |  |  |  |  |  |  |  |  |  |  |  |  |  |  |  |  |  |  |  |  |  |  |  |  |  |  |  |  |  |  |  |  | H |
| Alfa-c35 | 4.3 (1.1) |  |  |  |  |  |  |  |  |  |  |  |  |  |  |  |  |  |  |  |  |  |  |  |  |  |  |  |  |  |  |  |  |  |  |  |  |  |  |  |  |  |  |  |  |  |  |  |  |  |  | H |
| Alfa-c71 | 1.6 (0.25) |  |  |  |  |  |  |  |  |  |  | F |  |  |  |  |  |  |  |  |  |  |  |  |  |  |  |  |  |  |  |  |  |  |  |  |  |  |  |  |  |  |  |  |  |  |  |  |  |  | Y |  |
| Alfa-c76 | 0.6 (0.1) |  |  |  |  |  |  |  |  | F |  |  |  |  |  |  |  |  |  |  |  |  | V |  |  |  |  |  |  |  |  |  |  | Q |  |  |  |  |  |  |  |  | N |  |  |  |  |  |  |  | Y |  |
| Beta-c31 | 0.36 (0.08) |  |  |  |  |  |  |  |  |  |  |  |  |  |  |  |  |  |  |  |  |  |  |  |  |  | N |  |  |  |  |  |  |  |  |  |  |  |  |  |  |  |  |  |  |  |  | K |  |  |  | R |
| Beta-c55 | 0.27 (0.1) |  |  |  |  |  |  |  |  |  |  |  |  |  |  |  |  |  |  |  |  |  |  |  | I |  | I |  |  |  |  |  |  |  |  |  |  |  |  |  |  |  |  |  |  | H |  | K |  |  |  | R |
| Beta-c72 | 0.37 (0.05) |  |  |  |  |  |  |  | R |  |  |  |  |  |  |  |  |  |  |  |  |  |  |  |  | N |  |  | K |  |  |  |  |  |  |  | K |  |  |  |  |  |  |  |  | K |  |  |  | R | Y |  |
| Beta-c73 | 0.62 (0.1) |  |  |  |  |  |  |  |  |  |  |  |  |  |  |  |  |  |  | R |  |  |  |  |  | N |  |  |  |  |  |  |  |  |  | T | K |  |  |  |  |  |  |  |  | N |  |  |  | R | Y |  |
| v48-c72 | 2.2 (0.3) |  |  |  | S | P |  |  |  |  |  |  |  |  |  |  |  |  | S |  | L | N |  |  |  | N |  |  |  |  |  |  |  |  |  |  |  |  | E |  | V |  |  |  |  | K |  |  |  | R | Y |  |
| v48-c71 | 2.6 (0.25) |  |  |  |  |  |  |  |  |  | P |  |  |  |  |  |  |  |  |  |  |  |  |  |  | N |  |  |  |  |  |  |  |  |  |  | S |  |  |  | V |  |  |  |  | K |  |  |  | R | Y |  |
| v48-c43 | 0.4 (0.08) |  |  |  |  |  |  |  |  |  |  |  |  |  |  |  | L |  |  |  |  |  |  |  |  |  | N |  |  |  |  |  |  |  |  |  |  |  |  |  |  |  |  |  |  |  |  | K |  |  |  | R |
| v48-cn61 | 0.37 (0.1) |  |  |  |  |  |  |  |  |  |  |  |  |  |  |  |  |  |  |  |  |  |  |  |  |  |  |  |  |  |  |  |  |  |  |  |  |  |  |  |  |  |  |  |  |  |  |  |  |  |  |  |
| BA.1-c72 | 0.4 (0.09) |  | N |  |  |  |  |  |  |  |  |  |  |  |  | L | P |  |  |  |  |  |  |  |  | N | G |  | K |  |  |  |  |  |  |  |  |  |  |  |  | A | N | K |  | A |  |  | S | R | Y | H |
| **HSS** |  |  |  |  |  |  |  |  |  |  |  |  |  |  |  |  |  |  |  |  |  |  |  |  |  |  |  |  |  |  |  |  |  |  |  |  |  |  |  |  |  |  |  |  |  |  |  |  |  |  |  |  |
| WT-c41 | 1.6 (0.1) |  |  |  |  |  |  |  |  |  |  |  |  |  |  |  |  |  |  |  |  |  |  |  |  |  |  |  |  |  |  |  | D | M |  |  | K |  |  |  |  |  | N | K |  | R |  | R |  | R | Y |  |
| WT-c42 | 0.6 (0.05) |  |  |  |  |  |  |  |  |  |  |  |  |  |  |  |  |  |  |  |  |  |  |  |  |  |  |  |  |  |  |  | D | M |  |  | K |  |  |  |  |  | N | K |  | R |  |  |  | R | Y |  |
| WT-c43 | 0.5 (0.05) |  |  |  |  |  |  |  |  |  |  |  |  |  |  |  |  |  |  |  |  |  |  |  |  |  |  |  | K | R |  |  | D | M |  |  | K |  |  |  |  |  | N | K |  | R |  |  |  | R | Y |  |
| I358F-c41 | 0.6 (0.06) |  |  |  |  |  |  |  |  | F |  |  |  |  |  |  |  |  |  |  |  |  |  |  |  |  |  |  | K | T |  |  |  | R |  |  | K |  |  |  |  |  | N | K |  | A |  | R |  | R | Y |  |
| I358F-c42 | 0.5 (0.05) |  |  |  |  |  |  |  |  | F |  |  |  |  |  |  |  |  |  |  |  |  |  |  |  |  |  |  | K | T |  |  |  | R |  |  | K |  |  |  |  |  | N | K |  | R |  | R |  | R | Y |  |
| I358F-c43 | 0.73 (0.06) |  |  |  |  |  |  |  |  | F |  |  |  |  |  |  |  |  |  |  |  |  |  |  |  |  |  |  | K | T |  |  |  | R |  |  | K |  |  |  |  |  | N | K |  | K |  |  |  | R | Y |  |
| BA.1-c41 | 1.7 (0.15) | D |  |  |  |  |  |  |  |  |  |  |  |  |  | L | P | F |  |  |  |  |  |  |  | N |  |  | K |  |  | S | D | M |  |  | K |  |  |  |  |  | N | K |  | R |  |  |  | R | Y | H |
| BA.1-c42 | 1.7 (0.15) | D |  |  |  |  |  |  |  |  |  |  |  |  |  | L | P | F |  |  |  |  |  |  |  | N |  |  | K |  |  | S | D | M |  |  | K |  |  |  |  |  | N | K |  | R |  |  |  | R | Y |  |
| BA.1-c43 | 1.7 (0.15) | D |  |  |  |  |  |  |  |  |  |  |  |  |  | L | P | F |  |  |  |  |  |  |  | N |  |  | K | A |  |  |  | R |  |  | K |  |  |  |  |  | N | K |  | R |  |  |  | R | Y | H |
| BA.2-c41 | 0.86 (0.1) | D |  |  |  |  |  |  |  |  |  |  |  |  |  | F | P | F | A |  |  |  |  | N | S | N |  |  | K | R |  |  | D | K |  |  | K |  |  |  |  |  | N | K |  | R |  |  |  | R | Y | H |
| BA.2-c42 | 2.5 (0.25) | D |  |  |  |  |  |  |  |  |  |  |  |  |  | F | P | F | A |  |  |  |  | N | S | N |  |  | K | R |  |  | D | M |  |  | K |  |  |  |  |  | N | K |  | R |  |  |  | R | Y | H |

**Table S7. Binding affinities and sequences of clones from Fig. 4.** Binding affinities were determined from at least 3 independent replicates using GraphPad v.10 and one site specific binding model. In parenthesis are ± 95% CI values. All mutations relative to WT, as determined by Sanger sequencing of the selected clones are shown. For HSS selected clones, residues in gray are those in the stability determining region, which were not mutated.

#### Supporting information part PS2 – Analysis of error prone libraries for high stringency selections

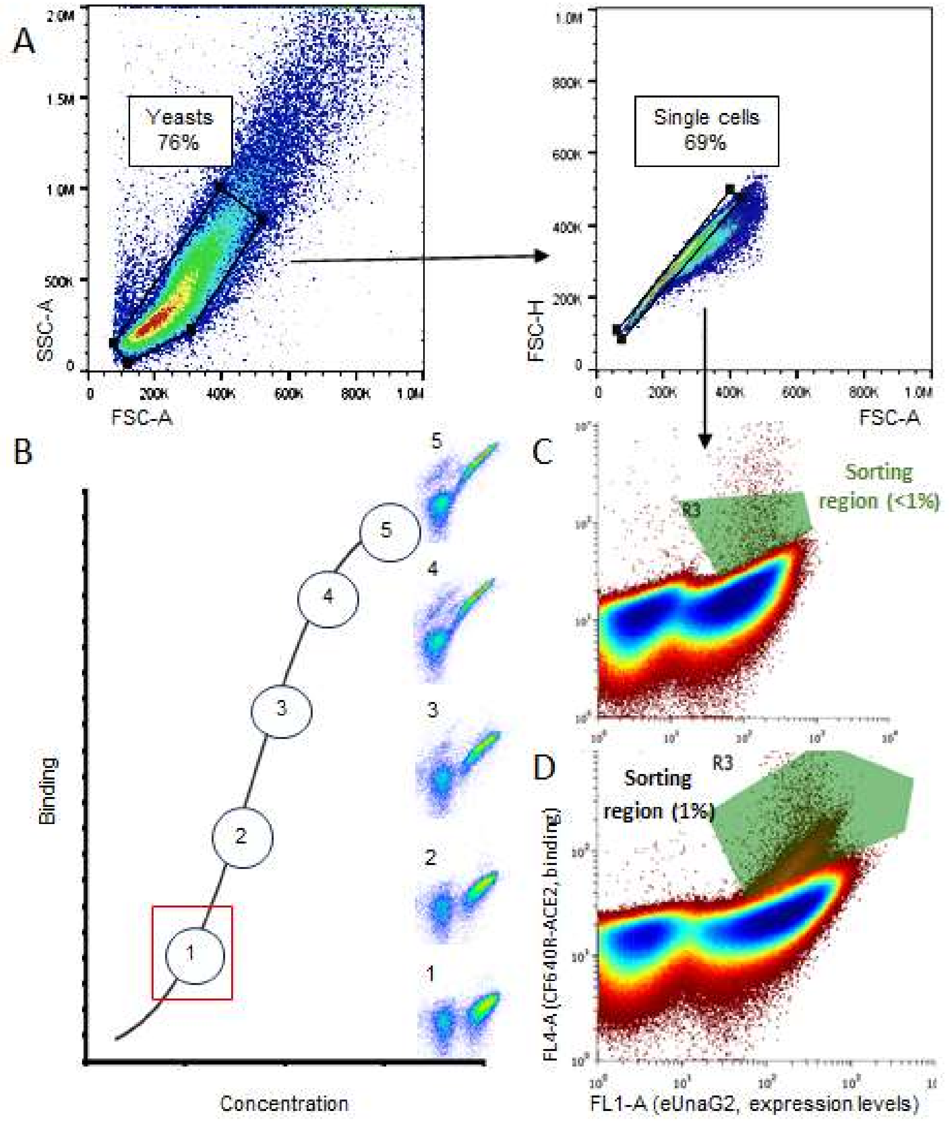

**Fig. S1 – Gating and selection strategies for in vitro evolution of the SARS-CoV-2 RBD domain.** (A) Gating strategy used during FACS sorting. Yeast cells were selected based on their forward and side scatter area parameters (FSC-A and SSC-A) and subsequently, single cells were gated using a diagonal plot of FCS-A versus height (FCS-H). (B) Sorting under different ACE-2 concentrations. The library was incubated with a range of ACE2 concentrations to identify the minimal concentration that promotes elevated signal (inset 1-3). Under these conditions, clones with stronger binding affinity had the highest competitive/signal advantage over the parental population. For LSS higher concentration conditions were used (insets 4–5). (C) Stepwise sorting strategy for HSS to enhance discrimination between improved clones and the parental population. Initially, below 1% of top signal cells were sorted in HSS. These selected cells were then cultured, expressed and subjected to a second round of sorting for higher enrichment of the enhanced binding population (D).

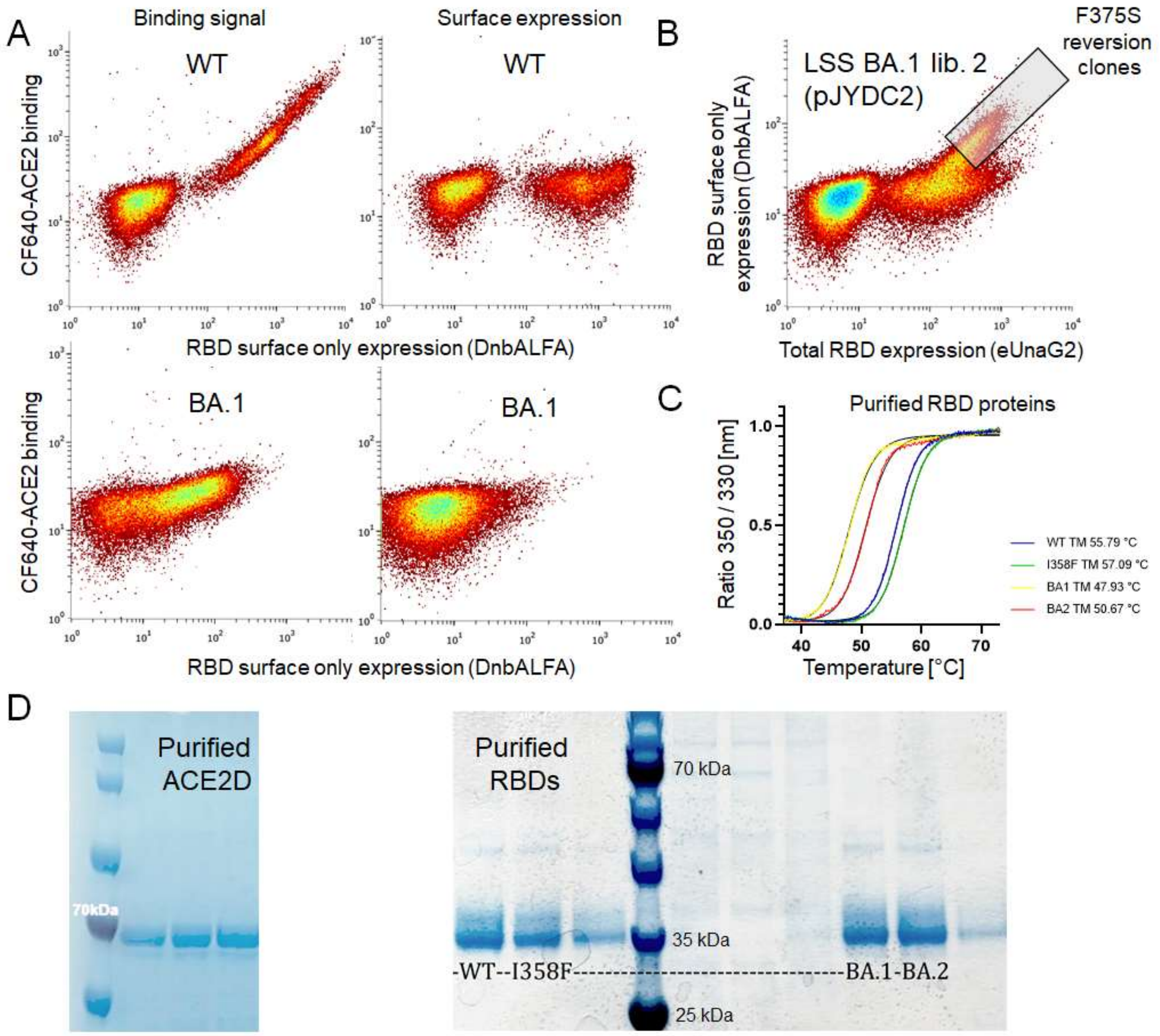

**Fig. S2 – Yeast surface display and stability analysis of purified proteins.** (A) Flow cytometry dot plot showing strong ACE2 binding and robust surface expression for the wild-type (WT) RBD, in contrast to markedly reduced binding and surface expression observed for the BA.1 RBD. (B) A dominant population displaying a reverted phenotype, relative to BA.1 in panel A, dominated in the population as early as the second round of selection. Surface versus total expression was assessed using the pJYDC2 plasmid. Sequencing identified the F375S reversion as responsible for the restored phenotype. (C) Tycho NT.6 (NanoTemper) thermal unfolding profiles of purified WT, WT+I358F, BA.1, and BA.2 RBDs, demonstrating substantial destabilization of both Omicron lineages relative to WT. (D) SDS-PAGE analysis of purified ACE2D and RBDs proteins with marker (BlueRay Prestained Protein Marker).

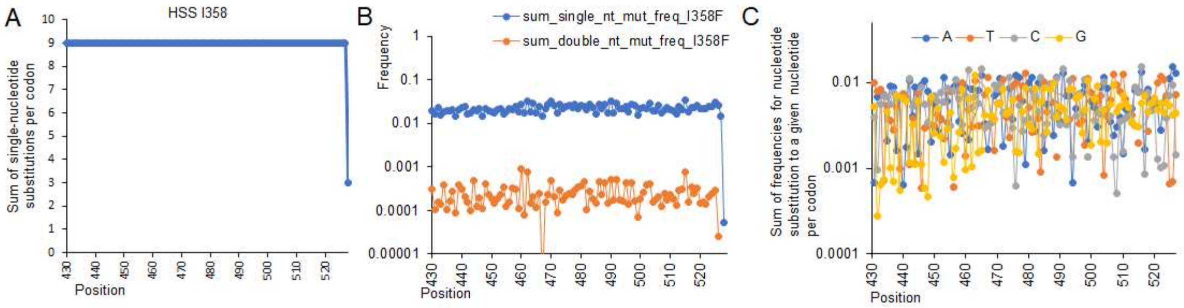

**Fig. S3 – WT+I358F library quality assessment.** (A) The number of observed single nucleotide substitutions per codon along the RBM, with 9 representing all mutations in all three positions. (B) Frequencies for all single- and double-nucleotide substitutions per codon, showing minimal positional deviations. (C) Sum of single-nucleotide mutation frequencies per mutant nucleotide and codon.

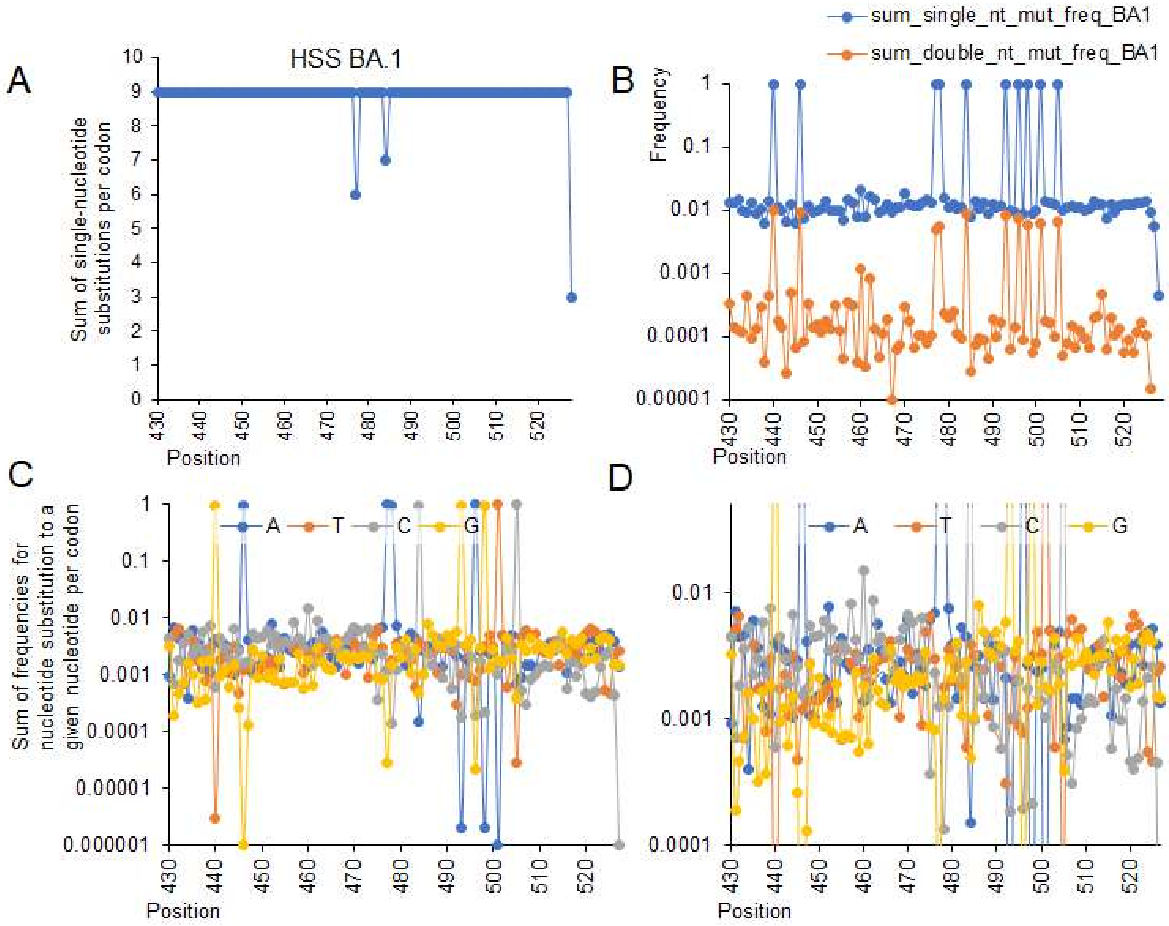

**Fig. S4 – BA.1 library quality assessment.** (A) The number of observed single nucleotide substitutions per codon along the RBM, with 9 representing all mutations in all three positions. (B) Frequencies for all single- and double-nucleotide substitutions per codon, showing minimal positional deviations. Note that the high frequencies is certain positions are of the BA.1 mutations. (C) Sum of single-nucleotide mutation frequencies per mutant nucleotide and codon with WT reference sequence. (D) Sum of single-nucleotide mutation frequencies per mutant nucleotide and codon – detail for frequencies from 0.001 to 0.01.

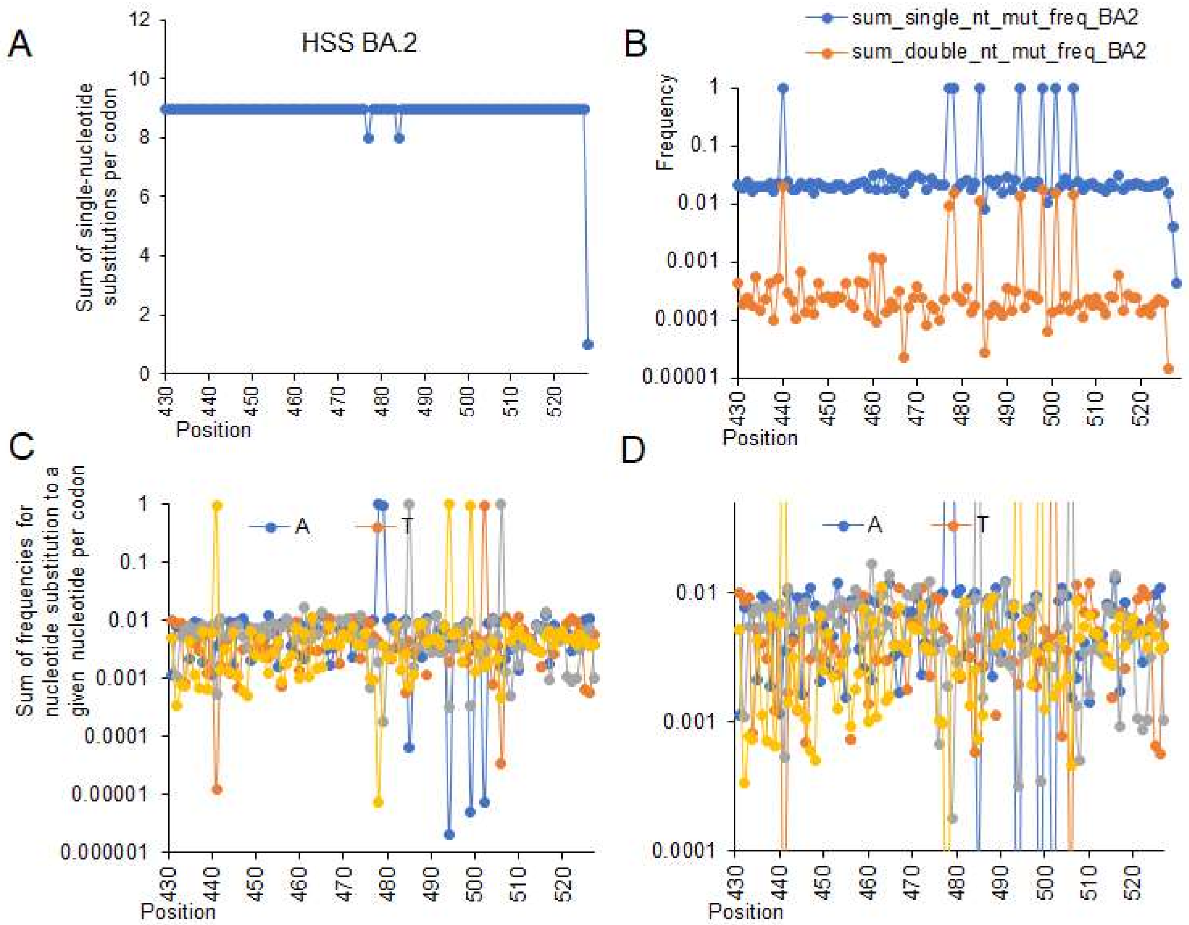

**Fig. S5 – BA.2 library quality assessment.** (A) The number of observed single nucleotide substitutions per codon along the RBM, with 9 representing all mutations in all three positions. (B) Frequencies for all single- and double-nucleotide substitutions per codon, showing minimal positional deviations. (C) Sum of single-nucleotide mutation frequencies per mutant nucleotide and codon with WT reference sequence. (D) Sum of single-nucleotide mutation frequencies per mutant nucleotide and codon – detail for frequencies from 0.001 to 0.01.

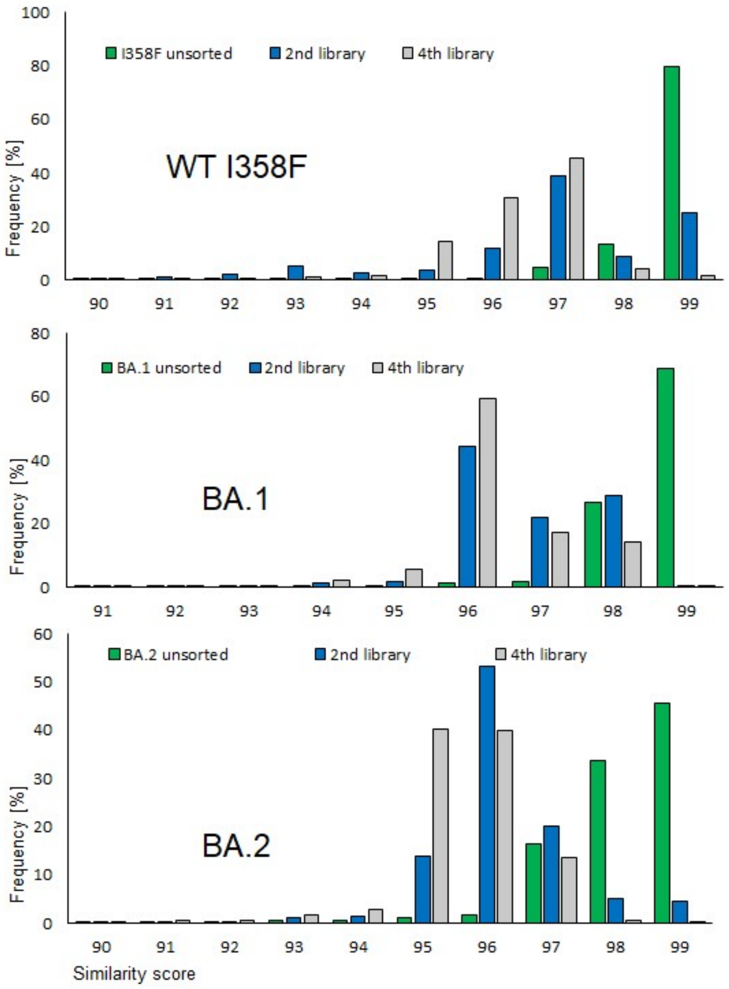

**Fig. S6 – Comparison of similarity scores for individual sequencing reads in HSS libraries.** Merged paired-end reads into a single sequence with WT, BA.1 and BA.2 sequences as the reference for the unsorted initial library. Populations after sorting in the 2nd and 4th rounds of the in vitro evolution are shown. The sequenced regions and corresponding merged paired-end read length for individual libraries differ, and therefore, similarity scores cannot be compared across libraries.

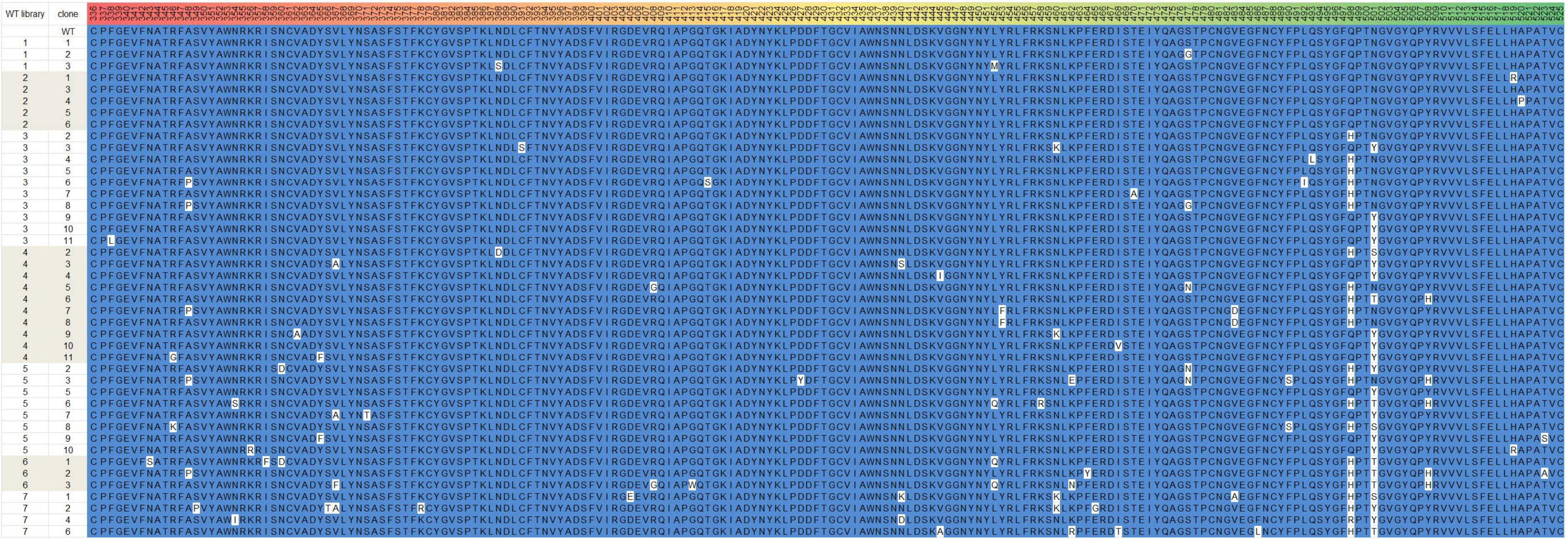

**Fig. S7 – Translated Sanger sequencing results for selected clones for WT LSS libraries 1 to 7**

| Alfa library | clone | |
|---|---|---|
| | WT | CPFGEVFNATRFASVYAWNRKRISNCVADYSVLYNSASFSTFKCYGVSPTKLNDLCFTNVYADSFVIRGDEVRQIAPGQTGKIADYNYKLPDDFTGCVIAWNSNNLDSKVGGNYNYLYRLFRKSNLKPFERDISTEIYQAGSTPCNGVEGFNCYFPLQSYGFQPTNGVGYQPYRVVVLSFELLHAPATVC |
| | Alfa | CPFGEVFNATRFASVYAWNRKRISNCVADYSVLYNSASFSTFKCYGVSPTKLNDLCFTNVYADSFVIRGDEVRQIAPGQTGKIADYNYKLPDDFTGCVIAWNSNNLDSKVGGNYNYLYRLFRKSNLKPFERDISTEIYQAGSTPCNGVEGFNCYFPLQSYGFQPTYGVGYQPYRVVVLSFELLHAPATVC |
| 1 | 1 | CPFGEVFNATRFASVYAWNRKRISNCVADYSVLYNSASFSTFKCYGVSPTKLNDLCFTNVYADSFVIRGDEVRQIAPGQTGKIADYNYKLPDDFTGCVIAWNSNNLDSKVGGNYNYLYRLFRKSNLKPFERDISTEIYQAGSTPCNGVEGFNCYFPLQSYGFQPTYGVGYQPYRVVVLSFELLHAPATVC |
| 1 | 2 | CPFGEVFNATRFASVYAWNRKRISNCVADYSVLYNSASFSTFKCYGVSPTKLNDLCFTNVYADSFVIRGDEVRQIAPGQTGKIADYNYKLPDDFTGCVIAWNSNNLDSKVGGNYNYLYRLFRKSNLKPFERDISTEIYQAGSTPCNGVEGFNCYFPLQSYGFQPTYGVGYQPYRVVVLSFELLHAPATVC |
| 1 | 3 | CPFGEVFNATRFASVYAWNRKRISNCVADYSVLYNSASFSTFKCYGVSPTKLNDLCFTNVYADSFVIRGDEVRQIAPGQTGKIADYNYKLPDDFTGCVIAWNSNNLDSKVGGNYNYLYRLFMKSNLKPFERDISTEIYQAGSTPCNGVEGFNCYFPLQSYGFQPTYGVGYQPYRVVVLSFELLHAPATVC |
| 2 | 1 | CPFGEVFNATRFASVYAWNRKRISNCVADYSVLYNSASFSTFKCYGVSPTKLNDLCFTNVYADSFVIRGDEVRQIAPGQTGKIADYNYKLPDDFTGCVIAWNSNNLDSKVGGNYNYLYRLFRKSNLKPFERDISTEIYQAGSTPCNGVEGFNCYFPLQSYGFQPTYGVGYQPYRVVVLSFELLHAPATVC |
| 2 | 2 | CPFGEVFNATRFASVYAWNRKRISNCVADYSVLYNSASFSTFKCYGVSPTKLNDLCFTNVYADSFVIRGDEVRQIAPGQTGKIADYNYKLPDDFTGCVIAWNSNNLDSKVGGNYNYLYRLFRKSNLKPFERDISTEIYQAGSTPCNGVKGFNCYFPLQSYGFQPTYGVGYQPYRVVVLSFELLHAPATVC |
| 2 | 3 | CPFGEVFNATRFASVYAWNRKRISNCVADYSVLYNSASFSTFKCYGVSPTKLNDLCFTNVYADSFVIRGDEVRQIAPGQTGKIADYNYKLPDDFTGCVIAWNSNNLDSKVGGNYNYLYRLFRKSNLKPFERDISTEIYQAGSTPCNGVEGFNCYFPLQSYGFQPTYGVGYQPYRVVVLSFELLHAPATVC |
| 3 | 5 | CPFGEVFNATRFASVYAWNRKRISNCVADYSVLYNSASFSTFKCYGVSPTKLNDLCFTNVYADSFVIRGDEVRQIAPGQTGKIADYNYKLPDDFTGCVIAWNSNNLDSKVGGNYNYLYRLFRKSNLKPFERDISTEIYQAGSTPCNGVEGFNCYFPLQSYGFHPTNGVGYQPYRVVVLSFELLHAPATVC |
| 3 | 2 | CPFGEVFNATRFASVYAWNRKRISNCVADYSVLYNSASFSTFKCYGVSPTKLNDLCFTNVYADSFVIRGDEVRQIAPGQTGKIADYNYKLPDDFTGCVIAWNSNNLDSKVGGNYNYLYRLFRKSNLKPFERDISTEIYQAGSTPCNGVEGFNCYFPLQSYGFHPTNGVGYQPYRVVVLSFELLHAPATVC |
| 3 | 9 | CPFGEVFNATRFASVYAWNRKRISNCVADYSVLYNSASFSTFKCYGVSPTKLNDLCFTNVYADSFVIRGDEVRQIAPGQTGKIADYNYKLPDDFTGCVIAWNSNNLDSKVGGNYNYLYRLFRKSNLKPFERDISTEIYQAGNTPCNGVEGFNCYFPLQSYGFHPTNGVGYQPYRVVVLSFELLHAPATVC |
| 4 | 1 | CPFGEVFNATRFASVYAWNRKRISNCVADYSVLYNSASFSTFKCYGVSPTKLNDLCFTNVYADSFVVRGDEVRQIAPGQTGKIADYNYKLPDDFTGCVIAWNSNNLDSKVGGNYNYQYRLFRKSNLKPFERDISTEIYQAGSTPCNGVEGFNCYFPLQSYGFQPTYGVGYQPYRVVVLSFELLHAPATVC |
| 4 | 4 | CPFGEVFNATRFASVYAWNRKRISNCVADYSVLYNSASFSTFKCYGVSPTKLNDLCFTNVYADSFVIRGDEVRQIAPGQTGKIADYNYKLPDDFTGCVIAWNSNNLDSKVGGNYNYLYRLFRKSNLKPFERDISTEIYQAGNTPCNGVEGFNCYFPLQSYGFHPTNGVGYQPYRVVVLSFELLHAPATVC |
| 4 | 5 | CPFGEVFNATRFASVYAWNRKRISNCVADYSVLYNSASFSTFKCYGVSPTKLNDLCFTNVYADSFVVRGDEVRQIAPGQTGKIADYNYKLPDDFTGCVIAWNSNNLDSKVGGNYNYLYRLFRKSNLKPFERDISTEIYQAGSTPCNGVEGFNCYFPLQSYGFQPTYGVGYQPYRVVVLSFELLHAPATVC |
| 4 | 6 | CPFGEVFNATRFASVYAWNRKRISNCVADYSFLYNSASFSTFKCYGVSPTKLNDLCFTNVYADSFVIRGDEVRQIAPGQTGKIADYNYKLPDDFTGCVIAWNSNNLDSKVGGNYNYLYRLFRKSNLKPFERDISTEIYQAGSTPCNGVEGFNCYFPLQSYGFRPTYGVGYQPYRVVVLSFELLQAPATVC |
| 4 | 7 | CPFGEVFNATRFASVYAWNRKRISNCVADYSVLYNSASFSTFKCYGVSPTKLNDLCFTNVYADSFVIRGDEVRQIAPGQTGKIADYNYKLPDDFTGCVIAWNSNNLDSKVGGNYNYLYRLFRKSNLKPFERDISTEIYQAGSTPCNGVEGFNCYFPLQSYGFQPTYGVGYQPYRVVVLSFELLHAPATVC |
| 4 | 9 | CPFGEVFNATRFASVYAWNRKRISNCVADYSVLYNSASFSTFKCYGVSPTKLNDLCFTNVYADSFVIRGDEVRQIAPGQTGKIADYNYKLPDDFTGCVIAWNSNNLDSKVGGNYNYLYRLFRESNLKPFERDISTEIYQAGSTPCNGVEGFNCYFPLQSYGFQPTYGVGYQPYRVVVLSFELLHAPATVC |
| 4 | 10 | CPFGEVFNATRFASVYAWNRKRISNCVADYSVLYNSASFSTFKCYGVSPTKLNDLCFTNVYADSFVIRGDEVRQIAPGQTGKIADYNYKLPDDFTGCVIAWNSNNLDSKVGGNYNYLYRLFRKSNLKPFERDISTEIYQAGSTPCNGVEGFNCYFPLQSYGFRPTYGVGYQPYRVVVLSFELLHAPATVC |
| 4 | 11 | CPFGEVFNATRFASVYAWNRKRISNCVADYSVLYNSASFSTFKCYGVSPTKLNDLCFTNVYADSFVIRGDEVRQIAPGQTGKIADYNYKLPDDFTGCVIAWNSNNLDSKVGGNYNYQYRLFRKSNLKPFERDISTEIYQAGSTPCNGVEGFNCYFPLQSYGFQPTYGVGYQPYRVVVLSFELLHAPATVC |
| 4 | 12 | CPFGEVFNATRFASVYAWNRKRISNCVADYSVLYNSASFSTFKCYGVSPTKLNDLCFTNVYADSFVIRGDEVRQIAPGQTGKIADYNYKLPDDFTGCVIAWNSNNLDSKVGGNYNYLYRLFRKSNLKPFERDTSTEIYQAGSTPCNGVEGFNCYFPLQSYGFQPTYGVGYQPYRVVVLSFELLHAPATVC |
| 4 | 13 | CPFGEVFNATRFASVYAWNRKRISNCVADYSVLYNSASFSTFKCYGVSPTKLNDLCFTNVYADSFVIRGDEVRQIAPGQTGKIADYNYKLPDDFTGCVIAWNSNNLDSKVGGNYNYQYRLFRKSNLKPFERDISTEIYQAGSTPCNGVEGFNCYFPLLSYGFHPTNGVGYQPYRVVVLSFELLHAPATVC |
| 5 | 2 | CPFGEVFNATRFASVYAWNRKRISNCVADYSVLYNSASFSTFKCYGVSPTKLNDLCFTNVYADSFVIRGDEVRQIAPGQTGKIADYNYKLPDDFTGCVIAWNSNNLDSKVGGNYNYLYRLFRKSNLKPFERDISTEIYQAGGSPCNGVEGFNCYFPLQSYGFQPTYGVGYQPYRVVVLSFELLHAPATVC |
| 5 | 5 | CPFGEVFNATRFASVYAWNRKRISNCVADYSALYNSASFSTFKCYGVSPTKLNDLCFTNVYADSFVIRGDEVRQIAPGQTGKIADYNYKLPDDFTGCVIAWNSNNLDSKVGGNYNYQYRLFRKSNLKPFERDISTEIYQAGSTPCNGVEGFNCYFPLQSYGFQPTYGVGYQPYRVVVLSFELLHAPATVC |
| 5 | 6 | CPFGEVFNATRFASVYAWNRKRISNCVADYSVLYNSASFSTFKCYGVSPTKLNDLCFTNVYADSFVIRGDEVRQIAPGQTGKIADYNYKLPDDFTGCVIAWNSNNLDSKVGGNYNYLYRLFRKSNLKPFERDISTEIYQAGSTPCNGVEGFNCYFPLQSYGFQPTYGVGYQPYRVVVLSFELLHAPATVC |
| 5 | 7 | CPFGEVFNATRFASVYAWNRKRISNCVADYSALYNSASFSTFKCYGVSPTKLNDLCFTNVYADSFVIRGDEVRQIAPGQTGKIADYNYKLPDDFTGCVIAWNSNNLDSKVGGNYNYLYRLFRKSNLKPFERDISTEIYQAGNTPCNGVEGFNCYFPLQSYGFHPTNGVGYQPYRVVVLSFELLHAPATVC |
| 5 | 8 | CPFGEVFNATRFASVYAWNRKRISNCVADYSVLYNSASFSTFKCYGVSPTKLNDLCFTNVYADSFVIRGDEVRQIAPGQTGKIADYNYKLPDDFTGCVIAWNSNNLDSKVGGNYNYLFRLFRKSNLKPFERDTSTEIYQAGSTPCNGVEGFNCYFPLQSYGFQPTYGVGYQPYRVVVLSFELLHAPATVC |
| 5 | 9 | CPFGEVFNATRFASVYAWNRKRFSNCVADYSVLYNSASFSTFKCYGVSPTKLNDLCFTNVYADSFVIRGDEVRQIAPGQTGKIADYNYKLPDDFTGCVIAWNSNNLDSKVGGNYNYLYRLFRKSNLKPFERDISTKIYQAGSTPCNGVEGFNCYFPLQSYGFQPTYGVGYQPYRVVVLSFELLHAPATVC |
| 5 | 10 | CPFGEVFNATRFASVYAWNRKRISNCVADYSVLYNSASFSTFKCYGVSPTKLNDLCFTNVYADSFVISGDEVRQIAPGQTGKIADYNYKLPDDFTGCVIAWNSNNLDSKVGGNYNYQYRLFRKSNLKPFERDISTEIYQAGSTPCEGVEGFNCYFPLQSYGFQPTYGVGYQPYRVVVLSFELLHAPATVC |
| 5 | 11 | CPFGEVFNATRFASVYAWNRKRISNCVADYSVLYNSASFSTFKCYGVSPTKLNDLCFTNVYADSFVIRGDEVRQIAPGQTGKIADYNYKLPDDFTGCVIAWNSNNLDSKVGGNYNYQYRLFRKSNLKPFERDISTEIYQAGSTPCNGVEGFNCYFPLQSYGFQPTYGVGYQPYRVVVLSFELLHAPATVC |
| 6 | 1 | CPLGEVFNATRFASVYAWNRKRISNCVADYSVLYNSASFSTFKCYGVSPTKLNDLCFTNVYADSFVIRGDEVRQIAPGQTGKIADYNYKLPDDFTGCVIAWNSNNLDSKVGGNYNYLFRLFRKSNLKPFERDISTEIYQAGSTPCNGVEGFNCYYPLQSYGFHPTNGVGYQPYRVVVLSFELLHPPATVC |
| 6 | 2 | CPFGEVFNATRFASVYAWNRKRISNCVADYSVLYNSASFSTFKCYGVSPTKLNDLCFTNVYADSFVIRGDEVRQIAPGQTGKIADYNYKLPDDFTGCVIAWNSNNLDSKVGGNYNYQYRLFRKSNLKPFERDISTEIYQAGSTPCNGVEGFNCYFPLQSYGFQPTYGVGYQPYRVVVLSFELLHAPATVC |
| 6 | 4 | CPFGEVFNATRFASVYAWNRKRISNCVADYSVLYNSASFSTFKCYGVSPTKLNDLCFTNVYADSFVVRGDEVRQIAPGQTGKIADYNYKLPDDFTGCVIAWNSNNLDSKVGGNYNYLYRLFRKSNLKPFERDISTEIYQAGSTPCNGVEGFNCYFPLQSYGFRPTYGVGYQPYRVVVLSFELLHAPATVC |
| 7 | 1 | CPFGEVFNATRFASVYAWNRKRISNCVADFSVLYNSASFSTFKCYGVSPTKLNDLCFTNVYADSFVIRGDEVRQIAPGQTGKIADYNYKLPDDFTGCVIAWNSNNLDSKVGGNYNYLYRLFRKSNLKPFERDISTEIYQAGSTPCNGVEGFNCYFPLQSYGFQPTYGVGYQPYRVVVLSFELLLAPATVC |
| 7 | 2 | CPFSEVFNATRFASVYAWNRMRISNCVADYSVLYNSASFSTFKCYGVSPTKLNDLCFTNVYADSFVVRGDEVRQIAPGQTGKIADYNYKLPDDFTGCVIAWNSNNLDSKVGGNYNYQYRLFRKSNLKPFERDISTEIYQAGNTPCNGVEGFNCYFPLQSYGFHPTNGVGYQPYRVVVLSFELLHAPATVC |
| 7 | 6 | CPFGEVFNATRFASVYAWNRKRFSNCVADYSVLYNSASFSTFKCYGVSPTKLNDLCFTNVYADSFVVRGDEVRQIAPGQTGKIADYNYKLPDDFTGCVIAWNSNNLDSKVGGNYNYQYRLFRKSNLKPFERDISTEIYQAGNTPCNGVEGFNCYFPLQSYGFQPTYGVGYQPYRVVVLSFELLHPPATVC |

**Fig. S8 – Translated Sanger sequencing results for selected clones for Alpha LSS libraries 1 to 7**

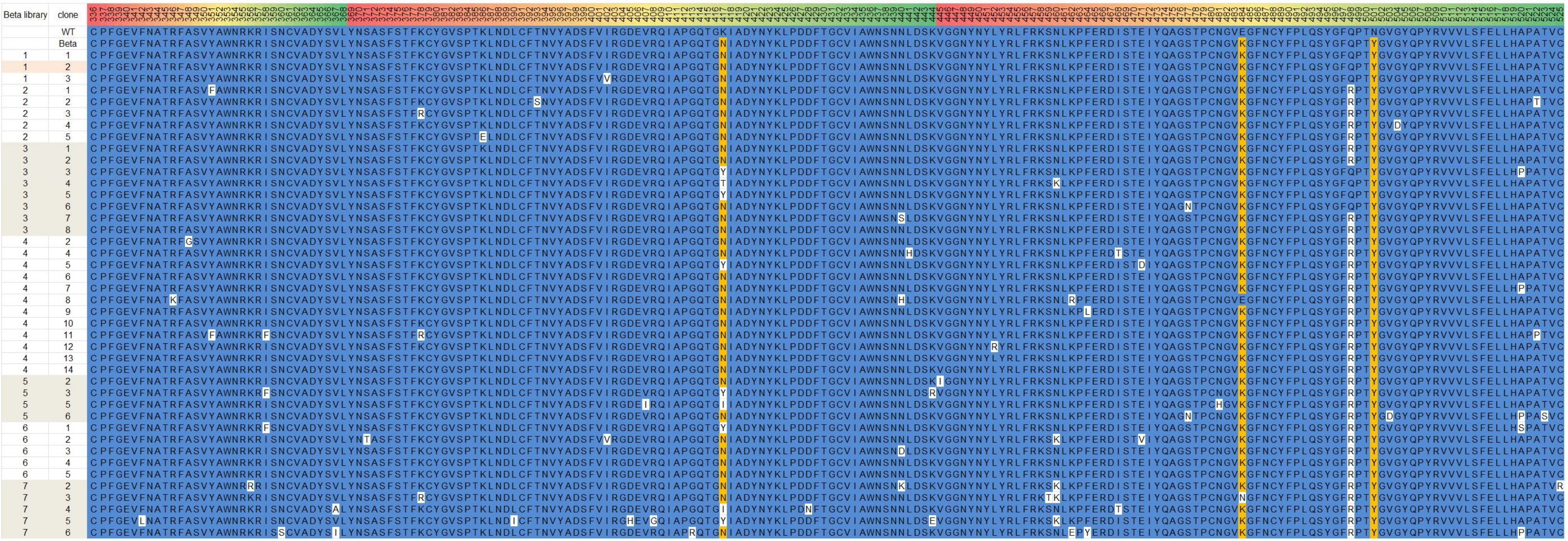

**Fig. S9 – Translated Sanger sequencing results for selected clones for Beta LSS libraries 1 to 7**

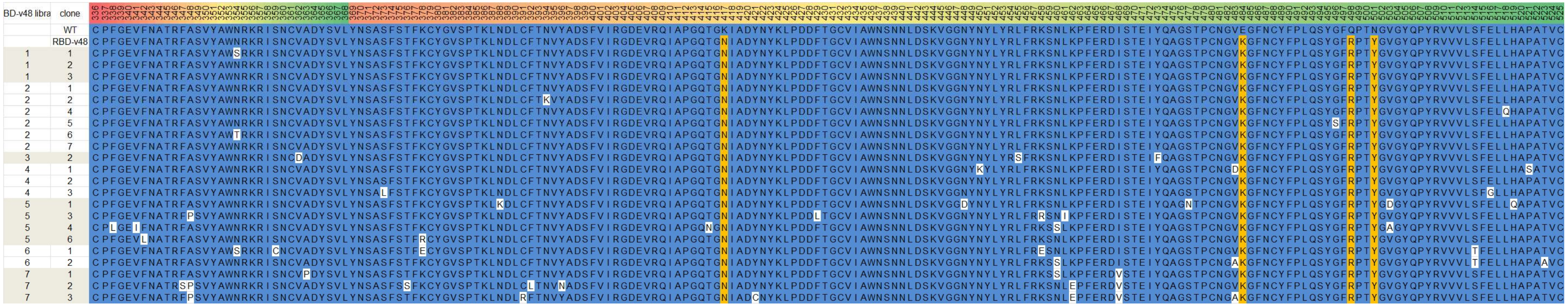

**Fig. S10 – Translated Sanger sequencing results for selected clones for RBD-v48 LSS libraries 1 to 7**

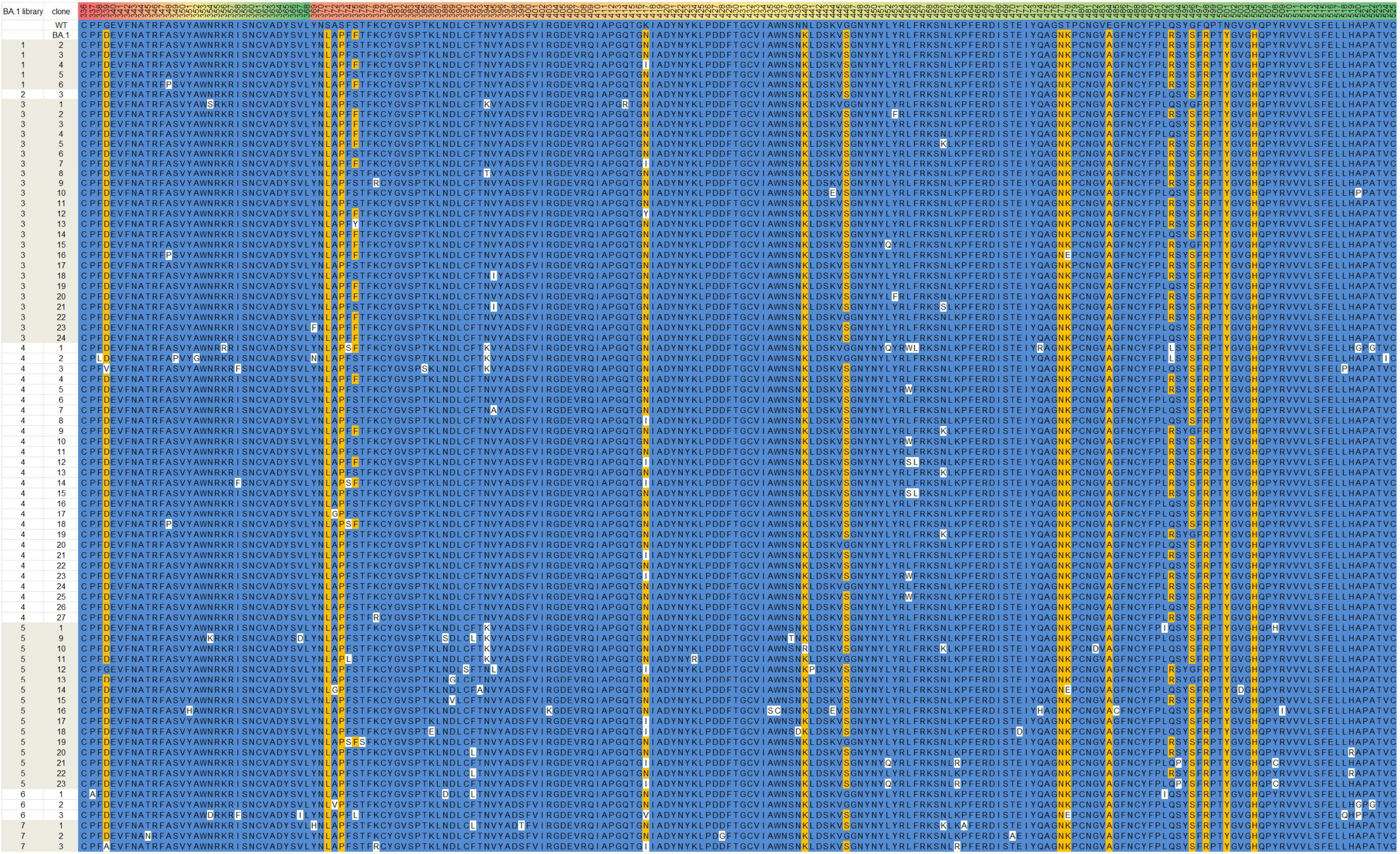

**Fig. S11 – Translated Sanger sequencing results for selected clones for BA.1 LSS libraries 1 to 7**

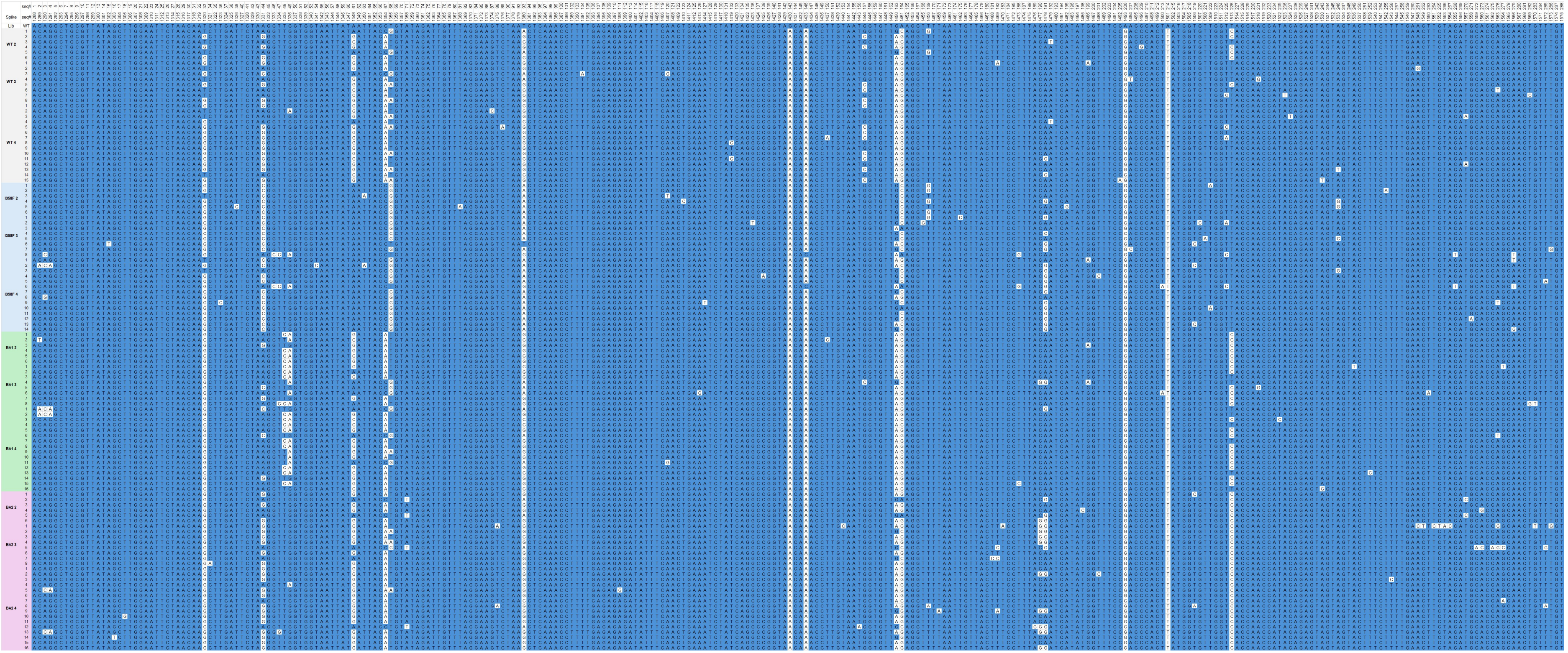

**Fig. S12 – Sanger sequencing results (nucleotides) for HSS libraries**

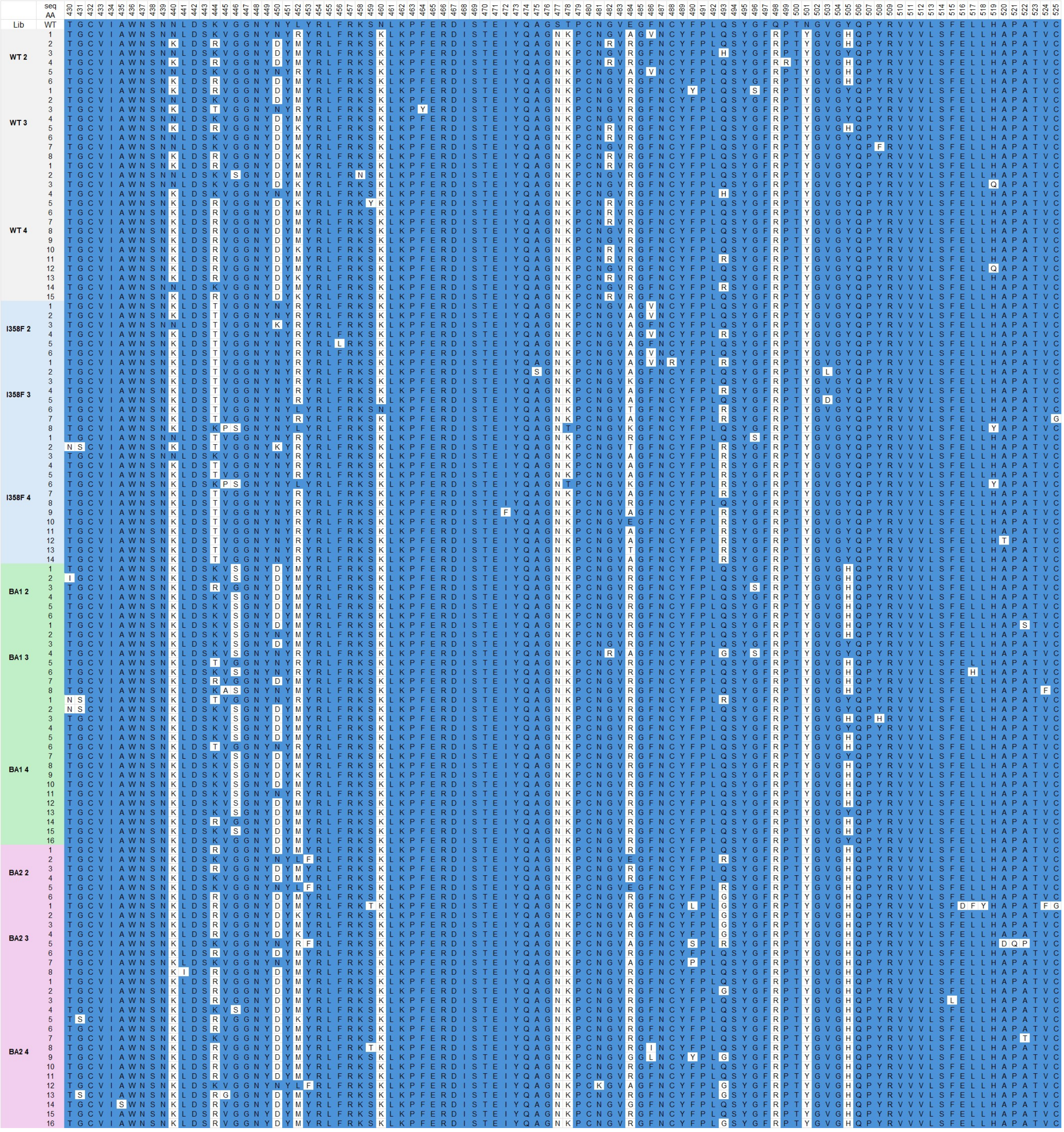

**Fig. S13 – Translated Sanger sequencing results for selected clones from HSS libraries**

#### Supporting information part PS3 – Analysis of mutations in low stringency selecti on libraries

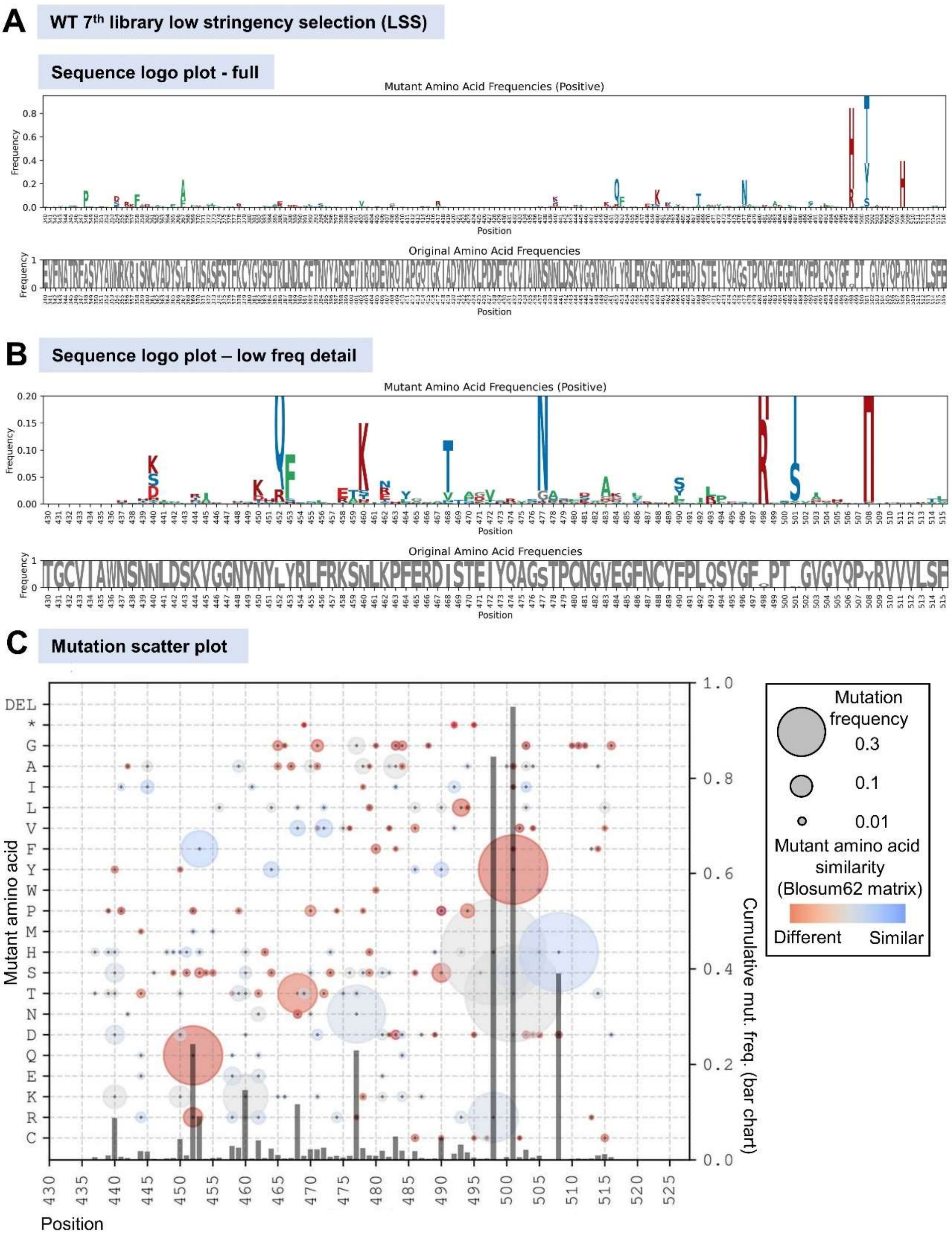

**Fig. S14 – Logo plots and mutation scatter plot for WT 7th library under LSS.** (A) Sequence logo plot (top panel) showing mutations in the library relative to the WT sequence. The bottom panel (in gray) displays the complementary frequency of the original amino acid at each position. (B) Sequence logo plot focusing on less frequent mutations in the library, with the y-axis frequency range set to 0–0.2. (C) Mutation scatter plot illustrating mutations in the population and their evolutionary distance from the original residues.

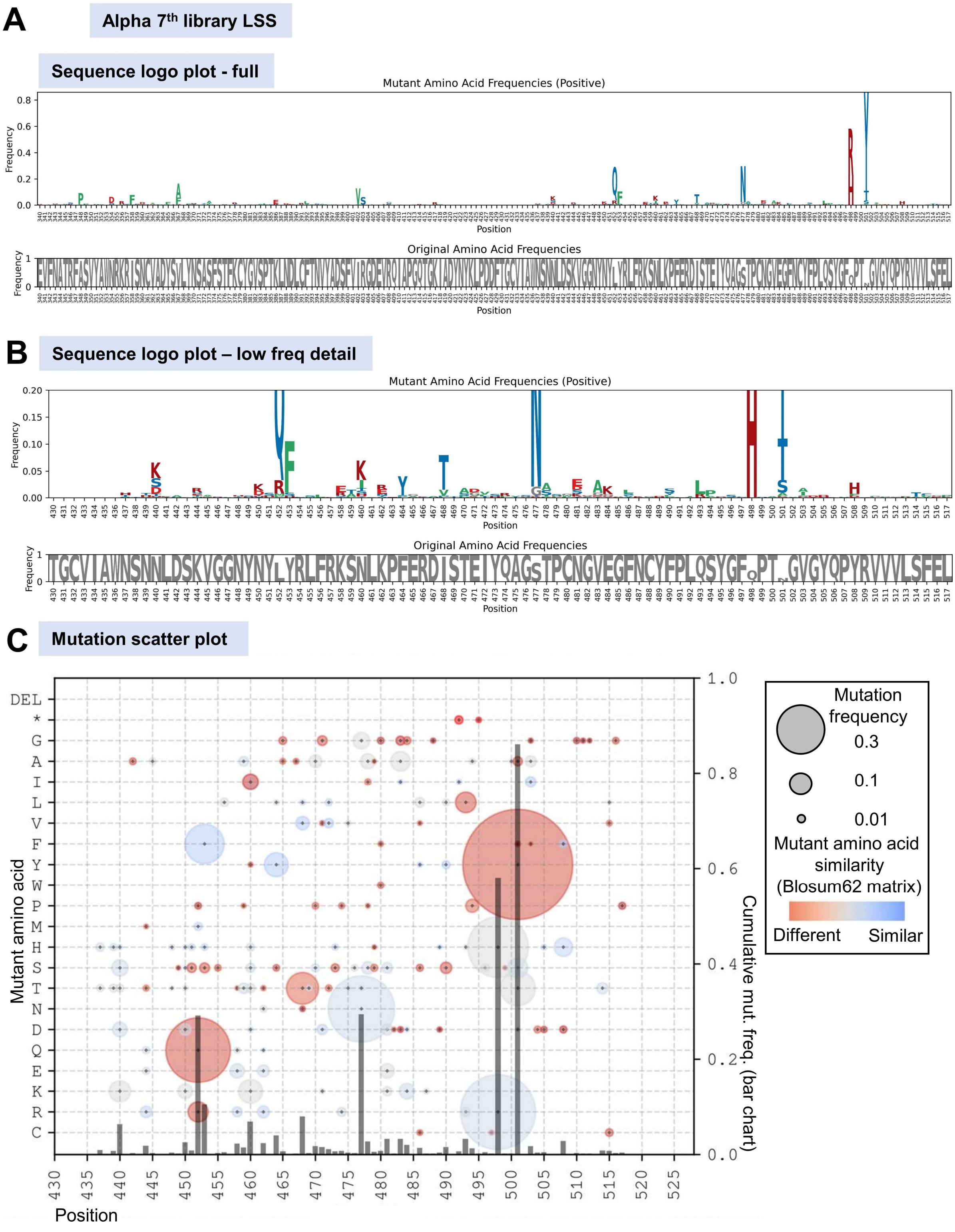

**Fig. S15 – Logo plots and mutation scatter plot for Alpha 7th library under LSS.** (A) Sequence logo plot (top panel) showing mutations in the library relative to the WT sequence. The bottom panel (in gray) displays the complementary frequency of the original amino acid at each position. (B) Sequence logo plot focusing on less frequent mutations in the library, with the y-axis frequency range set to 0–0.2. (C) Mutation scatter plot illustrating mutations in the population and their evolutionary distance from the original residues.

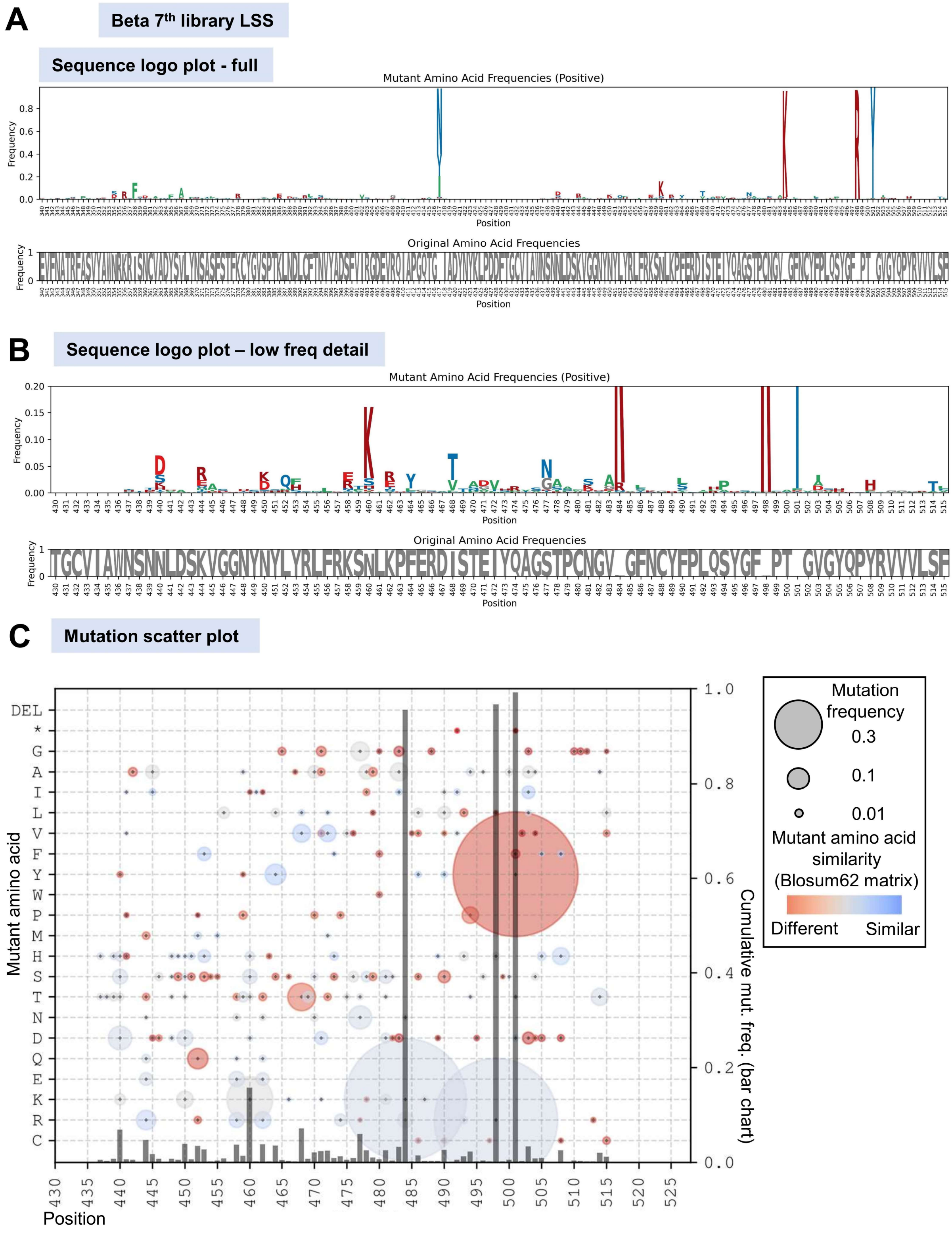

**Fig. S16 – Logo plots and mutation scatter plot for Beta 7th library under LSS.** (A) Sequence logo plot (top panel) showing mutations in the library relative to the WT sequence. The bottom panel (in gray) displays the complementary frequency of the original amino acid at each position. (B) Sequence logo plot focusing on less frequent mutations in the library, with the y-axis frequency range set to 0–0.2. (C) Mutation scatter plot illustrating mutations in the population and their evolutionary distance from the original residues.

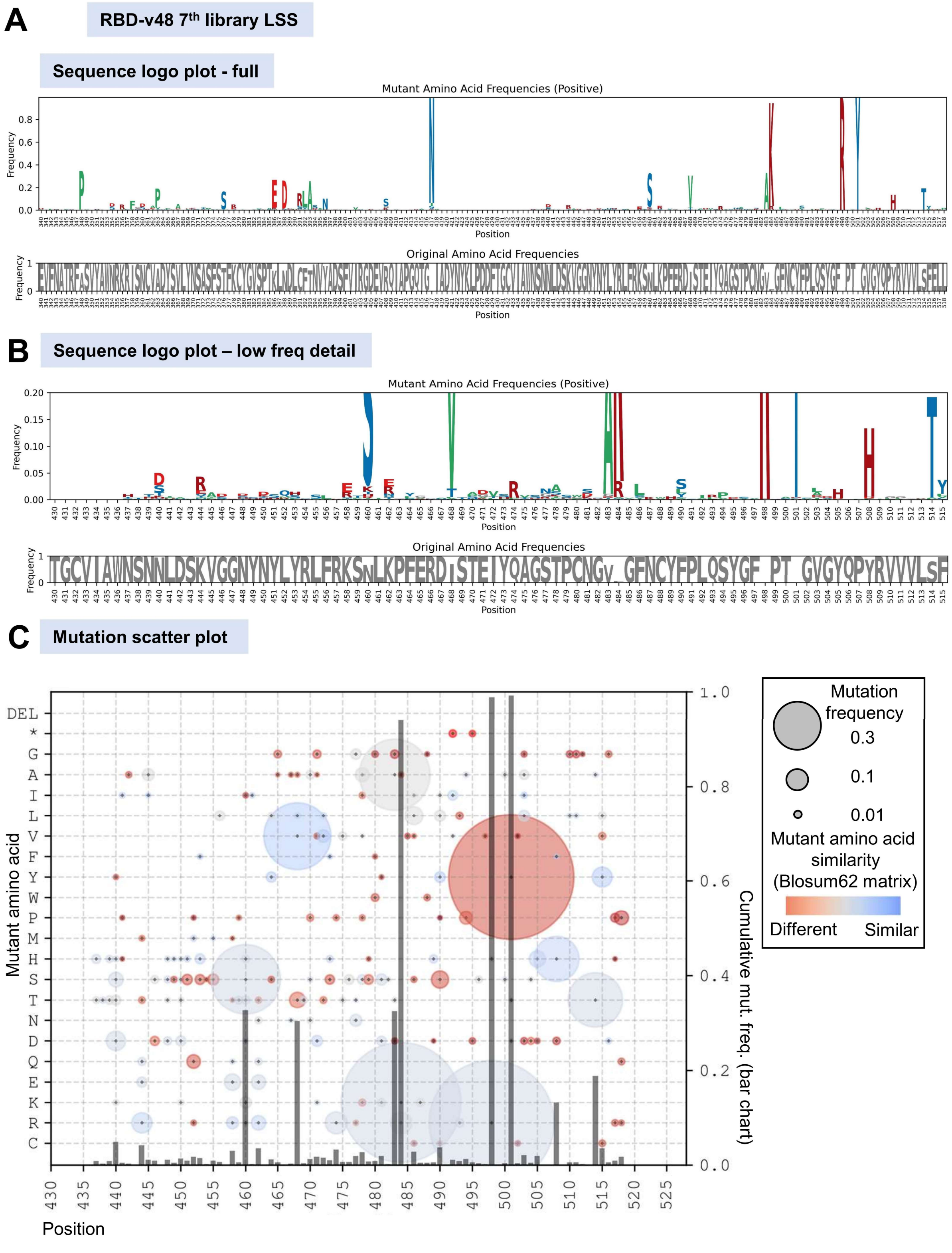

**Fig. S17 – Logo plots and mutation scatter plot for v48 7th library under LSS.** (A) Sequence logo plot (top panel) showing mutations in the library relative to the WT sequence. The bottom panel (in gray) displays the complementary frequency of the original amino acid at each position. (B) Sequence logo plot focusing on less frequent mutations in the library, with the y-axis frequency range set to 0–0.2. (C) Mutation scatter plot illustrating mutations in the population and their evolutionary distance from the original residues.

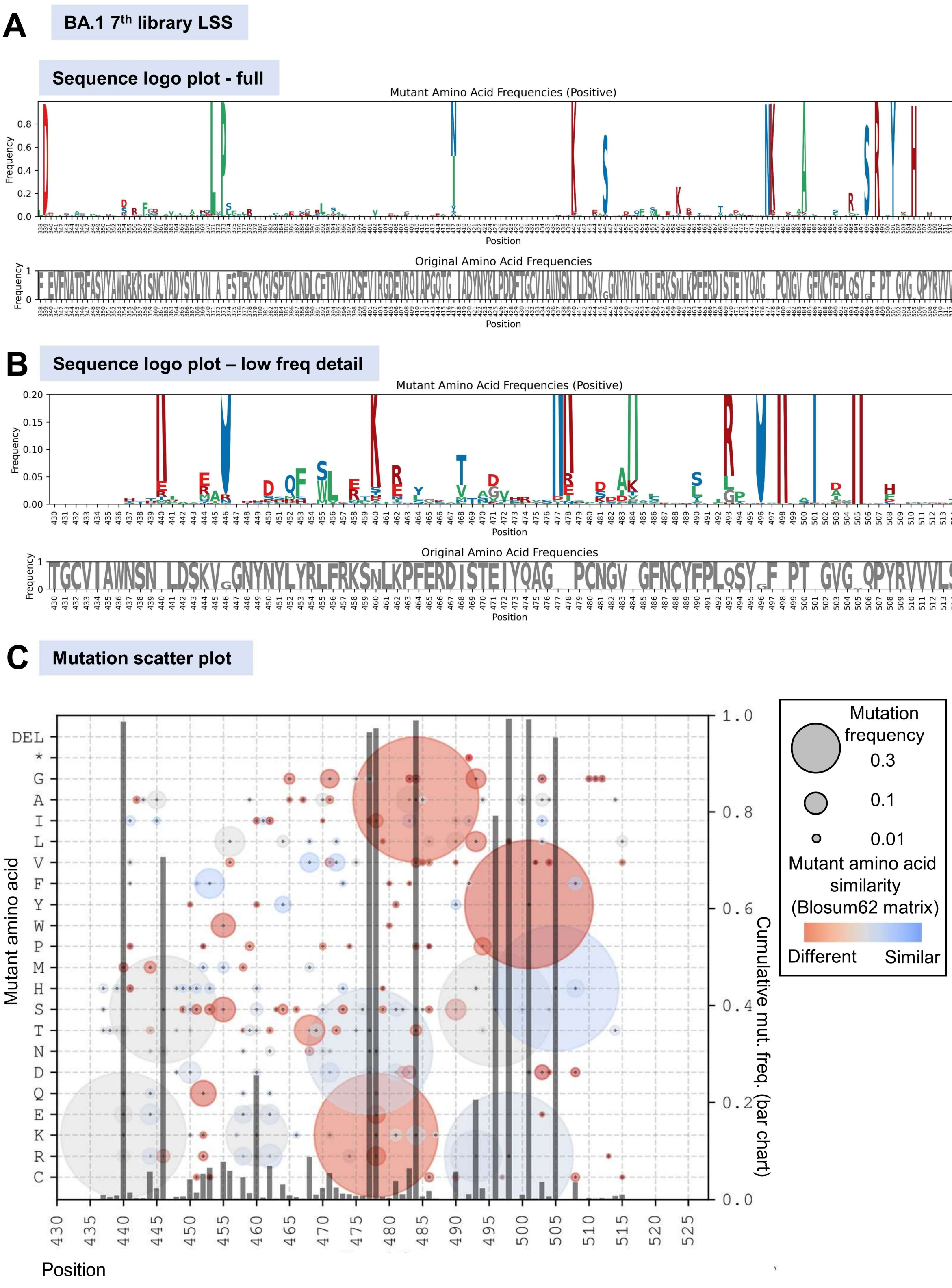

**Fig. S18 – Logo plots and mutation scatter plot for BA.1 7th library under LSS.** (A) Sequence logo plot (top panel) showing mutations in the library relative to the WT sequence. The bottom panel (in gray) displays the complementary frequency of the original amino acid at each position. (B) Sequence logo plot focusing on less frequent mutations in the library, with the y-axis frequency range set to 0–0.2. (C) Mutation scatter plot illustrating mutations in the population and their evolutionary distance from the original residues.

### Supporting information part PS4 – Analysis of mutations at different stages of high stringency selection libraries

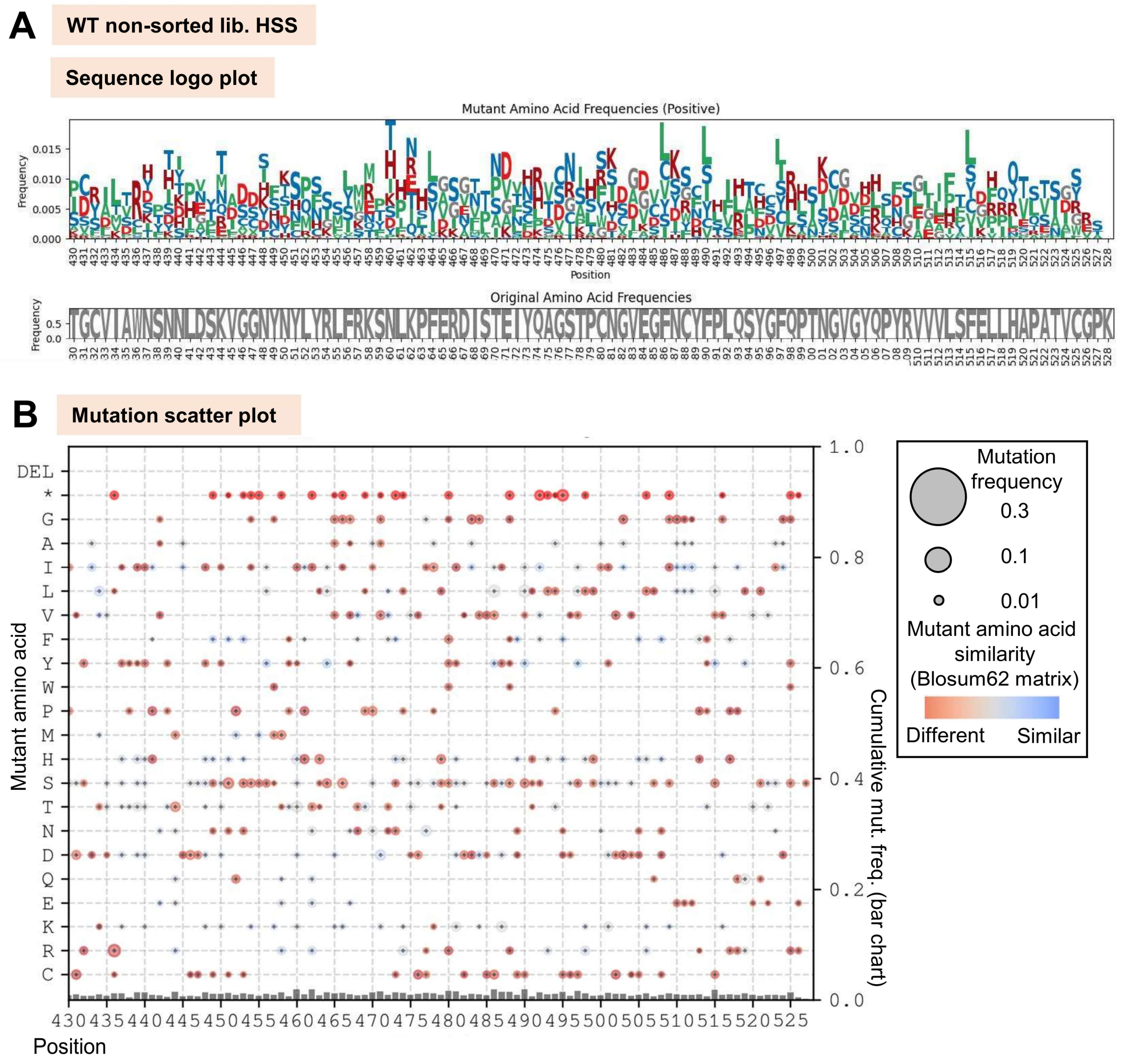

**Fig. S19 – Logo plot and mutation scatter plot for WT non-selected library for HSS.** (A) Sequence logo plot (top panel) showing mutations in the library relative to the WT sequence. The bottom panel (in gray) displays the complementary frequency of the original amino acid at each position. (B) Mutation scatter plot illustrating mutations in the population and their evolutionary distance from the original residues.

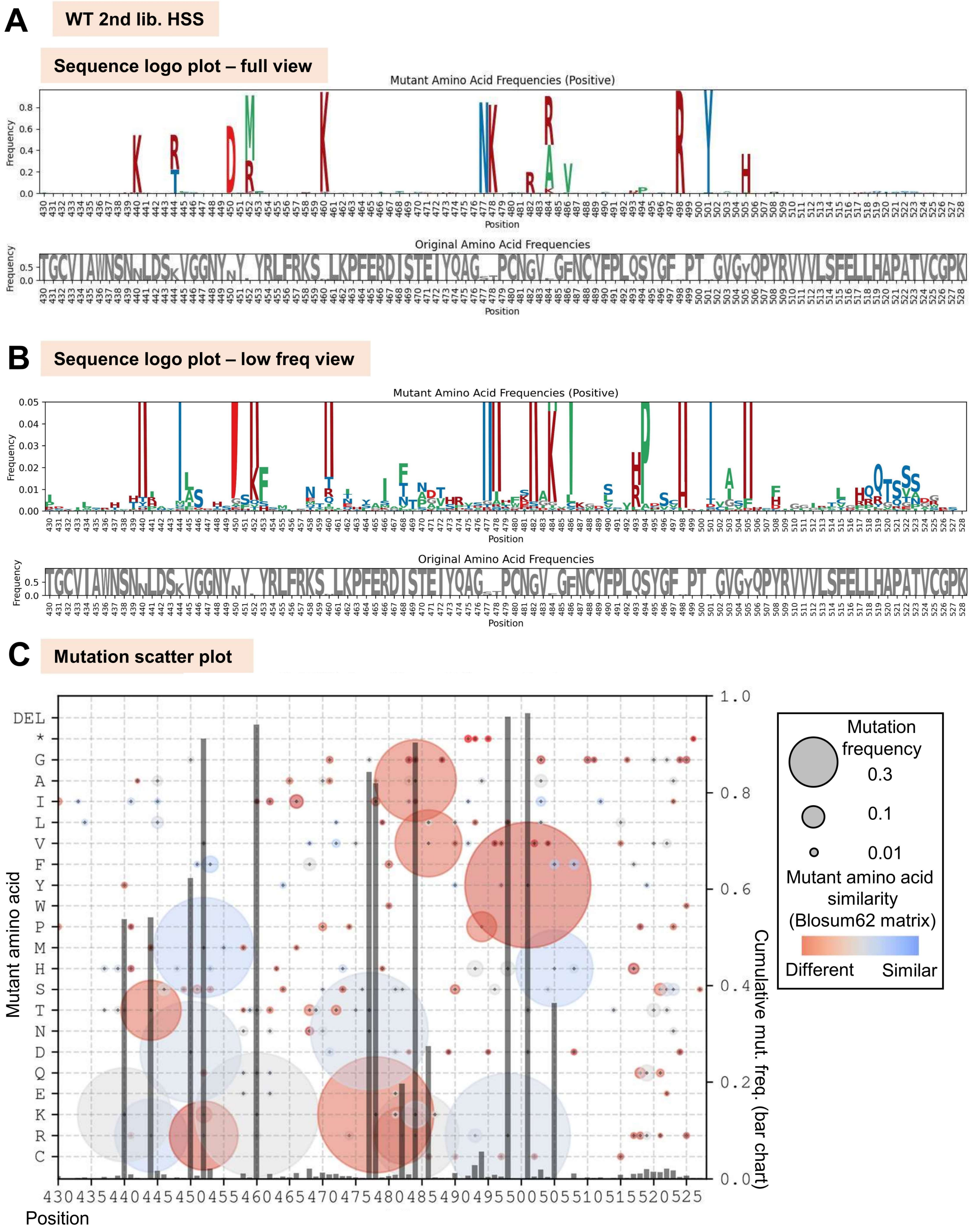

**Fig. S20 – Logo plots and mutation scatter plot for WT 2nd library selected under HSS.** (A) Sequence logo plot (top panel) showing mutations in the library relative to the WT sequence. The bottom panel (in gray) displays the complementary frequency of the original amino acid at each position. (B) Sequence logo plot focusing on less frequent mutations in the library, with the y-axis frequency range set to 0–0.2. (C) Mutation scatter plot illustrating mutations in the population and their evolutionary distance from the original residues.

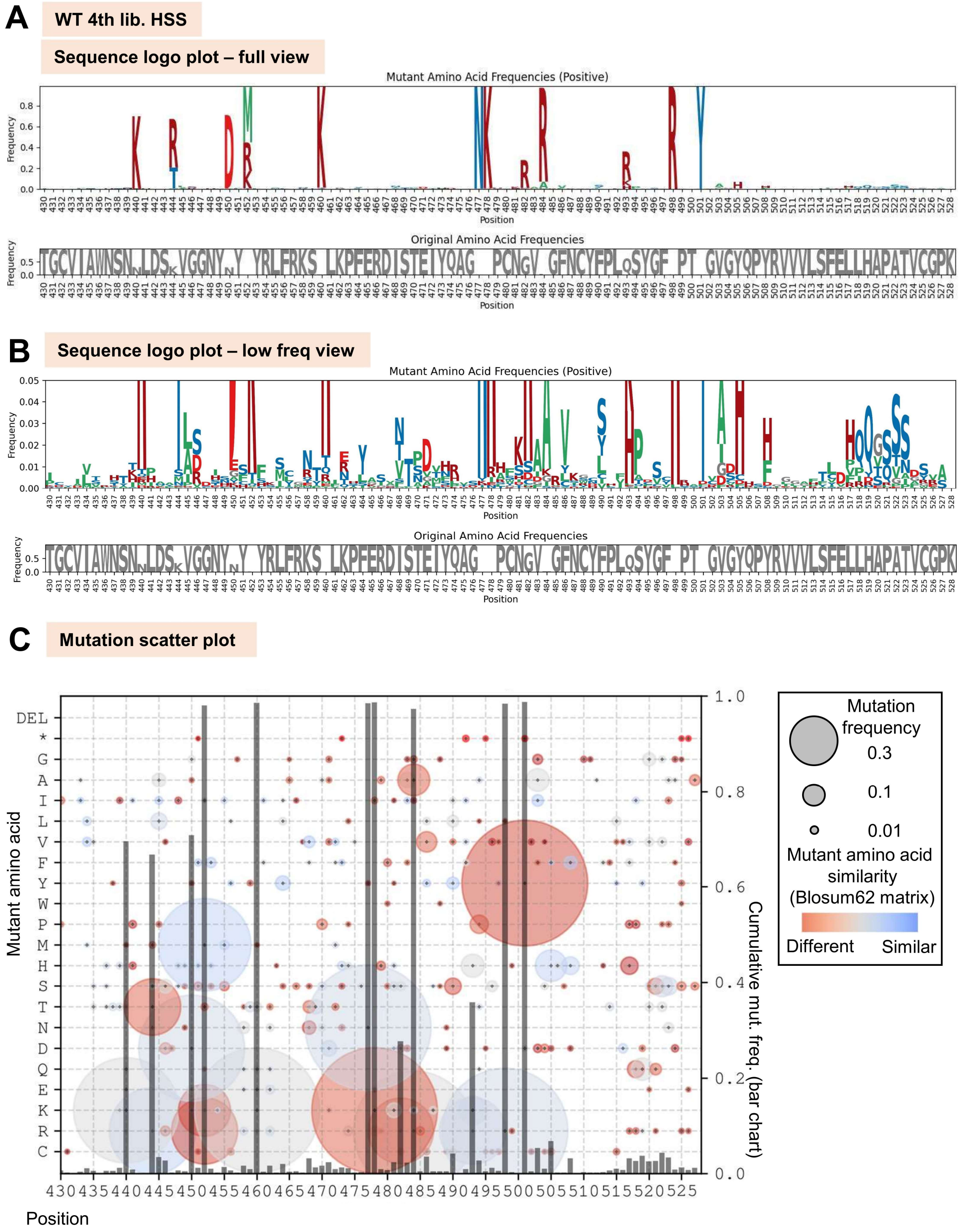

**Fig. S21 – Logo plots and mutation scatter plot for WT 4th library selected under HSS.** (A) Sequence logo plot (top panel) showing mutations in the library relative to the WT sequence. The bottom panel (in gray) displays the complementary frequency of the original amino acid at each position. (B) Sequence logo plot focusing on less frequent mutations in the library, with the y-axis frequency range set to 0–0.2. (C) Mutation scatter plot illustrating mutations in the population and their evolutionary distance from the original residues.

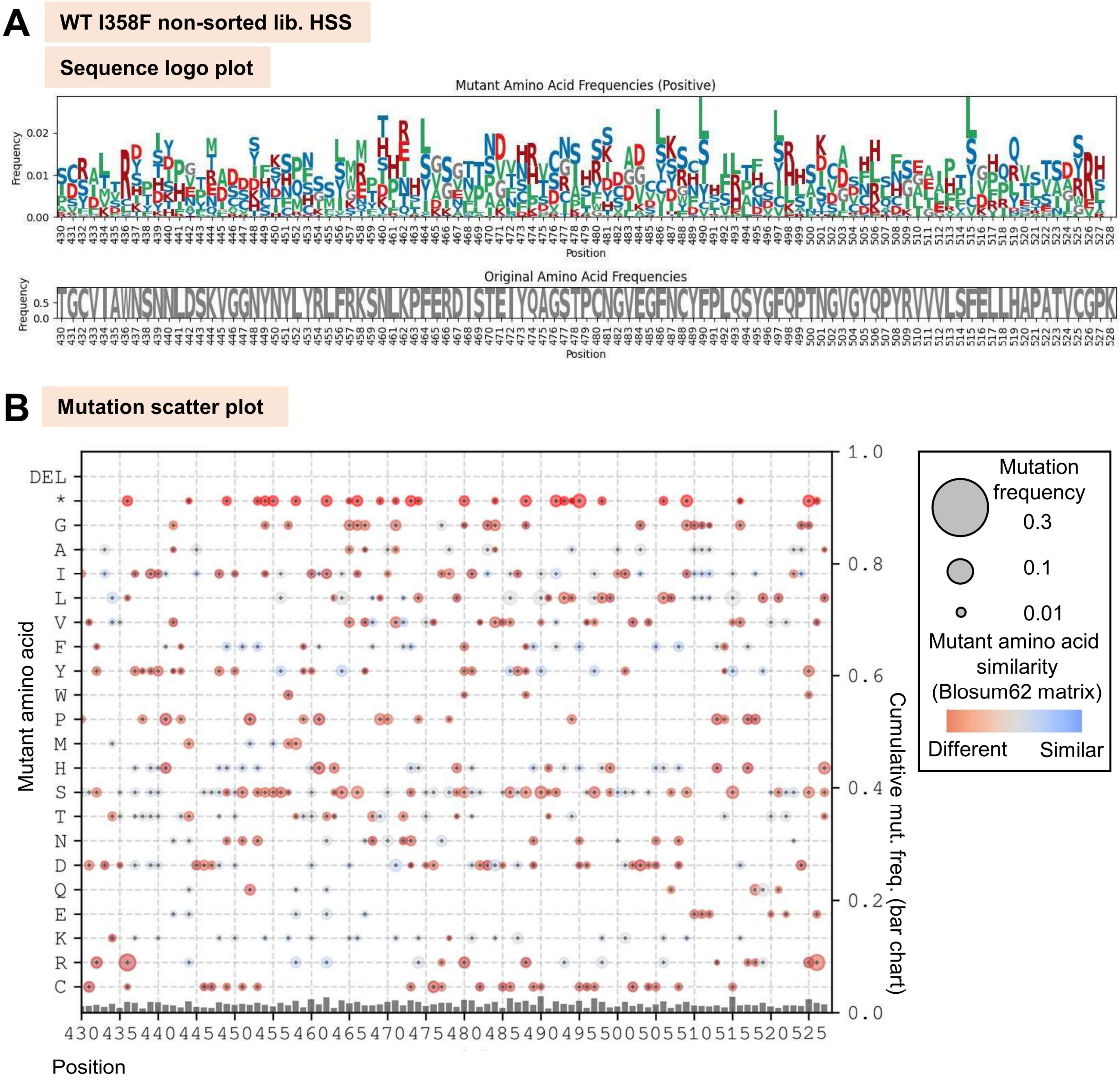

**Fig. S22– Logo plot and mutation scatter plot for WT I358F non-selected library for HSS.** (A) Sequence logo plot (top panel) showing mutations in the library relative to the WT sequence. The bottom panel (in gray) displays the complementary frequency of the original amino acid at each position. (B) Mutation scatter plot illustrating mutations in the population and their evolutionary distance from the original residues.

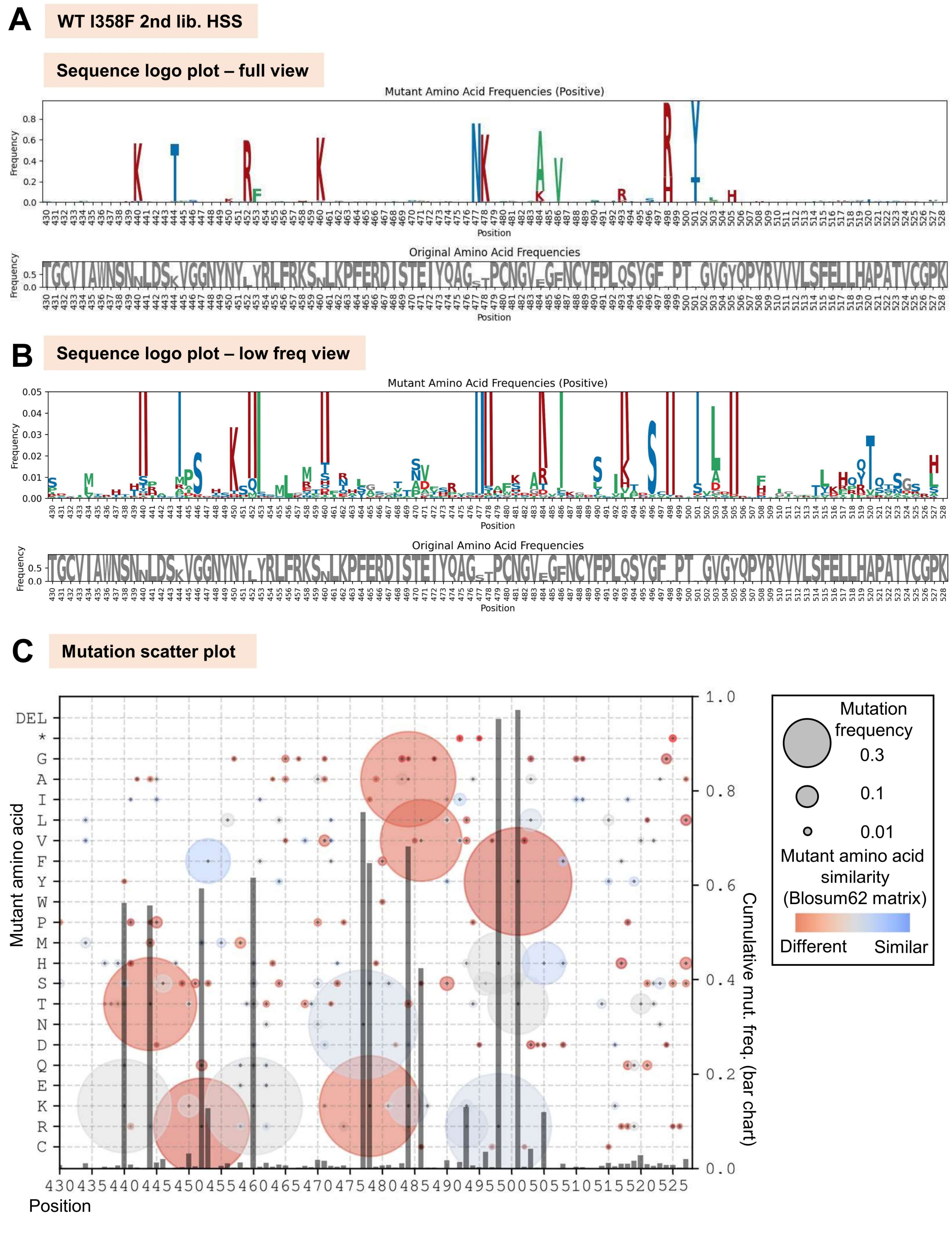

**Fig. S23 – Logo plots and mutation scatter plot for WT I358F 2nd library selected under HSS.** (A) Sequence logo plot (top panel) showing mutations in the library relative to the WT sequence. The bottom panel (in gray) displays the complementary frequency of the original amino acid at each position. (B) Sequence logo plot focusing on less frequent mutations in the library, with the y-axis frequency range set to 0–0.2. (C) Mutation scatter plot illustrating mutations in the population and their evolutionary distance from the original residues.

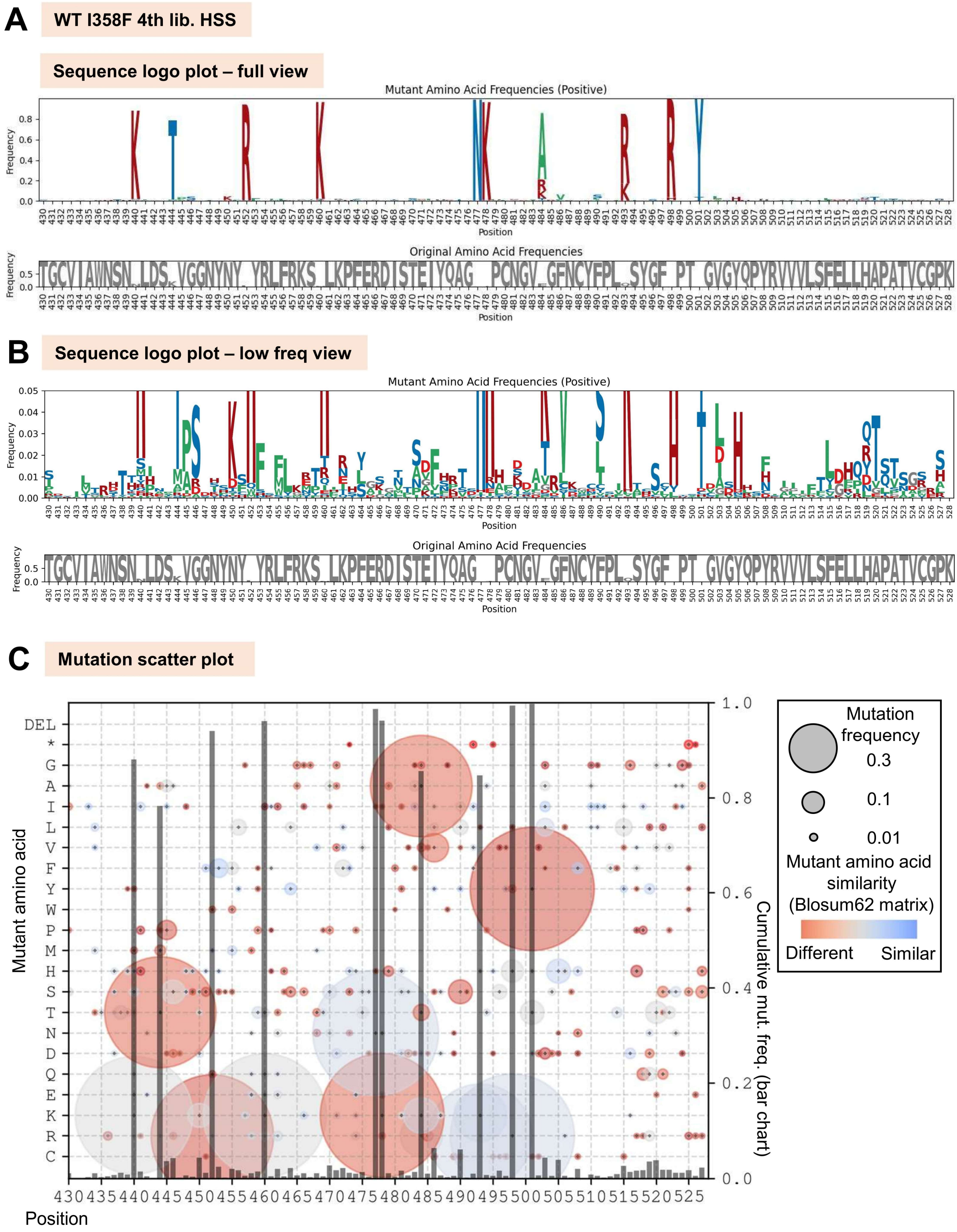

**Fig. S24 – Logo plots and mutation scatter plot for WT I358F 4th library selected under HSS.** (A) Sequence logo plot (top panel) showing mutations in the library relative to the WT sequence. The bottom panel (in gray) displays the complementary frequency of the original amino acid at each position. (B) Sequence logo plot focusing on less frequent mutations in the library, with the y-axis frequency range set to 0–0.2. (C) Mutation scatter plot illustrating mutations in the population and their evolutionary distance from the original residues.

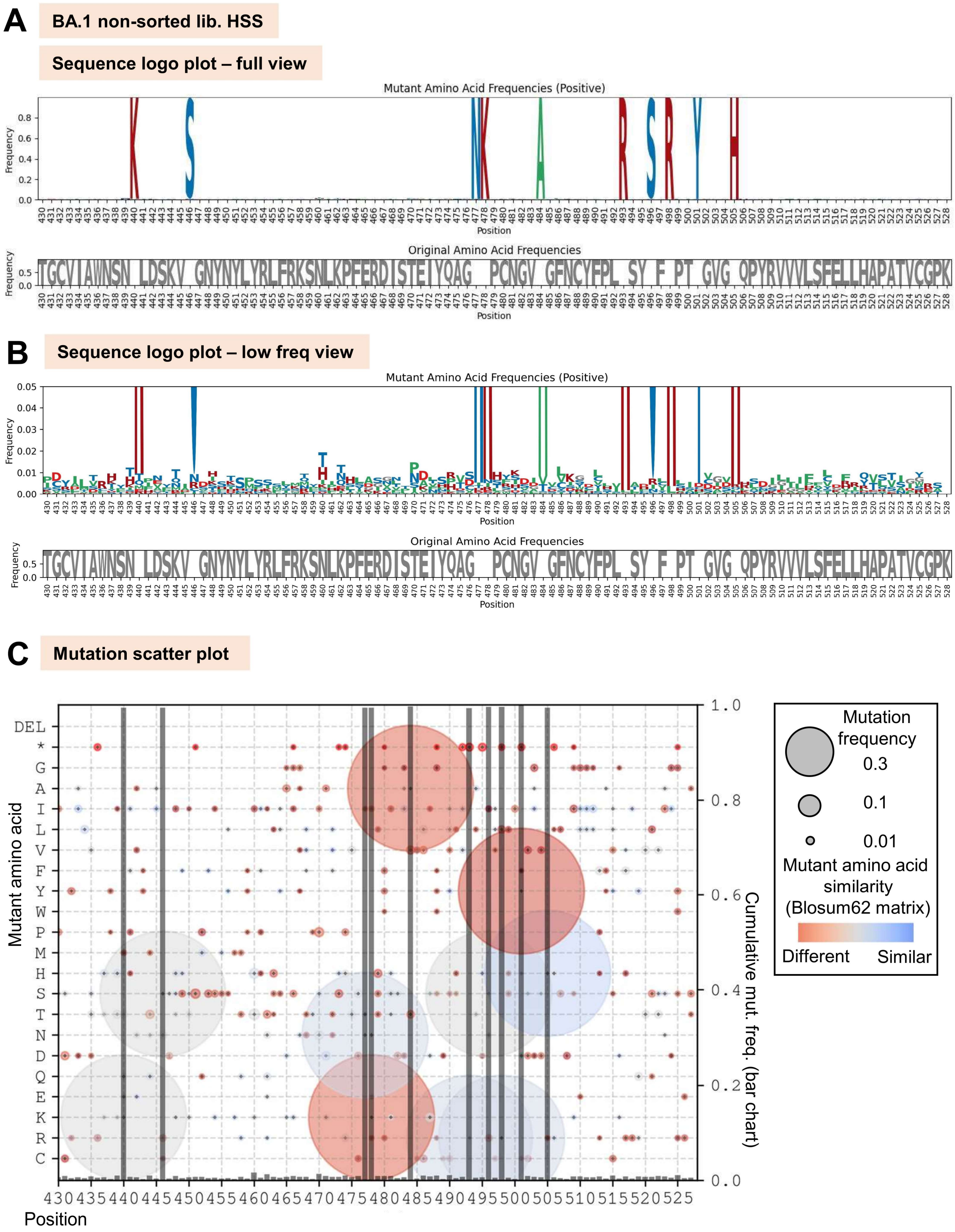

**Fig. S25 – Logo plots and mutation scatter plot for BA.1 non-selected library selected under HSS.** (A) Sequence logo plot (top panel) showing mutations in the library relative to the WT sequence. The bottom panel (in gray) displays the complementary frequency of the original amino acid at each position. (B) Sequence logo plot focusing on less frequent mutations in the library, with the y-axis frequency range set to 0–0.2. (C) Mutation scatter plot illustrating mutations in the population and their evolutionary distance from the original residues.

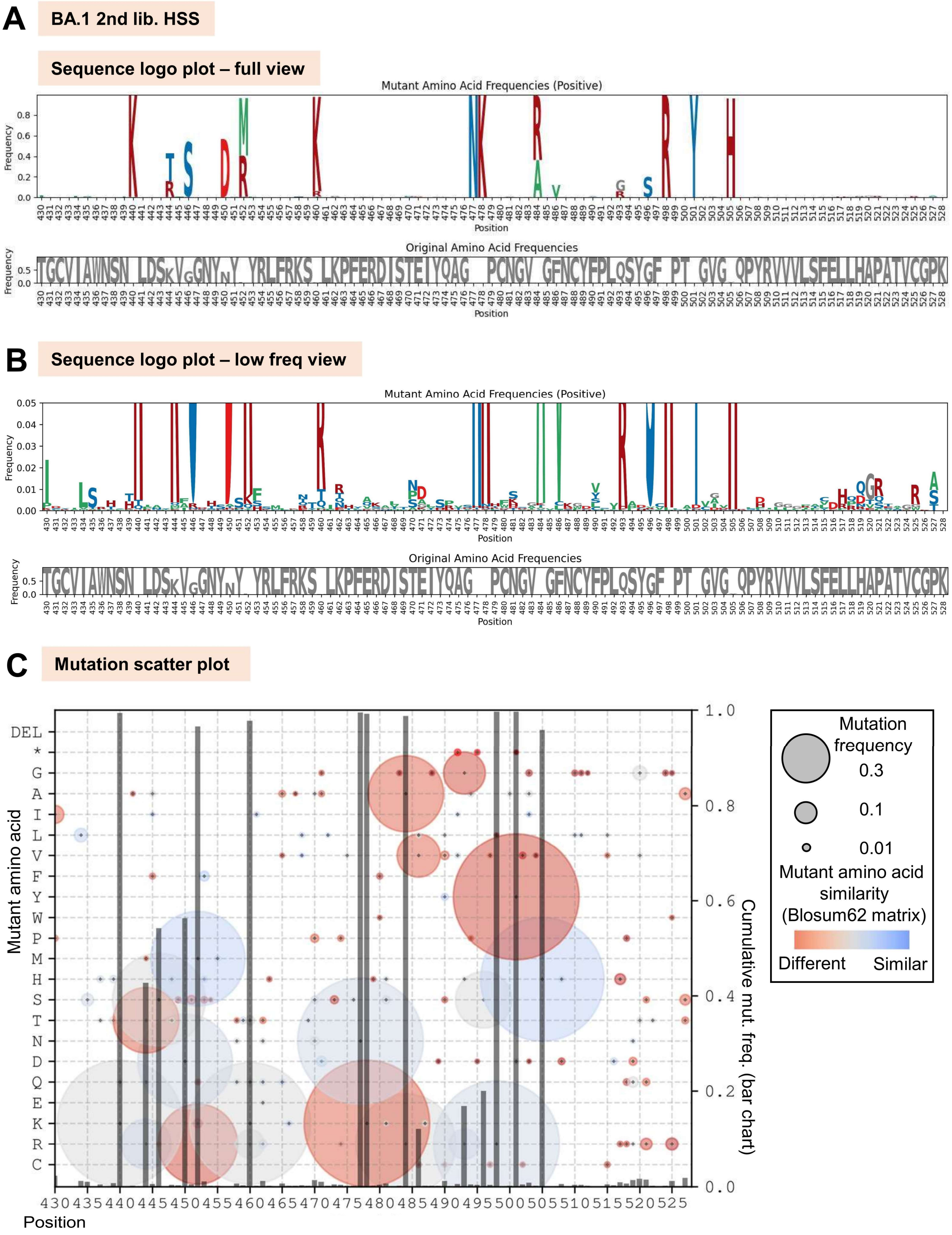

**Fig. S26 – Logo plots and mutation scatter plot for BA.1 2nd library selected under HSS.** (A) Sequence logo plot (top panel) showing mutations in the library relative to the WT sequence. The bottom panel (in gray) displays the complementary frequency of the original amino acid at each position. (B) Sequence logo plot focusing on less frequent mutations in the library, with the y-axis frequency range set to 0–0.2. (C) Mutation scatter plot illustrating mutations in the population and their evolutionary distance from the original residues.

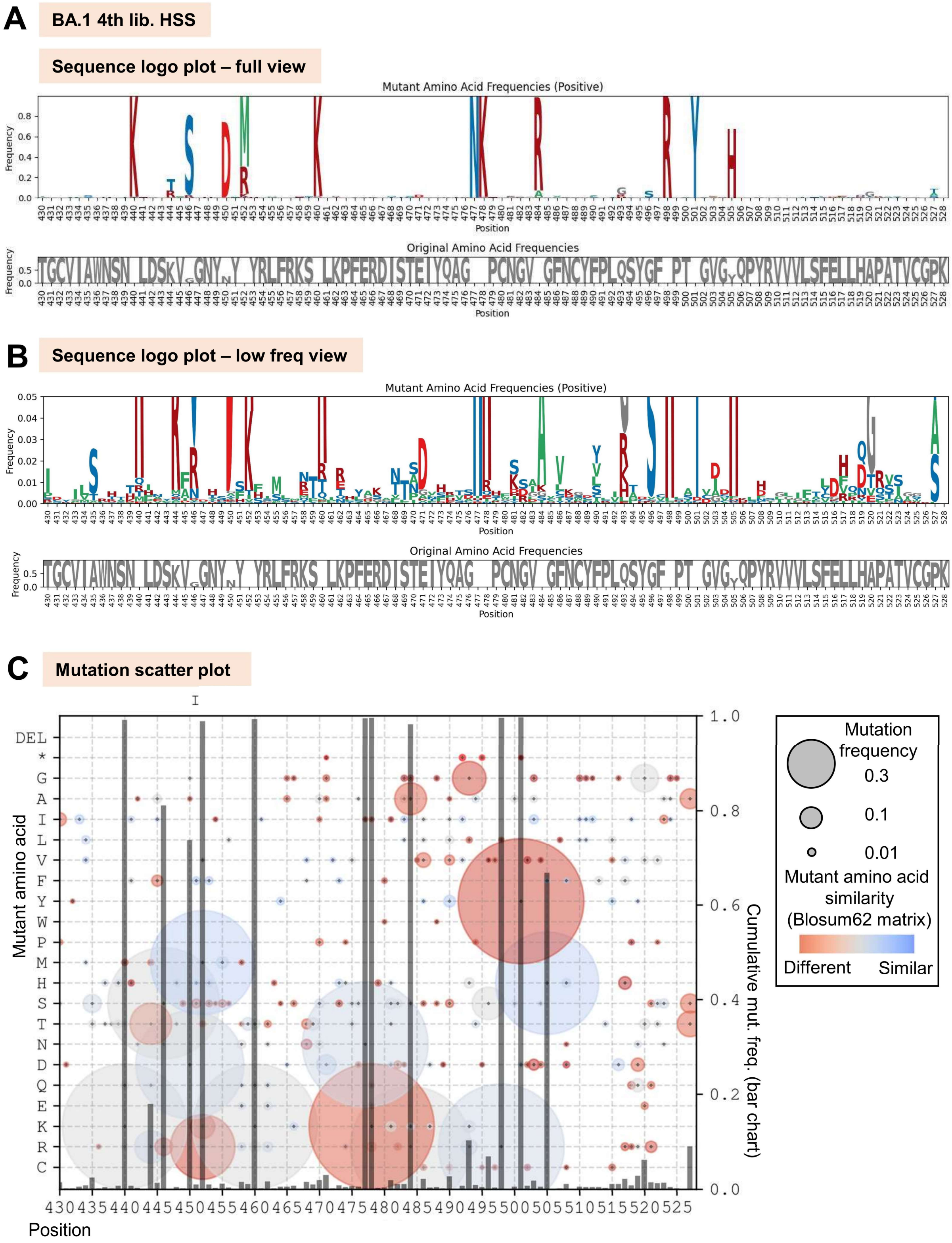

**Fig. S27 – Logo plots and mutation scatter plot for BA.1 4th library selected under HSS.** (A) Sequence logo plot (top panel) showing mutations in the library relative to the WT sequence. The bottom panel (in gray) displays the complementary frequency of the original amino acid at each position. (B) Sequence logo plot focusing on less frequent mutations in the library, with the y-axis frequency range set to 0–0.2. (C) Mutation scatter plot illustrating mutations in the population and their evolutionary distance from the original residues.

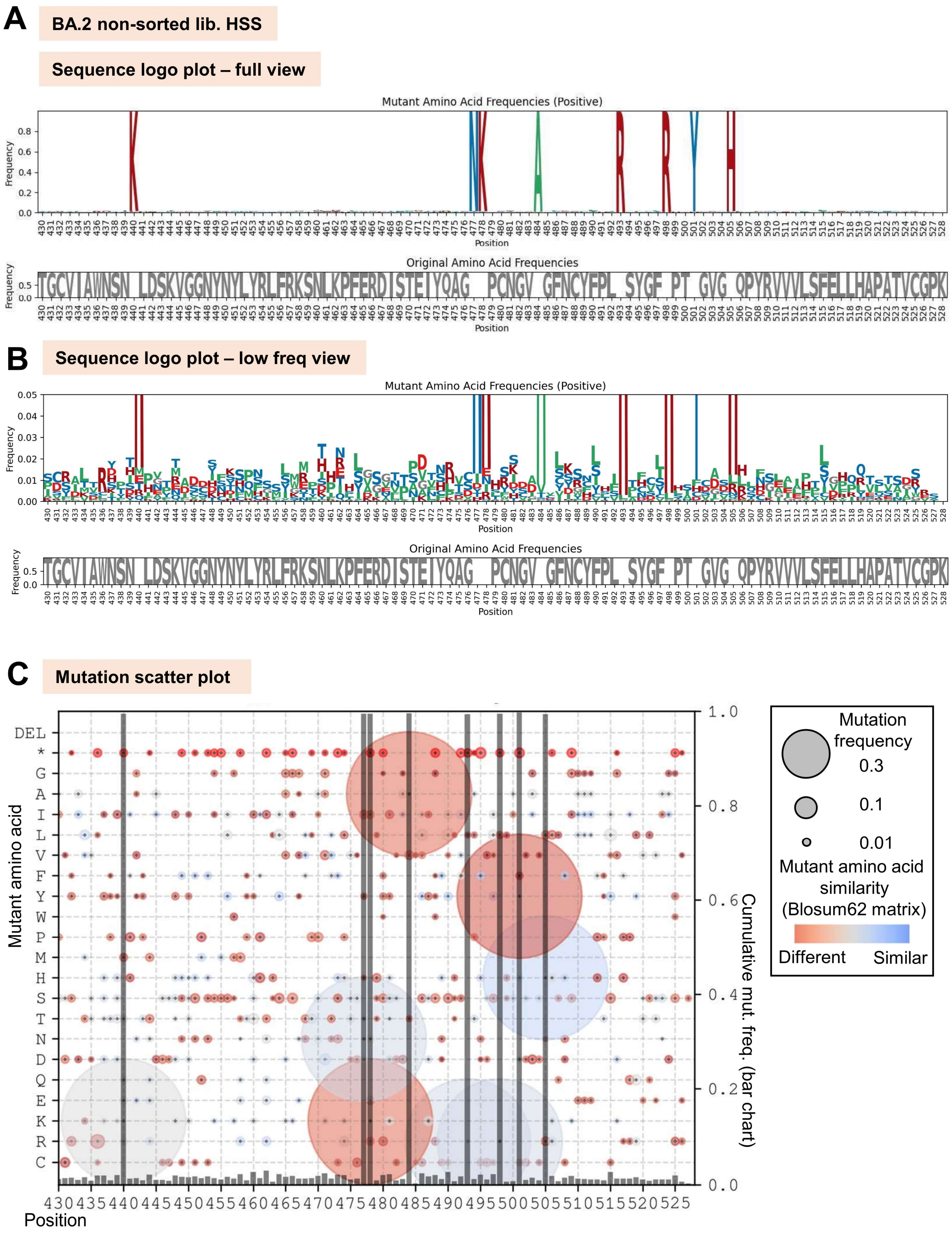

**Fig. S28 – Logo plots and mutation scatter plot for BA.2 non-selected library for HSS.** (A) Sequence logo plot (top panel) showing mutations in the library relative to the WT sequence. The bottom panel (in gray) displays the complementary frequency of the original amino acid at each position. (B) Sequence logo plot focusing on less frequent mutations in the library, with the y-axis frequency range set to 0–0.2. (C) Mutation scatter plot illustrating mutations in the population and their evolutionary distance from the original residues.

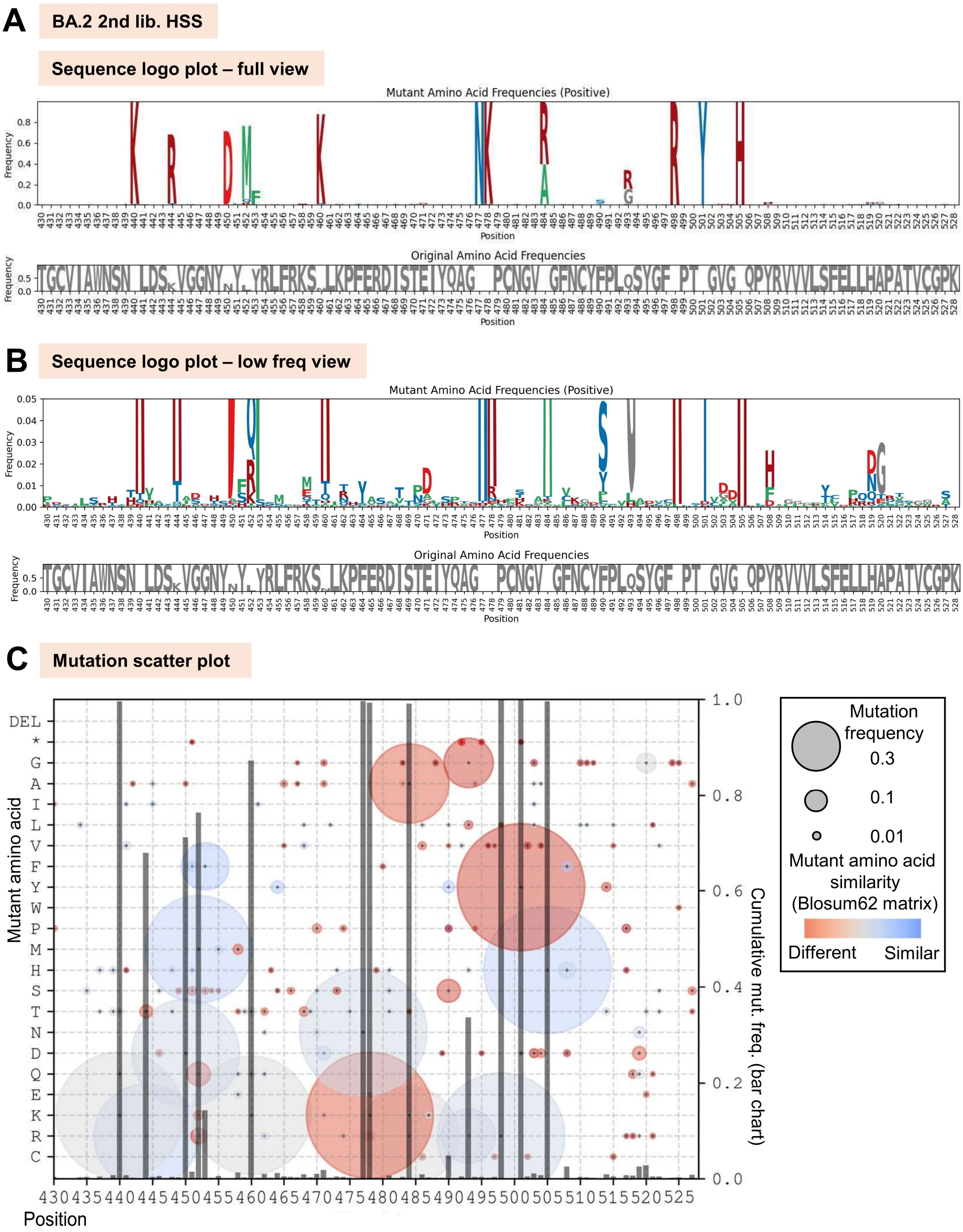

**Fig. S29 – Logo plots and mutation scatter plot for BA.2 2nd library selected under HSS.** (A) Sequence logo plot (top panel) showing mutations in the library relative to the WT sequence. The bottom panel (in gray) displays the complementary frequency of the original amino acid at each position. (B) Sequence logo plot focusing on less frequent mutations in the library, with the y-axis frequency range set to 0–0.2. (C) Mutation scatter plot illustrating mutations in the population and their evolutionary distance from the original residues.

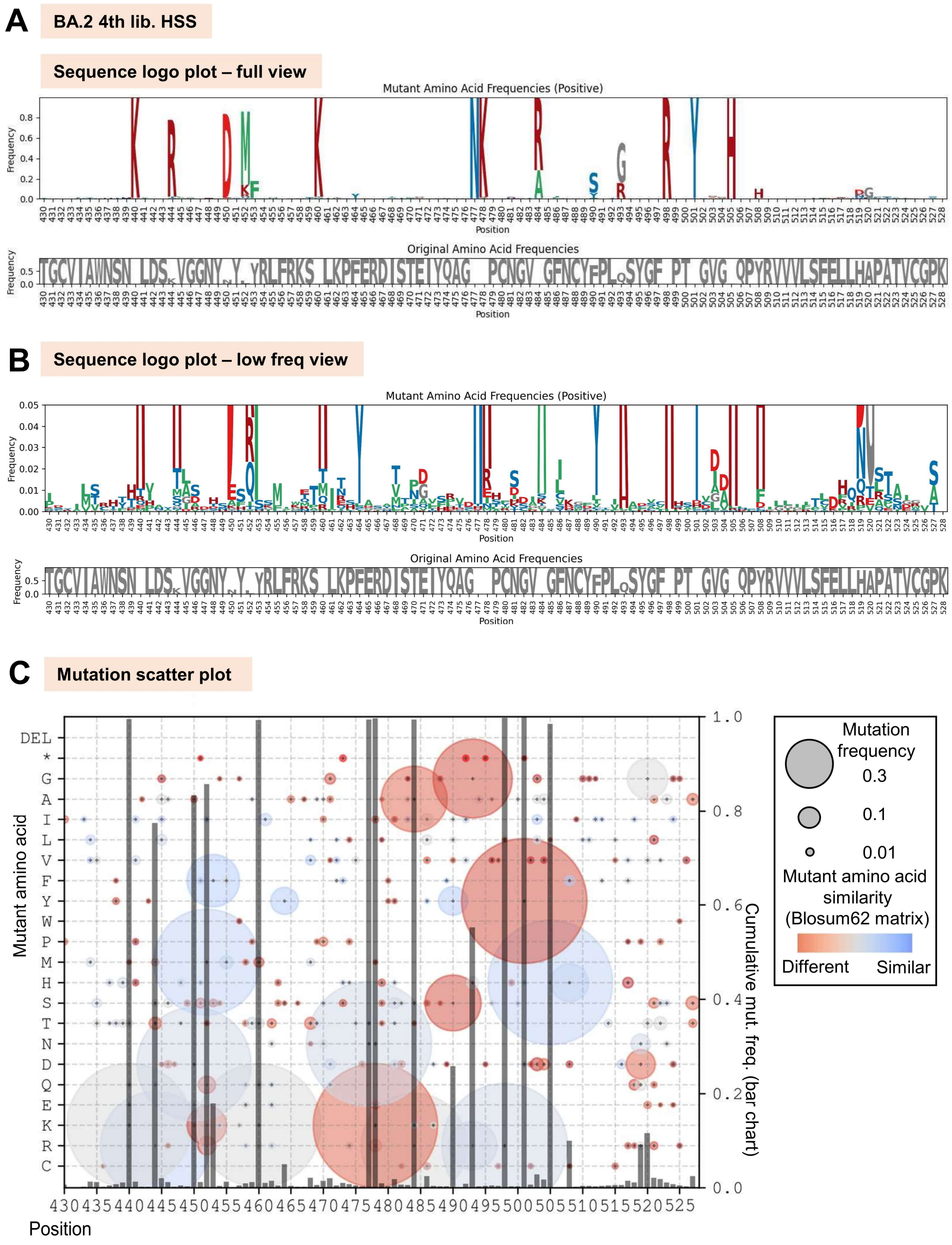

**Fig. S30 – Logo plots and mutation scatter plot for BA.2 4th library selected under HSS.** (A) Sequence logo plot (top panel) showing mutations in the library relative to the WT sequence. The bottom panel (in gray) displays the complementary frequency of the original amino acid at each position. (B) Sequence logo plot focusing on less frequent mutations in the library, with the y-axis frequency range set to 0–0.2. (C) Mutation scatter plot illustrating mutations in the population and their evolutionary distance from the original residues.

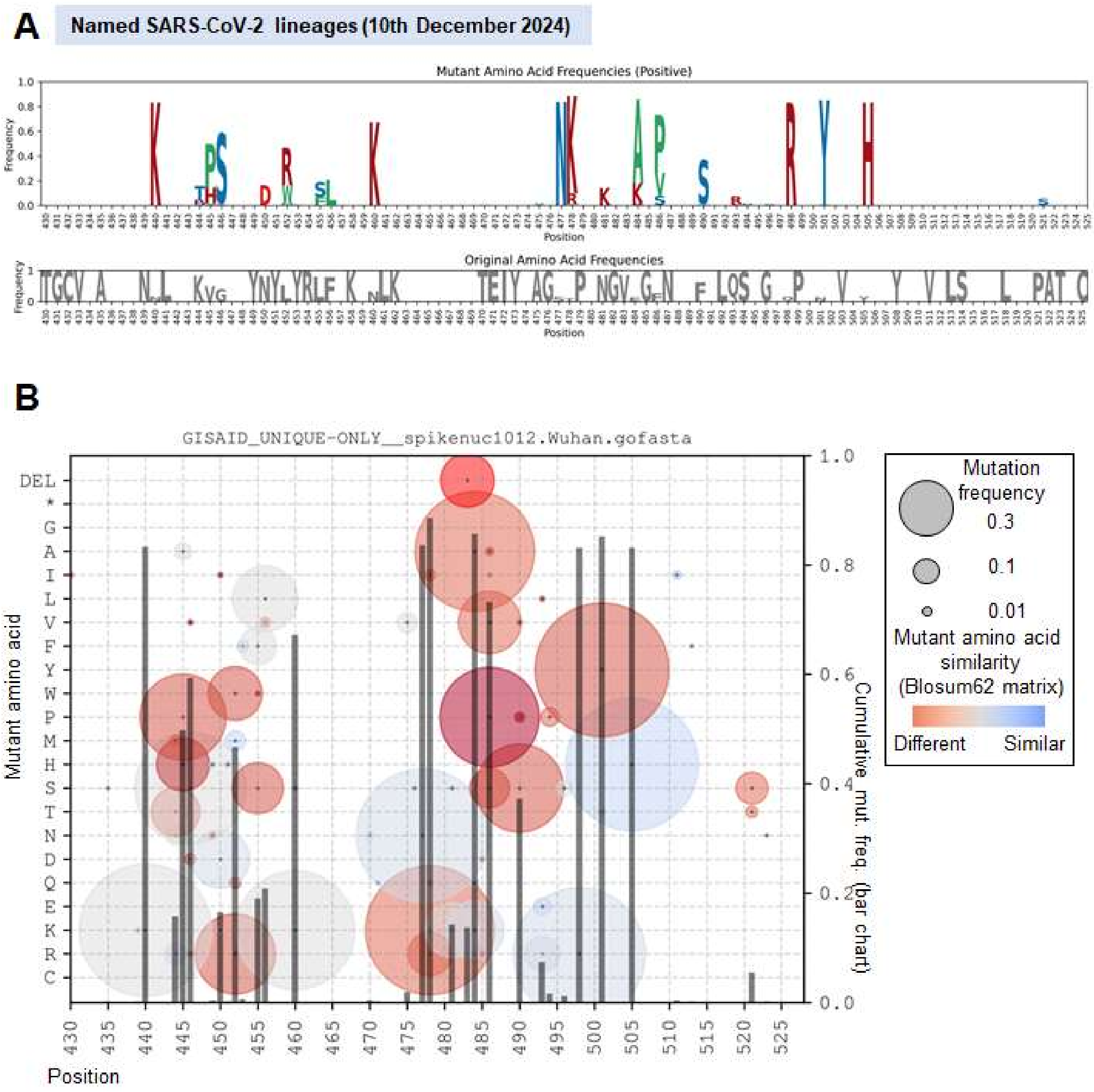

**Fig. S31 – Logo plots and mutation scatter plot for the complete set of Pango named lineages** curated on the 10th December 2024 and downloaded from https://github.com/corneliusroemer/pango-sequences.

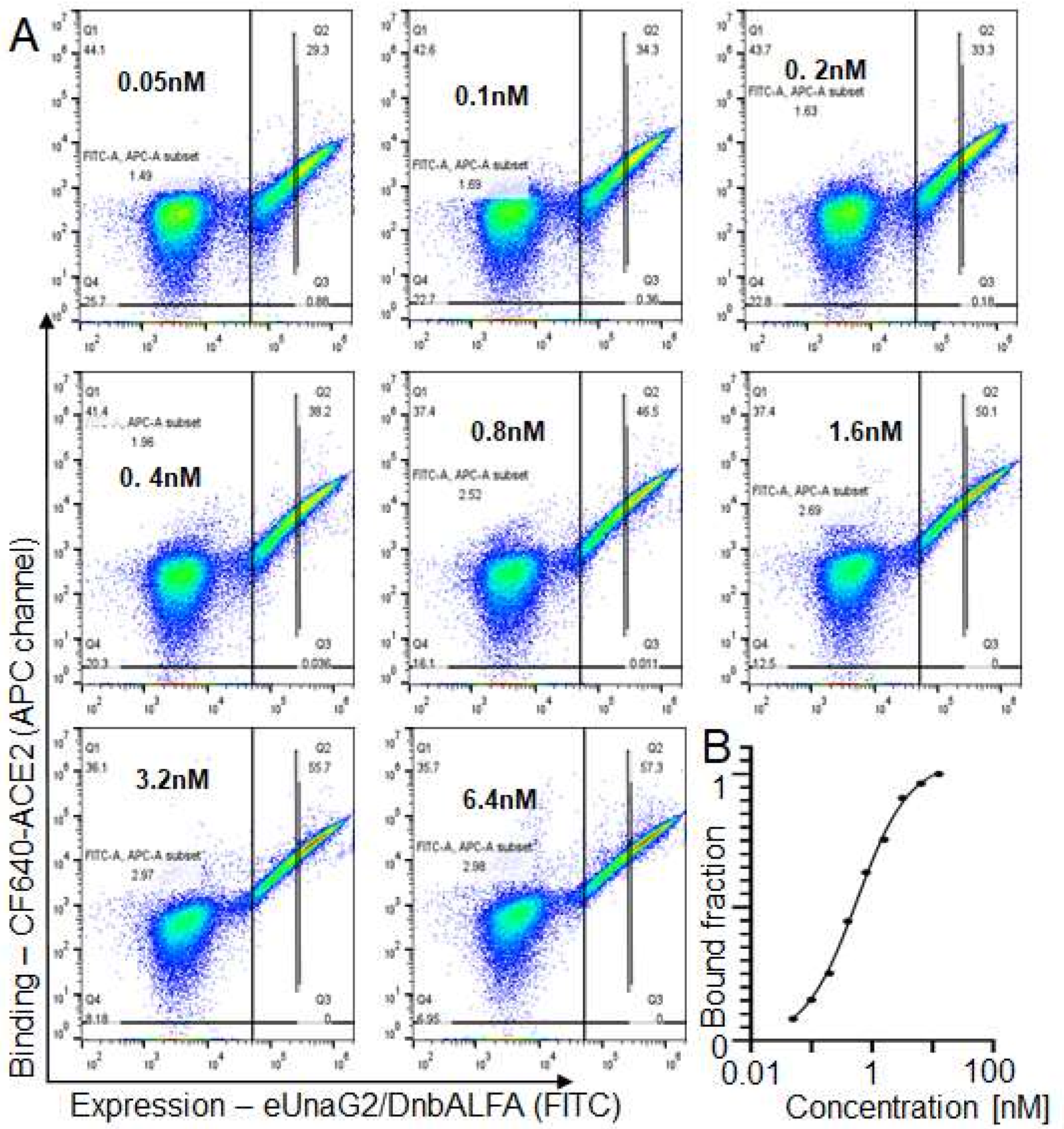

**Fig. S32 – Gating strategy for yeast display-based binding affinity measurements**. (**A**) A narrow vertical subset/subpopulation was defined to normalize the expression rates across samples. Specific binding (APC-A signal intensity) for each measurement was determined by subtracting the negative binding (Q1 median APC-A intensity) from the positive binding (Q2>subset median APC-A intensity). (**B**) Specific binding values at each ACE-2 concentration were fitted using GraphPad (Methods).
